# Pharmacoproteomic profiling identifies secreted markers for aberrant drug action

**DOI:** 10.1101/2024.10.16.618637

**Authors:** Sascha Knecht, Mathias Kalxdorf, Johanna Korbeń, Toby Mathieson, Daniel C. Sevin, Bernhard Kuster, Richard Kasprowicz, Melanie Z. Sakatis, H. Christian Eberl, Marcus Bantscheff

## Abstract

Adverse drug reactions (ADRs) contribute significantly to late-stage attrition in drug discovery due to their unpredictability and enigmatic underlying mechanisms. Here we applied mass spectrometry-based proteomics to assess the effects of 46 approved or retracted drugs with various levels of concerns for drug-induced liver injury and annotated for mitochondrial mechanisms, along with 8 tool compounds, on the secretome of a hepatocyte liver model. We observed distinct clusters of non-canonical secretion, and intracellular thermal proteome profiling linked dysregulated mechanisms to extracellular markers. Functional follow-up confirmed lysosomal alterations by cationic-amphiphilic drugs, connected damage of the respiratory chain to Rab7-dependent secretion of mitochondrial proteins, and linked drug-induced endoplasmic reticulum stress to reduced basal secretion. Perturbation of sphingolipid biosynthesis pathways specifically induced secretion of the cargo sorting protein SDF4 whilst suppressing secretion of its cargo proteins. Thermal stability changes of clusters of membrane proteins in distinct subcellular compartments suggest local accumulation as important driver for unexpected drug effects through direct and indirect interactions.

## Introduction

Drug efficacy and safety are two critical aspects of drug development. To understand molecular mechanisms driving efficacy and adverse drug reactions (ADRs) systems-wide omics approaches are gaining significance. Omics profiling enables the identification of affected pathways and provides clues for causal events leading to defined desirable or undesirable endpoints. Large-scale proteomics studies have been conducted to investigate the effects of small-molecule drugs on cell lines, leading to an improved understanding of compound mechanism of action (MoA) and ADRs ^1, 2^. However, most studies have focused on intracellular effects providing limited insights into biomarkers that would enable monitoring of pre-clinical animal models or patients in a clinical setting. Secreted proteins can be detected in both model systems and serum samples routinely collected during preclinical and clinical studies, and thus can potentially be used as translational markers. However, systematic information to which extent intracellular mechanisms are reflected in the extracellular proteome is limited.

The liver has the highest susceptibility to adverse drug reactions because of its important role in the biotransformation of xenobiotics. Owing to the first-pass effect, orally administered drugs can transiently reach high concentrations in the liver, potentially leading to drug-mediated toxicity in this organ. The term drug-induced liver injury (DILI) summarizes unexpected adverse effects that drugs cause to the liver by damaging hepatocytes and other liver-resident cell types ^3–5^. DILI is a leading cause for attrition of drug candidates ^3, 4^ and has been observed for a variety of structurally diverse drugs, leading to a wide spectrum of clinical manifestations. To date, the FDA has annotated 750 approved drugs to cause DILI with classifications related to their respective risk potentials ^3, 6^. A substantial proportion of these drugs were withdrawn from the market due to unpredictable hepatotoxicity despite having met preclinical and clinical testing expectations and receiving regulatory approval. This illustrates the need for markers and procedures that can identify and predict safety concerns and toxicities during the preclinical stage.

Here, we studied the effects of 46 DILI drugs and 8 tool compounds on the secretome of the HepaRG hepatocyte model system in the differentiated state (dHepaRG) which maintains key hepatic functions including drug transporters and xenobiotic-metabolizing enzymes ^7–13^. We found marked differences in the secretomes of drugs with similar efficacy mechanisms in line with different degrees of DILI concern. We demonstrated by thermal proteome profiling and other functional studies that protein secretion patterns can be related to distinct intracellular mechanisms dysregulated by these compounds. Taken together, our data provide insights into a surprisingly diverse range of mechanisms that are dysregulated by DILI drugs and the manifestation of these mechanisms in the secretome of liver cell models, suggesting novel marker proteins for monitoring such unwanted effects in preclinical studies and in patients.

## Results

### DILI compounds induce secretion of proteins from specific cellular organelles

For secretome profiling in the dHepaRG hepatocyte model system, we selected 46 drugs with DILI concerns covering a broad chemical space and eight tool compounds to probe a range of cellular functions (Fig. 1a, Supplementary Table 1, Supplementary Fig. 1a). Cells were incubated with DILI compounds in triplicates at 2 and 20 µM, and supernatants were collected after 8 h of drug treatment within a 2 h collection window after switching to serum-free medium (Fig. 1a). The switch to serum-free medium was necessary to increase the dynamic range of proteomics experiments ^14^ , and we kept the serum-free windows as short as practical to minimize the negative impact of serum withdrawal and ensure cellular integrity ^15^. DILI compound concentrations used reflected the median maximum blood concentration (Cmax) of the panel and 10-fold this concentration (Supplementary Table 1). These concentrations are below the 20x to 200x Cmax commonly used for in vitro organ toxicity predictions ^7, 16, 17^ and thus closer to clinically relevant concentrations. Tool compounds were tested at the indicated concentrations to obtain selectivity (Fig. 1b, Supplementary Table 2). For a subset of the DILI and tool compounds additional secretomics analyses were performed after 2 h of drug treatments to distinguish early from late secretion events (Supplementary Table 2). LDH activity and lactic acid levels were determined in every supernatant sample to assess cell health and plasma membrane integrity. The maximal LDH activity was determined by lysing the control cells (equals 100 % cell death) and the percentage of cell death upon compound was determined as percentage to the maximal LDH activity. Only samples with low LDH activity (≤ 5% cell death compared to control samples), indicative for no or only low cell leakage, were used for proteomic analysis. Afatinib, chloroquine, cromoglicic acid, haloperidol, nefazodon, stavudine and tamoxifen induced high extracellular LDH, indicative of cell leakage at the highest tested concentration, and therefore only samples from the lower tested concentrations were analyzed. In contrast, compounds such as amiodarone, perhexiline, troglitazone led to increased extracellular lactic acid levels (Supplementary Fig. 1b-f), indicative of respiratory chain inhibition and, according to the FDA DILIrank dataset ^6^, consistent with the annotation “most-DILI concern” which refers to drugs that have the highest potential to cause DILI and severe clinical outcomes.

**Fig. 1.**
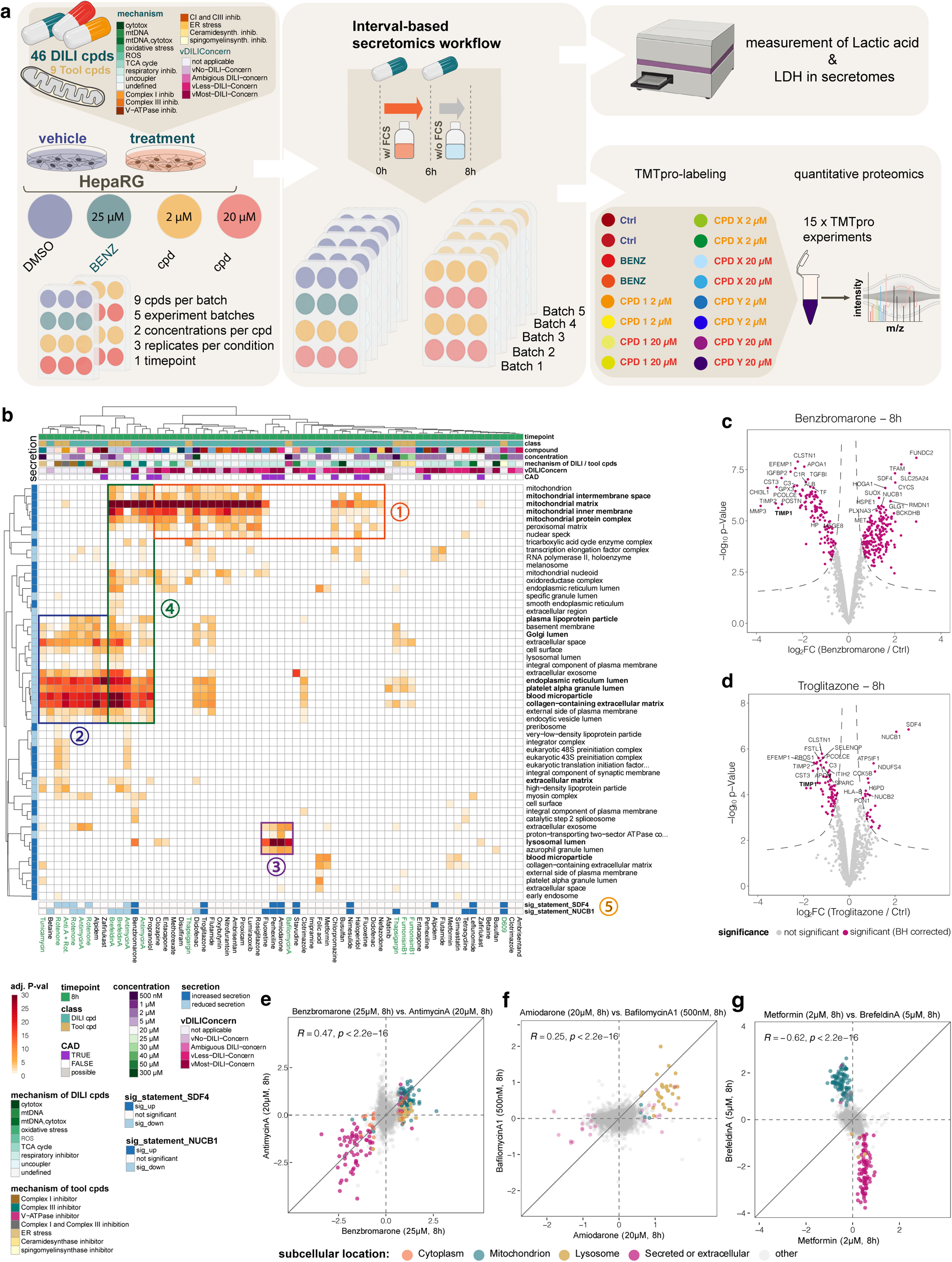
DILI compounds induce the secretion of proteins from distinct cellular organelles. **a**, Schematic representation of the interval-based secretomics screening using DILI- and tool-compounds in dHepaRG cells. **b**, Heatmap displaying significant GO-terms (cellular component) derived from the differential secretome analysis of DILI- and tool-compound treated HepaRG cells (n=2) versus the time-matched control (n=2). Differentially secreted proteins were determined via LIMMA. Significance thresholds were (p(Benjamini Hochberg) < 0.05 and log_2_ fold change > 2x standard deviation of the individual treatment. Color intensities indicate the adjusted (BH-corrected) p-value (-log_10_-transformed) of the GO-term. Only GO-terms are displayed that were significant upon three or more compounds. Rows are clustered by Pearson correlation; columns are clustered by Euclidian distance. Row annotation indicates whether proteins were downregulated in their secretion (light blue) or upregulated in their secretion (dark blue) upon compound treatment. Characteristic clusters of DILI- and tool-compounds are grouped: (1) compounds that induced the secretion of mitochondrial proteins, (2) compounds which perturbed the canonical secretion of proteins; (3) compounds, which induced the secretion of lysosomal proteins; (4) compounds which induced mixed effects. (5) compounds which induced a secretion of Golgi-resident proteins NUCB1 and SDF4. **c**, Volcano plot showing proteins quantified in the secretome of benzbromarone treated dHepaRG cells 8h post stimulus. Displayed are the log_2_ fold changes and the p-values (-log_10_-transformed) determined by LIMMA of benzbromarone treated dHepaRG cells (n=3) versus the time matched vehicle (DMSO)-controls (n=3). Dotted line indicates significance cut-offs. Proteins passing the significance thresholds (p(Benjamini Hochberg) < 0.05 and log_2_ fold change > 2x standard deviation of the individual treatment are colored in purple. d, same as c for 8h troglitazone treatment. **e-g**, Pairwise correlation plots of compound induced secretion patterns of: **e**, benzbromarone and antimycin A. **f**, amiodarone and bafilomycin A1, **g**, metformin and brefeldin A. x- and y-axes are represented as log_2_ fold change (compound vs. DMSO). Each point represents a protein, color represents the subcellular location based on the UniprotKB annotation: purple: proteins annotated as secreted or extracellular, teal: mitochondrial proteins, yellow: lysosomal proteins, orange: cytoplasmic proteins, grey: proteins with subcellular locations other than dose described. Pearson correlation (R) is given, statistical significance is given by the p-value (p).

For quantitative proteomics analysis samples were encoded with isobaric mass-tags ^18, 19^ and subjected to multiplexed mass spectrometric analysis where two vehicle controls and two benzbromarone treatment samples were combined with three DILI compound treatments covering both concentrations in duplicate (the two samples with lowest LDH signal were selected) (Fig. 1a). In total 488 secretome samples were analyzed in 39 multiplexed experiments leading to a total of 321 LC-MS runs after fractionation.

On average 3000 ± 200 proteins were detected in supernatants of dHepaRG cells of which 179 ± 20 were annotated in UniProtKB ^20^ either as secreted or extracellular (Supplementary Table 3a, 3b). Whilst the majority of proteins detected in the cell culture supernatants remained unaltered under conditions where lactate levels / LDH activity were not increased, drug treatment affected protein levels in the secretome spanning from one protein for Nimesulide and Simvastatin (2 µM) to 593 proteins for Antimycin A (20 µM) (Supplementary Table 3b). We observed a trend towards more differentially regulated secreted proteins at higher compound concentrations, whereby an increased secretion could be frequently (-log_10_ adj. P-value (enrichment) < 0.05) observed for proteins located in the mitochondrion and cytoplasm, whilst reduced secretion was frequently observed for proteins annotated as secreted or extracellular (Supplementary Table 3b).

Comparing the intensity distribution of proteins in the total dHepaRG proteome with the cell culture supernatants from e.g. the prototypical DILI compounds benzbromarone and troglitazone (Supplementary Fig. 2a-d), revealed no bias towards the most abundant intracellular proteins in the secretome confirming that the compound induced secretion events cannot be explained by cell leakage.

Looking into the secretomes obtained with benzbromarone (uricosuric agent) and troglitazone (PPARγ agonist) in some detail, we observed the rapid onset of secretion of mitochondrial, cytoplasmic, nuclear and lysosomal proteins and the two Golgi resident proteins SDF4 and NUCB1 at both timepoints (Fig. 1b-d, Supplementary Fig. 2e-h) as well as reduced basal secretion of acute-phase proteins, extra-cellular matrix proteins and lipoproteins, e.g. TIMP1 or APOB (Fig. 1c-d, Supplementary Fig. 2e-f). Capillary western blot (WES) analysis of TIMP1 in the supernatants after 8h (Supplementary Fig. 2i), confirmed impaired TIMP1 secretion and provided further evidence that the observed secretion events were not the result of compromised cell viability, since cell leakage would lead to higher TIMP1 abundances in the cell culture supernatant. Mitochondrial proteins in the secretome were mostly enriched from the mitochondrial matrix (Supplementary Fig. 3a-c), suggesting secretion of a subset of the mitochondrial proteome.

GO-term enrichments for the secretomes obtained for the entire compound set indicated that significantly regulated proteins in the secretome originated from distinct subcellular compartments, including mitochondria, ER, and lysosomes (Fig. 1b). Hierarchical clustering using enriched GO-terms identified distinct groups of DILI compounds including:

(1) compounds that induced the secretion of mitochondrial proteins, (e.g disulfiram, ambrisentan, piroxicam, troglitazone, benzbromarone), which were mostly enriched from the mitochondrial matrix, including proteins involved in fatty acid metabolism or the TCA cycle; (2) compounds which perturbed the canonical secretion of proteins such as acute-phase proteins, extra-cellular matrix or lipoproteins (e.g. alpidem, betaine, rotenone, troglitazone, diclofenac); (3) compounds which induced the secretion of lysosomal proteins such as cathepsins and glycosidases (e.g. amiodarone, fluoxetine, perhexiline, zafirlukast, benzbromarone); (4) compounds which induced mixed effects, e.g. the secretion of mitochondrial proteins into the cell culture supernatant and perturbation of the canonical secretion of proteins (e.g. diclofenac, troglitazone, benzbromarone, propranolol, flutamide). In addition, a number of compounds (e.g. troglitazone, benzbromarone fluoxetine, perhexiline, amiodarone) induced the secretion of Golgi-resident proteins NUCB1 and SDF4 (5).

Compounds with similar secretion profiles were not necessarily characterized by structural similarities or shared primary indications or cellular targets. Even structurally related compounds with the same cellular target and similar indication, like the thiazolidinedione-based PPARγ agonists troglitazone, rosiglitazone, and pioglitazone, displayed very different secretome-modulating effects (Fig. 1b). For instance, pioglitazone (less DILI concern) had no impact on secretion, while rosiglitazone (less DILI concern) induced the secretion of mainly mitochondrial proteins, and troglitazone (most DILI concern) induced the secretion of mitochondrial proteins and impaired basal secretion.

Tool compounds with distinct mechanisms affecting specific organelles exhibited similar acute organellar secretion patterns and shared clusters with DILI compounds (Fig. 1b). For instance, the complex I inhibitor rotenone impaired basal protein secretion und clustered into the same group as e.g. alpidem, whilst the complex III inhibitor antimycin A impaired basal secretion, induced the non-conventional secretion of mitochondrial proteins and clustered into the same group as benzbromarone or troglitazone (Fig. 1b). The V-ATPase inhibitor bafilomycin A1 mainly induced the secretion of lysosomal proteins und clustered together with the cationic-amphiphilic drugs (CADs) amiodarone and perhexiline (Fig. 1b). The finding that DILI-compound induced secretion patterns can be related to distinct intracellular mechanism dysregulated by these compounds was further characterized by direct correlation of secretome profiles with those of tool compounds (Supplementary Fig. 4a):

The secretion profile of benzbromarone, a known respiratory chain inhibitor ^21^, correlated well with that obtained for the secretion inhibitor brefeldin A ^22^ (Supplementary Fig. 4b) and the complex III inhibitor antimycin A ^23^ (Fig. 1e), suggesting impairment of mitochondrial function and a functional deficit in the secretory pathway by benzbromarone. A similar correlation could be observed upon comparison of the tool compounds antimycin A and brefeldin A (Supplementary Fig. 4c), suggesting that also tool compounds with known MoA have additional, yet undescribed effects on cellular organelles. The beta-blocker propranolol and the ER-stress inducer thapsigargin also correlated, inducing the release of mitochondrial proteins and impairing basal protein secretion (Supplementary Fig. 4d) whereas the CAD amiodarone and the V-ATPase inhibitor bafilomycin (Fig. 1f) which blocks lysosomal acidification both induced the release of lysosomal proteins. In contrast, the type 2 diabetes drug metformin, had a secretion profile that was anticorrelated with that of brefeldin A (Fig. 1g) and other tool compounds (Fig. 4a), indicating a positive effect on cell viability which is in agreement with the DILI-rank annotation ‘less DILI concern’.

Taken together, we observed distinct clusters of protein secretion in response to treatment of the dHepaRG liver cell model with DILI compounds including non-canonical secretion of proteins from lysosomes and mitochondria and other organelles. These non-canonical secretion effects could be mimicked by tool compounds with known mechanism of action on the respective organelles. This let us hypothesize that a surprisingly large range of intracellular mechanisms in distinct organelles are dysregulated by non-related DILI compounds, and that these dysregulations manifest in the secretome of liver cells. We therefore set out to investigate the intracellular mechanisms responsible for these observed secretome signatures using exemplary DILI- and tool compounds (Supplementary Table 2).

### Thermal proteome profiling confirms organelle specific effects of DILI compounds

Next, we characterized intracellular effects of a subset of the DILI drugs and tool compounds by Thermal Proteome Profiling (TPP) in dHepaRG cells (Fig. 2a). TPP is a quantitative mass spectrometry technique that measures drug effects on protein thermal stability and protein abundances across the proteome, thus enabling identification of direct drug targets as well as indirect drug effects ^24–27^.

**Fig. 2.**
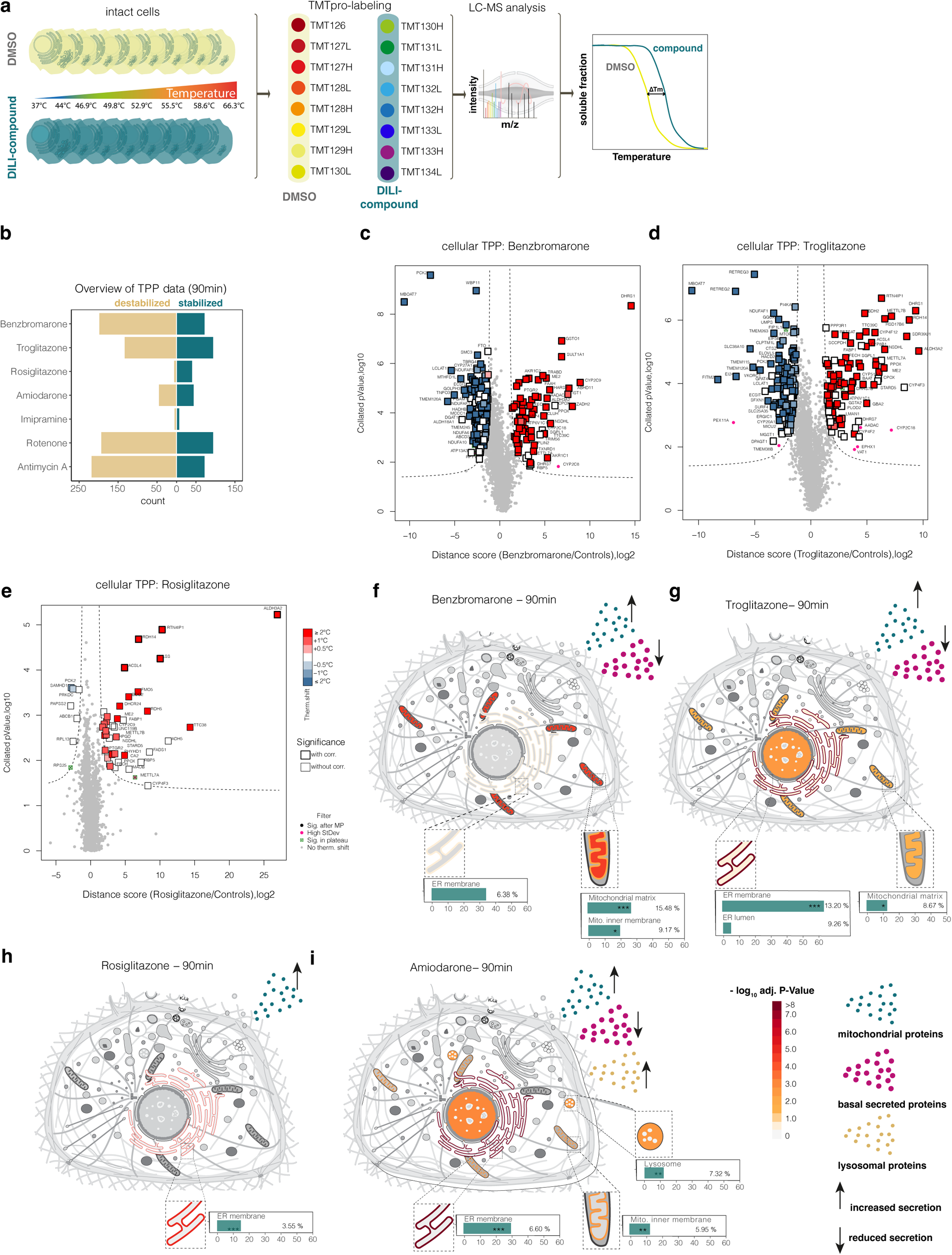
DILI compounds induce widespread protein thermal stability changes in dHepaRG cells. **a**, Schematic overview of the cellular TPP workflow using intact dHepaRG cells and TMTpro-labeling scheme allowing the combination of control- and treated samples in a single MS-run. Data from two independent experiments per treatment have been analyzed. **b**, Overview of protein thermal stability changes after 90 min of compound treatment. Bar graphs showing number of significant proteins stabilized (green) or destabilized (brown) grouped by treatment. **c**, Protein thermal stability changes in dHepaRG cells upon treatment with benzbromarone (90 min). Volcano plot displays distance scores and collated P values (-log10 transformed) of proteins quantified in benzbromarone (25 µM, n=2) versus DMSO-treated (n=2) dHepaRG cells. The ratio-based approach included LIMMA analysis for ratios between treatment and control groups obtained at each temperature, aggregation of retrieved P values per protein by Brown’s method and multiple testing adjustment Benjamini–Hochberg (BH) correction. Dotted line indicates significance cut-offs. Proteins passing significance cut-off are colored according to their Tm shift, bold edges indicate Benjamini-Hochberg corrected P-values with P< 0.05, light gray dots depict proteins that were not significantly affected. **d**, same as c for the treatment with 20 µM troglitazone (90 min). **e**, same as c for 20 µM rosiglitazone (90 min). **f**, Total count of proteins with significant protein thermal stability changes upon 90 min treatment with benzbromarone grouped by their subcellular location. Percentage values represent the protein fraction relative to the average protein count of the respective cellular organelle. Asterisks denote significant enrichments for proteins of a respective cellular organelle encoded by: P ≤ 0.05: *; P ≤ 0.01: **; P ≤ 0.001: ***. P-values were calculated with a Fisher-exact test. **g**, same as f for troglitazone. **h**, same as f for rosiglitazone. **i**, same as f for amiodarone.

The selected compounds mainly affected protein thermal stability (Fig. 2b-e) whilst with the exception of the tool compounds rotenone and antimycin A1 only a few proteins were altered in abundance after 90min of cell treatment (Supplementary Fig. 5a, b, Supplementary Table 4). Cell treatment with rosiglitazone and the CAD impramine, both annotated with ‘less DILI concern’, induced only few thermal stability changes (Fig. 2b, Supplementary Fig. 5b). In contrast, benzbromarone, troglitazone and amiodarone that are annotated with ‘most-DILI concern’ and the tool compounds antimycin A and rotenone induced thermal stability changes for a broad range of proteins, with the vast majority being destabilized (Fig. 2b, Supplementary Fig. 5b).

For most of the tested compounds, we did not expect to detect binding to their primary targets due to very low or complete absence of expression in the dHepaRG cell system. An exception was antimycin A for which we observed strong thermal stabilization of respiratory complex III subunits (Supplementary Table 4). By contrast, we observed thermal stability changes for several previously reported off-targets such as stabilization of the benzbromarone off-target FABP4 ^28^ (Fig. 2e, Supplementary Table 4), stabilization of the troglitazone and rosiglitazone off-target ACSL4 ^29^ (Fig. 2d-e) as well as stabilization of cytochrome P450 enzymes CYP2C8 and CYP2C9 by troglitazone in line with reported inhibition by the drug ^30^. Similarly, Rotenone - despite not having a significant stabilization effect on mitochondrial complex I subunits - induced thermal stabilization of microtubules and proteasome-subunits consistent with the reported inhibition of the formation of microtubules and decreased proteasome activity ^31, 32^.

Enrichment analysis revealed that compound induced thermal stability effects clustered in distinct cellular organelles affecting in many cases predominantly membrane proteins (Fig. 2f-i, Supplementary Fig. 6a-k, Supplementary Table 5). For example, troglitazone led to a significant enrichment of proteins located in the ER membrane (log_10_ adj. P-value = 20.30), the cytoplasm (log_10_ adj. P-value = 6.36), the mitochondrial matrix (log_10_ adj. P-value = 1.43) and the nucleus (log_10_ adj. P-value = 2.97) (Fig. 2g, Supplementary Fig. 6c, Supplementary Table 5), which is in line with the observed effects on protein secretion (increased secretion of mitochondrial proteins and reduced basal protein secretion). In contrast, for rosiglitazone treatment significant enrichment of stability changes was limited to the ER membrane (log_10_ adj. P-value = 5.17, Fig. 2h, Supplementary Fig. 6d), likely mediated through rosiglitazone metabolism by ER resident enzymes, including CYP2C9 ^33^ and as evidenced by the enriched GO term “oxidation-reduction process” (Supplementary Table 6). Moreover, thermal stability changes on several mitochondrial proteins suggests a causal relationship with the increased secretion of mitochondrial proteins described above. The lysosomotropic drug amiodarone (Fig. 2i, Supplementary

Fig. 6e, Supplementary Table 5) significantly affected lysosomal proteins (log_10_ adj. P-value = 2.52) in addition to the ER membrane (log_10_ adj. P-value = 8.69), the cytoplasm (log_10_ adj. P-value = 4.74), and the inner mitochondrial membrane (-log_10_ adj. P-value = 2.48), which is in line with the observed effects on protein secretion (increased secretion of mitochondrial proteins, reduced basal protein secretion and increased secretion of lysosomal proteins). Similar correlations between observed thermal stability changes and secretion profiles were observed for most of the tested compounds (Supplementary Fig. 6a-k, Supplementary Table 5) thus, corroborating the hypothesis that a diverse range of intracellular mechanism in distinct organelles are dysregulated by non-related DILI compounds that are reflected in the observed secretome signatures.

### Thermal stabilities of multiple transmembrane proteins are affected by DILI compounds

Whilst compounds with ‘less DILI concern’ such as rosiglitazone mainly affected proteins without or with only one TM domain (Fig. 3a), DILI compounds such as benzbromarone, troglitazone, amiodarone, antimycin A elicited widespread thermal stability changes with a trend towards thermal destabilization for proteins with three or more transmembrane domains (Fig. 3a, Supplemental Fig. 7a-c). When reducing the troglitazone concentrations 20-fold to 1 µM (6-fold below Cmax), Tms of only seven transmembrane proteins were affected and the trend towards destabilization disappeared demonstrating that this is a concentration-dependent effect (Fig. 3b). In time-dependent studies with troglitazone, however, effects on proteins with more than three transmembrane domains did not fundamentally change suggesting that these effects are likely directly caused by the parental molecule (Supplementary Fig. 7d-f).

**Fig. 3.**
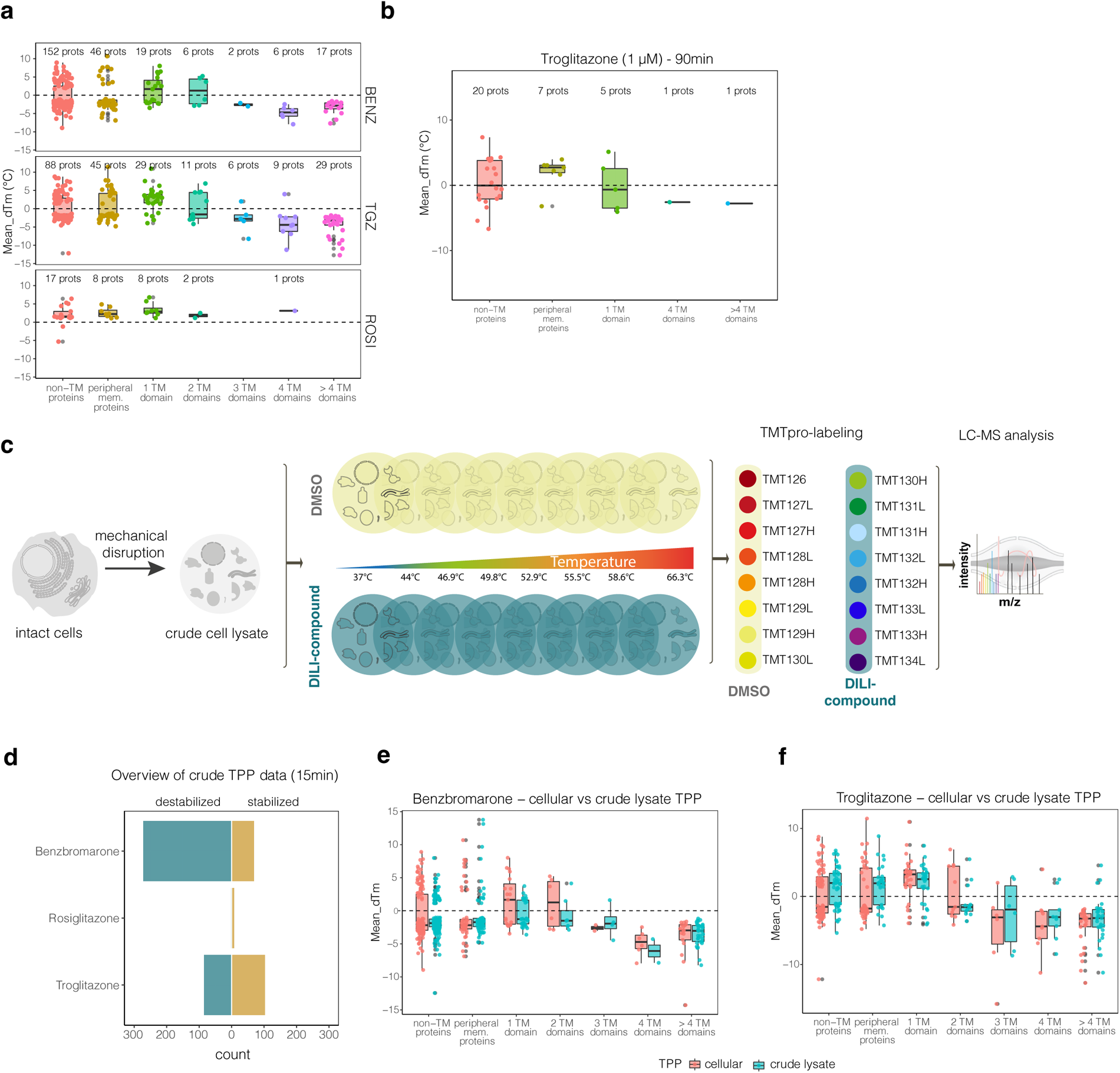
DILI compounds affect the thermal stabilities of a multitude of transmembrane proteins. **a**, Comparison of mean dTm (°C) for proteins with significant thermal stability shift > 1 °C or < 1 °C upon benzbromarone, troglitazone and rosiglitazone treatment (90 min) of dHepaRG cells and grouped by their number of transmembrane domains as annotated in UniprotKB. Number of proteins per group are indicated. Dotted line denotes a dTm of 0. Center line, median; box limits, upper and lower quartiles; whiskers, maximum and minimum value of the dataset. Points denote the dTm value for each protein in the respective group. **b**, same as a for the treatment with 1 µM troglitazone (90 min). **c**, schematic representation of the crude lysate TPP workflow for the assessment of direct drug effects. Intact dHepaRG cells are mechanically disrupted. The resulting crude extract is split into four aliquots and either treated with DMSO as vehicle control or with the DILI compound and subsequently heated to different temperatures. **d**, Overview of protein thermal stability changes after 15 min of compound treatment using crude HepaRG cell lysates. Bar graphs showing number of significant proteins stabilized (green) or destabilized (brown) grouped by treatment. **e**, Comparison of mean dTm (°C) for proteins with significant thermal stability shift from cellular TPP (red) and crude lysate TPP experiments (blue) with benzbromarone and grouped by their number of transmembrane domains as annotated in UniprotKB. Dotted line denotes a dTm of 0. Center line, median; box limits, upper and lower quartiles; whiskers, maximum and minimum value of the dataset. Points denote the dTm value for each protein in the respective group. **f**, same as **g** for troglitazone

When repeating TPP experiments in crude cell extracts ^34^ (Fig. 3c-d) generated by mechanically disrupting dHepaRG cells in absence of detergents, we observed a similar trend towards destabilization of proteins with three or more TM domains for benzbromarone and troglitazone but not for rosiglitazone (Fig. 3d) consistent with observations in the cell-based TPP experiments (Fig. 3e-f).

As downstream signaling effects affecting protein thermal stability of membrane proteins can be practically excluded under these conditions and upon short term (15 min) incubation with compounds, we conclude that the observed widespread effects on membrane protein stability are a direct consequence of DILI compound accumulation at or in the affected membrane compartments and off-target binding.

This is consistent with previous studies for thiazolidinediones like troglitazone that reported off-target effects for many transmembrane proteins which are independent of PPARγ.

In summary, we demonstrate that DILI compounds affect a specific subset of transmembrane proteins localized in specific organelles such as ER membrane and mitochondria and these effects are reflected in the corresponding secretomes where proteins from these cellular compartments are enriched.

### Secretomics and thermal proteome profiling of lysosomotropic compounds

The DILI compound set contained several compounds classified as cationic amphiphilic drugs (CADs) ^35^ such as amiodarone, perhexiline , fluoxetine, zafirlukast and clozapine and molecules with CAD-like structures (e.g. entacapone). CADs are characterized by a hydrophobic aromatic ring system and a hydrophilic sidechain with an ionizable amine functional group ^36^ allowing them to interact with- and to diffuse across lysosomal membranes. Once inside the acidic lysosomal lumen, the amine is protonated^37, 38^, leading to trapping and CAD accumulation, potential pH changes^35, 36, 39–41^, altered lipid metabolism, and possibly drug-induced phospholipidosis (DIPL) ^37^.

In our secretomics screen (Fig. 1b), most CADs including amiodarone, perhexiline, fluoxetine, zafirlukast and clozapine induced the secretion of lysosomal proteins after 8h when tested at 20 µM but not at 2 µM. Exceptions were propranolol, imipramine and chlorpromazine, which only induced the release of mitochondrial proteins or reduced basal secretion, suggesting preferential accumulation of these compounds in the ER and mitochondria (Fig. 1b). The released lysosomal content preferentially contained soluble lysosomal proteins, such as the cathepsins CTSL, CTSA, CTSD, CTSZ, CTSH or the hydrolases MAN2B1, SIAE, HEXA and HEXB (Fig 4a, b). Repeat experiments reading out after 2 h did not yield any lysosomal content in supernatants consistent with a time and concentration dependent effect.

In order to probe that this is indeed a feature common to drugs with CAD-like properties, we included the EGFR inhibitor afatinib in these experiments which has not yet been flagged as CAD but has been observed to be trapped in lysosomes ^42^. When comparing the secretomes induced by afatinib (logP: 3.6, pKa: 8.81, 10 µM) to that of lapatinib (logP: 5.1, pKa: 7.2, 20 µM) a less basic inhibitor directed against the same targets, we observed an enrichment of soluble lysosomal proteins in the cell culture supernatants only with afatinib (Fig. 4c-e, Supplementary Fig. 8a-b) and protein identities agreed well with those secreted with the prototypic CAD amiodarone (logP: 7.6 and pKa 8.47) that is more similar in basicity. In summary these results suggest that the secretion profiles of CADs and their lysomotropic properties are both concentration and time dependent and strongly influenced by the basicity of the molecules.

Lysosomal accumulation of CADs is thought to shift the pH towards more alkaline conditions which blocks dissociation of lysosomal proteins from the mannose-6 phosphate receptor (MPR300/IGF2R) that transports lysosomal proteins to the endolysosomal compartment and impairs recycling of MPR300 back to the TGN ^35, 43, 44^. The direct link of lysosomal protein secretion to lysosomal pH changes was confirmed by blocking proton transport into the lysosome with the V-ATPase inhibitor bafilomycin A1 ^45^. Similar to CADs, Bafilomycin A1 induced the secretion of lysosomal proteins into the secretome after 8h in the dHepaRG model (Fig. 4f-g, Supplementary Fig. 8c-d). Moreover, we observed a decrease in fluorescence of acidic vesicular structures stained with Lysotracker Green DND-26 upon treatment with bafilomycin A1 in HepG2 cells after 2 and 8h (Supplementary Fig. 8e), thus confirming the disruption of the acidic lysosomal environment. Taken together these data confirm a direct link between the lysosomal pH and lysosomal protein release.

In TPP experiments with bafilomycin A1 (90 min incubation, Fig. 4h) the thermal stabilization of the membrane-localized V-ATPase subunits ATP6V0A1 and ATP6V0A2 was observed , which confirms previous reports that identified the V0 subunit as molecular target of bafilomycin A1^45^. Consistent with the secretomics data, we observed a decrease in abundance and thermal stability of soluble lysosomal proteins and an increase in abundance of basal secreted proteins, e.g. ECM proteins indicating blocked conventional secretion (Fig. 5i, Supplementary Fig. 8f-g). The observed thermal stability changes in the lysosome were very similar to those observed with the classic CAD amiodarone (Fig. 4j). Notably, amiodarone stabilized the cytoplasmic V-ATPase subunits ATP6V1A, ATP6V1C1, ATP6V1D, ATP6V1E1 but not V0 suggesting a different, potentially allosteric mechanism for modulating lysosomal pH and protein transport (Fig. 4j-k).

**Fig. 4.**
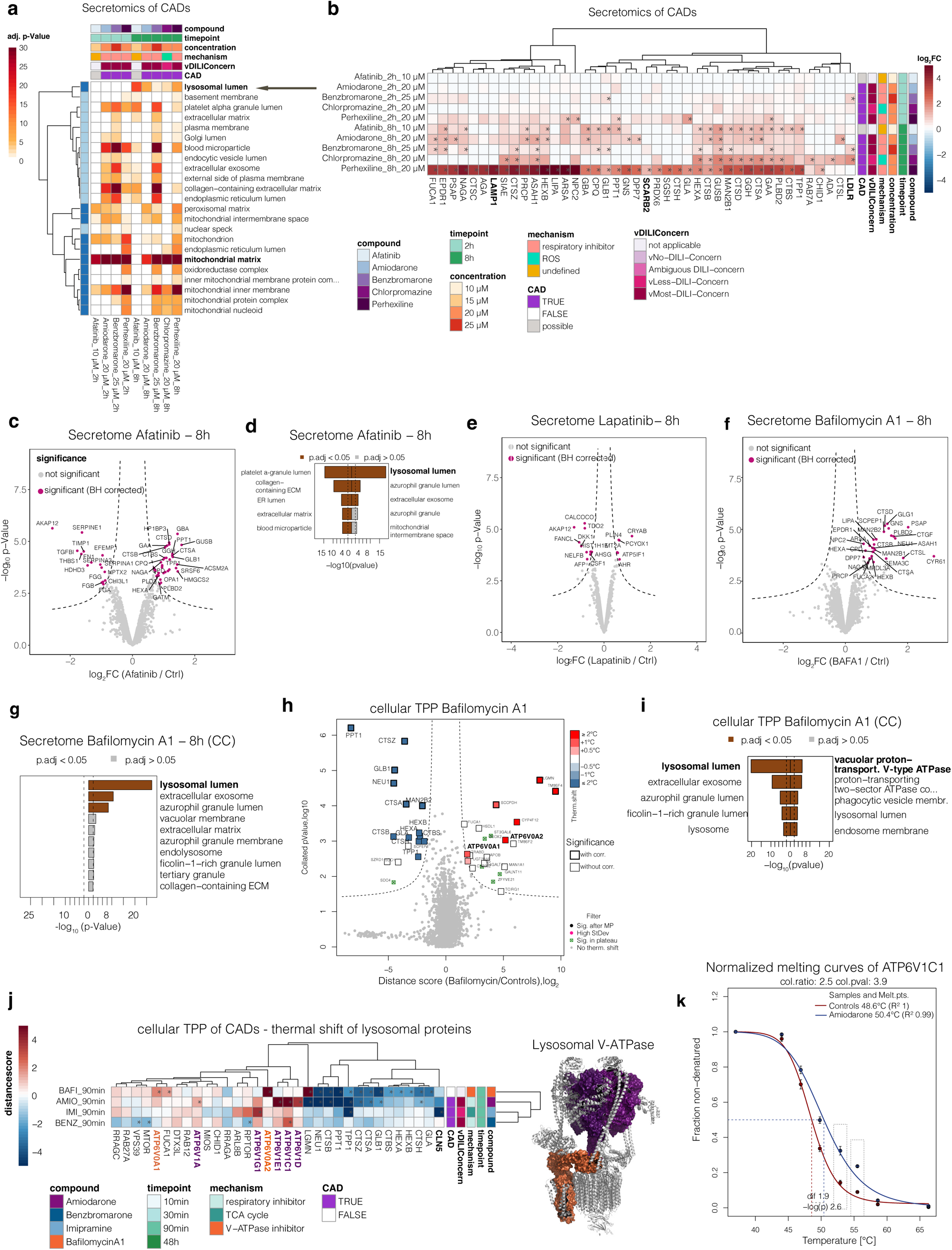
CADs induce a time- and concentration dependent release of lysosomal proteins into the cell culture supernatant. **a**, CADs induce the secretion of lysosomal proteins. Heatmap displaying significant GO-terms (cellular component) derived from the differential secretome analysis of CAD treated HepaRG cells (n=3) versus the time-matched control (n=3 for 2h and n =2 for 8h). Differentially secreted proteins were determined via LIMMA. Significance thresholds were (p(Benjamini Hochberg) < 0.05 and log_2_ fold change > 2x standard deviation of the individual treatment. Color intensities indicate the adjusted (BH-corrected) p-value (-log_10_-transformed) of the GO-term. Only GO-terms are displayed that were significant upon three or more CADs. Rows are clustered by Pearson correlation. Row annotation indicates whether proteins were downregulated in their secretion (light blue) or upregulated in their secretion (dark blue) upon compound treatment. Interesting GO-terms are indicated in bold. **b**, Heatmap displaying lysosomal proteins found in the secretome upon treatment of dHepaRG cells with CADs across different timepoints irrespective of their significance. Displayed are log_2_ fold changes to the respective time matched control. Statistically significant changes are denoted with asterisks (*). **c**, Changes in protein secretion upon treatment with 10 µM afatinib (8h). Volcano plot showing proteins quantified in the secretome of afatinib treated dHepaRG cells 8h post stimulus. Displayed are the log_2_ fold changes and the p-values (-log_10_-transformed) determined by LIMMA of afatinib treated dHepaRG cells (n=3) versus the time matched vehicle (DMSO)-controls (n=3). Dotted line indicates significance cut-offs. Proteins passing the significance thresholds (p(Benjamini Hochberg) < 0.05 and log_2_ fold change > 2x standard deviation of the individual treatment are colored in purple. **d**, Top ten most significantly enriched GO-terms (cellular component) in the secretome of afatinib treated dHepaRG cells. GO enrichment was tested with an Fisher’s exact test. Significant GO-terms (P(BH corrected) < 0.05) are depicted as brown bars. **e**, same as c for the treatment with 20 µM lapatinib; 8h post stimulus. **f**, same as c for the treatment with the V-ATPase inhibitor bafilomycin A1, 8h post stimulus. **g**, same as d for Bafilomycin A1 (8h). **h**, Protein thermal stability changes in dHepaRG cells upon treatment with 500 nM bafilomycin A1 (90 min). Volcano plot displays distance scores and collated P values (-log10 transformed) of proteins quantified in bafilomycin a1 (20 µM, n=2) versus DMSO-treated (n=2) dHepaRG cells. The ratio-based approach included LIMMA analysis for ratios between treatment and control groups obtained at each temperature, aggregation of retrieved P values per protein by Brown’s method and multiple testing adjustment Benjamini–Hochberg (BH) correction. Dotted line indicates significance cut-offs. Proteins passing significance cut-off are colored according to their Tm shift, bold edges indicate Benjamini-Hochberg corrected P-values with P< 0.05, light gray dots depict proteins that were not significantly affected. **i**, Top five most significantly enriched GO-terms (cellular component) in the TPP of Bafilomycin A treated dHepaRG cells. GO enrichment was tested with a Fisher’s exact test. Significant GO-terms (P(BH corrected) < 0.05) are depicted as brown bars. **j**, Thermal shift of lysosomal proteins and impact of CADs on V-ATPase subunits. Heatmap displaying lysosomal proteins exhibiting a thermal stability shift upon treatment of dHepaRG cells with CADs for 90min irrespective of their significance. Displayed are distancescores. Statistically significant changes are denoted with asterisks (*). Proteinstructure shows the lysosmal V-ATPase. Purple colored subunits show the cytoplasmic V1 portion of the V-ATPase. Organge colored subunits show the V0 portion of the V-ATPase. **k**, reconstructed median melting curves for the cytoplasmic V-ATPase subunit ATP6V1C1 in the control- and amiodarone treated cells. Data are presented as median ± s.d. Mean melting point shifts (dTm) and corresponding P values (LIMMA) are indicated.

**Fig. 5.**
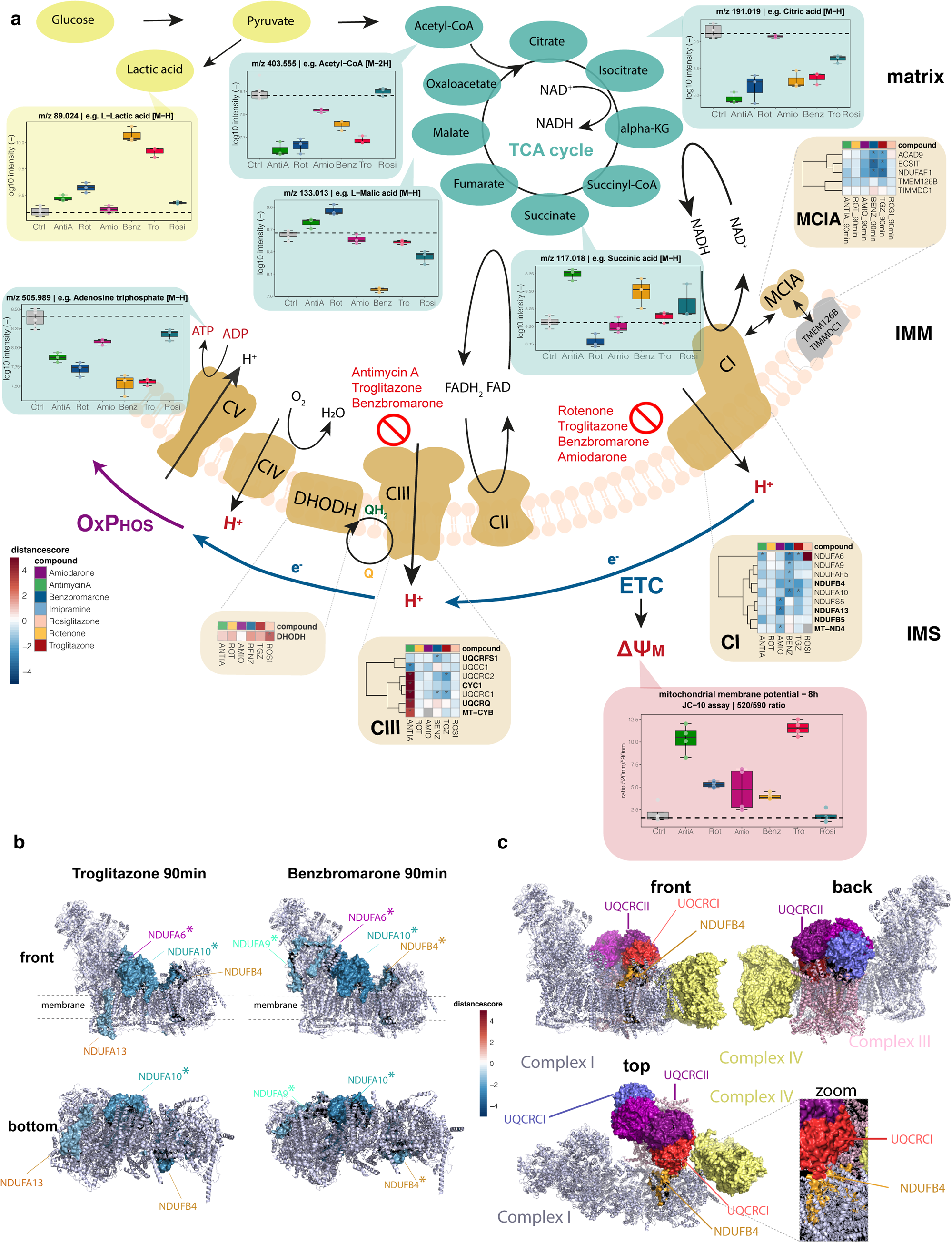
DILI compounds with mitochondrial secretion acutely destabilize proteins of the inner mitochondrial membrane and induce metabolic perturbations. **a**, Schematic overview of the respiratory chain and the observed thermal stability effects (brown boxes), changes in metabolites (yellow and green boxes) and changes in the mitochondrial membrane potential (MMP) in mitochondria upon DILI compound treatment (red box). Heatmaps in brown boxes displaying proteins of the respiratory chain exhibiting a thermal stability shift upon treatment of dHepaRG cells with DILI compounds for 90min irrespective of their significance. Displayed are distancescores. Statistically significant changes are denoted with asterisks (*). Boxplots (yellow or green boxes) depict MS1 intensities (-log_10_ transformed) of glycolytic metabolites (yellow box) or TCA cycle metabolites (green boxes) compound treatment. Data points denote MS1 intensities derived from each of the three independent biological replicates. Boxplots in the purple box display changes of the mitochondrial membrane potential as ratios of the fluorescence measurement at 520 and 590nm determined with an JC-10 assay after 8h of compound treatment. Points denote the 520/590nm ratios derived from 4 independent biological replicates. **b**, Molecular visualization of MRC complex I (structure was derived from PDB database, identifier: 5XTH). Upper panel shows the front view, lower panel shows the bottom view. Surface representations indicate thermal destabilized subunits after 90 min compound treatment with the MRC inhibitors troglitazone (left) and benzbromarone (right). Color indicates the distancescore. Dotted lines represent the inner mitochondrial membrane. Statistically significant thermal stability changes are denoted with asterisks (*). **c**, Molecular visualization of the human respiratory supercomplex I1III2IV1 (respirasome, structure was derived from PDB database, identifier: 5XTH). Surface representations indicate CI, CIII and CIV subunits of interest. Zoom shows interaction-site of CI subunit NDUFB4 with the CIII subunit UQCRCI.

In summary, CADs induced a time- and concentration dependent release of lysosomal proteins into the cell culture supernatant 8h post treatment and thermal destabilization of soluble lysosomal proteins, consistent with accumulation of CADs accumulate in acidic compartments. Both, lysosomal protein release and thermal destabilization of these proteins at earlier time points were phenotypical equivalent to blocking the H^+^ transporting V-ATPase with bafilomycin A1 and can be considered as mechanistic biomarkers for molecules that accumulate in acidic organelles and interfere with lysosomal acidification.

### DILI compounds with mitochondrial secretion acutely destabilize proteins of the inner mitochondrial membrane

Release of mitochondrial proteins into cell culture supernatants was observed for 27 compounds amongst which many have previously been identified as respiratory chain inhibitors, e.g., benzbromarone, diclofenac, alpidem, troglitazone (Fig. 1b, Supplementary Fig. 9a). Mitochondrial protein release was already observed 2h post treatment showing strong enrichment of proteins from the mitochondrial inter-membrane space and the mitochondrial matrix (Supplementary Fig. 3a, Supplementary Fig. 9a, Supplementary Fig. 10a-b). These protein subsets are characterized by containing reactive cysteine residues and iron-sulfur clusters (Supplementary Fig. 3c, Supplementary Fig. 10c) and are involved in fatty acid beta oxidation, TCA cycle, or acyl-CoA metabolic processes (Supplemental Fig. 3b). Notably, for the majority of compounds, we observed the secretion of a number of core- and accessory subunits of complex I situated in the N-and Q modules (Supplementary Table 3a, 3c).

Next, we investigated how tool molecules inhibiting the mitochondrial respiratory chain influenced release of mitochondrial proteins. Even at concentrations as high as 20 µM, the complex III inhibitor antimycin A induced mitochondrial protein release only at the 8h time point whilst complex I inhibition with rotenone led to reduced basal secretion, but no release of mitochondrial proteins was observed (Fig. 1b, Supplementary Fig. 9a). However, when cells were treated with both inhibitors, mitochondrial protein release was already observed after 2h, indicating at least additive effects. In summary, these data suggest that the acute release of mitochondrial proteins is a reaction to mitochondrial stress resulting from acute inhibition of the mitochondrial respiratory chain by DILI compounds.

To follow up on this hypothesis we used TPP to investigate thermal stability effects in mitochondria of a set of DILI and tool molecules inducing mitochondrial protein release (Fig. 5a. Supplementary Fig. 9b-e).Whilst no thermal stability effects were observed in complex I upon rotenone treatment, antimycin A stabilized its known target cytochrome B (MT-CYB ) of CIII as well as the complex III subunits UQCRC1, UQCRC2, CYC1, UQCRQ (Fig. 5a, Supplementary Fig. 9b-c)demonstrating direct binding to the CIII complex.

For cells treated with the DILI compounds troglitazone, benzbromarone and amiodarone, we observed substantial destabilization of multiple TM proteins of the inner mitochondrial membrane. (Fig. 5a, b and Supplementary Fig. 9c), including several subunits of CI and CIII.

Following up in a time course, significant destabilization of different membrane-spanning subunits within complex Ìs proton transporting modules ^46^ ND4 (NDUFB4, NDUFB5, MT-ND4), and ND1 (NDUFA13) was already observed 10 min after treatment of dHepaRG cells with troglitazone (Supplementary Fig. 11a). After 90 min we found that troglitazone destabilized complex I subunits located in the matrix arm modules ND2 (NDFA10) and N (NDUFA6) (Fig. 5a-b, Supplementary Fig. 11a). A similar picture was observed with benzbromarone and amiodarone, both destabilizing membrane-spanning subunits within the ND4-module (NDUFB4 or MT-ND4) and within the matrix arm localized ND2-module (NDUFA10 and NDUFS5) (Fig. 5a-b, Supplementary Fig. 11b). Moreover, amiodarone destabilized ND1-module subunit NDUFA13, whilst benzbromarone additionally destabilized the N- and Q-module subunits NDUFA6 and NDUFA9. Notably, troglitazone and benzbromarone both destabilized the CI matrix arm-associated MCIA complex (ACAD9, ECSIT, NDUFAF1), which was previously shown to be critical for complex I assembly and stability (Fig. 5a) ^47, 48^ as well as different complex III subunits such as UQCRC1, UQCRC2 and UQCRFS1 (Fig. 5a). Notably, CI subunit NDUFB4 is known to be in close contact with CIII subunit UQCRC1^49, 50^ (Fig. 5c). These results suggest that troglitazone, benzbromarone and amiodarone indirectly affect complexes I and III function by destabilizing the CI-membrane interface.

Further, we observed numerous other proteins of the inner mitochondrial membrane to be thermally destabilized upon compound treatment such as SFXN1, OXA1L, HADHA, HADHB, and various solute carrier proteins, fulfilling critical functions for the respiratory chain and mitochondrial integrity, such as organization of mitochondrial cristae structure (Supplementary Fig. 9b, Supplementary Table 4).

Interestingly, the insulin-sensitizing drug rosiglitazone which is thought to exert its antidiabetic effect in a similar fashion like troglitazone through inhibition of complex I, did not induce significant thermal stability changes in complex I or complex III subunits (Fig. 5a, Supplementary Fig. 9c). Instead, we observed a significant thermal stabilization of DHODH, which provides ubiquinol to complex III ^51^. In summary, the TPP data suggest an acute impairment of mitochondrial function for troglitazone, benzbromarone, amiodarone, and antimycin A, through major effects on CI and CIII.

### Metabolic perturbations upon DILI compound treatment confirm an acute MRC inhibition

We next assessed the functional state of mitochondria by measuring the mitochondrial membrane potential using a JC-10 fluorescence assay after 2 and 8h of compound treatment (Fig. 5a, Supplementary Fig. 12a, Supplementary Table 9). Loss of mitochondrial membrane potential (MMP (Ψm)) is indicative for bioenergetic stress as it reflects electron transport and oxidative phosphorylation processes. The complex III inhibitor antimycin A, troglitazone as well as benzbromarone already decreased the MMP (Ψ_m_) after 2h. CI inhibition with rotenone decreased the MMP only after 8h. No significant effects were observed for amiodarone and rosiglitazone (Fig. 5a, Supplementary Fig. 12a). These results suggest, that benzbromarone and troglitazone acutely induce mitochondrial dysfunction. Moreover, based on the rapid depolarization of the MMP upon antimycin A compared to rotenone, complex I inhibition alone cannot explain the strong depolarization observed with troglitazone.

In order to trace compound-induced mitochondrial dysfunctions back to distinct respiratory chain complexes we conducted untargeted cellular metabolomics experiments on dHepaRG cells treated with antimycin A, rotenone, troglitazone, benzbromarone, rosiglitazone, and amiodarone for 8 hours (Fig. 5a, Supplementary Fig. 12b-e). Inhibition of the respiratory chain and the resulting impairment of ATP generation via OXPHOS induced a metabolic shift to glycolysis^52^ as evident by elevated lactic acid levels and led to characteristic changes of TCA cycle metabolites (Fig. 5a, Supplementary Table 12b-c).

Electrons entering the MRC at complex I from NADH or complex II via succinic acid recycle NADH and FADH to NAD+ and FAD+ ^53^. Upon CI dysfunction, electrons from accumulating NADH are transferred to pyruvate, producing lactic acid, replenishing NAD+ and leading to decreased acetyl-CoA levels ^52–57^. We observed a moderate increase of lactic acid with antimycin A and rotenone, whilst benzbromarone and troglitazone led to high intracellular lactic acid levels. Rosiglitazone only slightly increased lactic acid (Fig. 5a). Acetyl-CoA levels decreased with all compounds except rosiglitazone. Together these results indicate a MRC inhibition at the level of complex I upon rotenone, benzbromarone and troglitazone.

CI inhibition with rotenone and accumulation of NADH inhibits TCA cycle enzymes and decreases succinic acid levels, demonstrating, that succinate stills enter the TCA cycle as long as CIII is functional ^58^. In contrast, CIII inhibition with antimycin A impairs FADH2/FAD^+^ recycling and the conversion of succinate to fumarate leading to succinic acid accumulation as reflected in our data. We observed increased succinic acid levels, notably with benzbromarone, rosiglitazone, and slightly with troglitazone, suggesting CIII impairment (Fig. 5a, Supplementary Fig. 12c). Moreover, all compounds, except for amiodarone, decreased citric acid, and benzbromarone, rosiglitazone and troglitazone decreased fumaric acid and malic-acid levels, further indicating strongly reduced TCA cycle activity and conversion of malate to pyruvate to increase glycolysis.

Moreover, acute energy depletion due to impaired OXPHOS was reflected in decreased ATP and phospho-creatine levels, observed with all compounds, most notably with benzbromarone and troglitazone (Fig. 5a, Supplementary Fig. 12d). Correspondingly, there was an increase in AMP and creatine with benzbromarone, troglitazone and rosiglitazone. (Supplementary Fig. 12d). Inhibition of CI and CIII is associated with ROS generation, evidenced by decreased levels of antioxidants glutathione and ascorbic acid with all compounds, indicating increased oxidative stress and further confirming respiratory chain inhibition (Supplementary Fig. 12e).

Overall, our findings confirm respiratory chain impairment, a switch to glycolysis and oxidative stress induced by the compounds.

### Mitochondrial protein release is an active process which is dependent on Rab7

To further explore the mechanism and export route of mitochondrial proteins upon drug treatment, we tested the effect of small molecule inhibitors targeting conventional and unconventional secretion on mitochondrial protein release upon benzbromarone treatment (Fig. 6a-g, Supplementary Table 11).

**Fig. 6.**
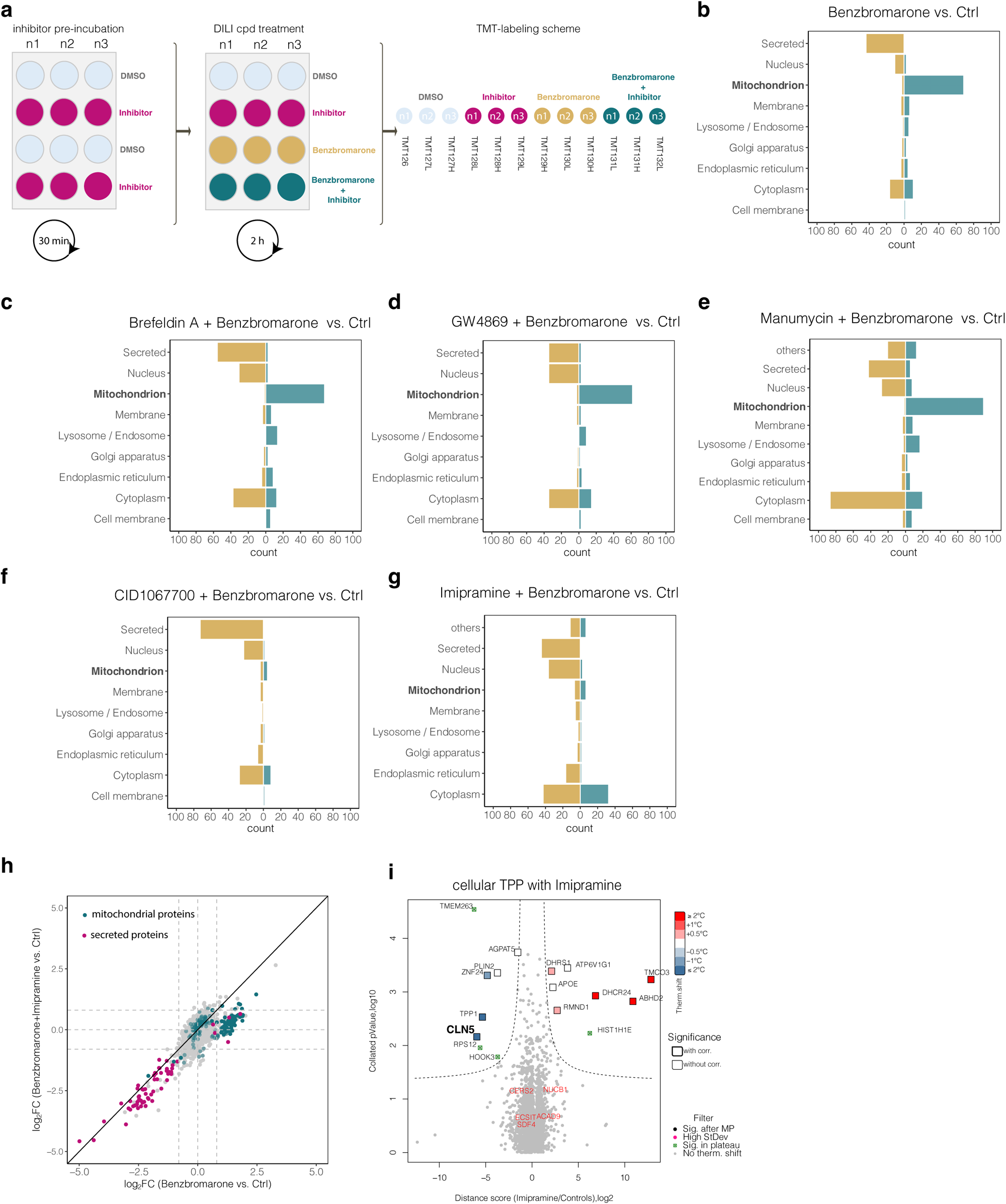
Mitochondrial protein release is an active process which is dependent on Rab7. **a**, Schematic illustration to test the effect of small molecule inhibitors targeting conventional and unconventional secretion on mitochondrial protein release upon benzbromarone treatment. Differentiated HepaRG cells in 12 well plates were treated with benzbromarone (25 µM), secretion inhibitor, the combination or DMSO as control. Tested inhibitors were brefeldin a (1 µM), GW4869 (500 nM), manumycin (500 nM), CID1067700 (25 µM) and imipramine (10 µM). dHepaRG cells were preincubated for 30 min with the inhibitors or DMSO as control, followed by the treatment with benzbromarone. Supernatants were harvested after 2h and the samples were combined into one sample for mass spectrometric analysis individually per inhibitor using TMT-12 isobaric mass tags. For each condition three independent biological replicates were used. **b**, Bar graph showing proteins significantly changing in the secretomes of benzbromarone versus DMSO treated dHepaRG cells grouped by their subcellular location annotation (based on UniProt annotation). Brown bars indicate significantly lower secreted proteins, green bars indicate a significant upregulation in protein secretion. Displayed are the counts for each subcellular location. c, same as b for the combined treatment of benzbromarone with brefeldin a. **d**, same as b for the combined treatment of benzbromarone with GW4869. **e**, same as b for the combined treatment of benzbromarone with manumycin. **f**, same as b for the combined treatment of benzbromarone with CID1067700. **g**, same as b for the combined treatment of benzbromarone with imipramine. **h**, Changes in benzbromarone dependent protein secretion patterns upon addition of imipramine after 2h treatment. X-axis displays log_2_ protein fold changes of benzbromarone treated dHepaRG cells versus time matched controls to identify benzbromarone dependent secretion. Y-axis displays log_2_ protein fold changes of benzbromarone, and imipramine treated dHepaRG cells versus time matched controls to identify inhibitor-dependent alterations. mitochondrial proteins are shown in green, secreted proteins are shown in purple. **i**, Protein thermal stability changes in dHepaRG cells upon treatment with 20 µM imipramine (90 min). Volcano plot displays distance scores and collated P values (-log10 transformed) of proteins quantified in imipramine (20 µM, n=2) versus DMSO-treated (n=2) dHepaRG cells. The ratio-based approach included LIMMA analysis for ratios between treatment and control groups obtained at each temperature, aggregation of retrieved P values per protein by Brown’s method and multiple testing adjustment Benjamini–Hochberg (BH) correction. Dotted line indicates significance cut-offs. Proteins passing significance cut-off are colored according to their Tm shift, bold edges indicate Benjamini-Hochberg corrected P-values with P< 0.05, light gray dots depict proteins that were not significantly affected.

Inhibition of the classical secretion pathway using brefeldin A, did not inhibit the benzbromarone induced secretion of mitochondrial proteins but further reduced basal secretion (Fig. 6c), suggesting the involvement of an unconventional secretion pathway for the release of mitochondrial proteins.

Inhibition of the ESCRT independent exosome pathway by GW4869 also did not affect the secretion of mitochondrial proteins (Fig 6d). The farnesyltransferase inhibitor manumycin, modulating ESCRT dependent exosome pathways, even increased levels of mitochondrial proteins in cell culture supernatants (Fig. 6e). Next, we hypothesized that the mitochondrial protein release might originate from the recently described mitochondria derived vesicles (MDV) which bud directly from the surface of mitochondria^59–64^. The small GTPase RAB7 has been found to be a critical factor for the transport of MDVs and indeed inhibition of RAB7 with the antagonist CID1067700 blocked release of mitochondrial proteins (Fig 6f).

Further, the CAD Imipramine has been found to block the release of microvesicles and exosomes by inhibition of acid sphingomyelinase (aSMase)^65^. and when applied in combination with benzbromarone a strong reduction of mitochondrial protein release was observed (Fig. 6g-h). In TPP we observed destabilization of CLN5 by imipramine, which was recently found to play a critical role in controlling the localization of Rab7 to the endosomal membrane and its activation ^66^ (Fig. 6i) consistent with a Rab7-dependent mechanism.

In summary these results demonstrate that the release of mitochondrial proteins by DILI compounds is an active and Rab7-dependent process induced by oxidative stress resulting from perturbations in respiratory complexes.

### DILI compounds impair basal protein secretion by induction of ER stress in dHepaRG cells

Next, we further investigated mechanisms leading to impaired protein secretion, which was observed for many compounds used in this study. Whilst DILI compounds such as troglitazone and perhexiline have been reported to interfere with the secretion of individual proteins such as APOB ^7, 67, 68^ our data set provides insights on a wide range of endogenously secreted proteins over a large panel of compounds. For many compounds (e.g. rotenone, troglitazone, or antimycin A) impairment of basal secretion was already detected 2h after compound addition (Fig. 7a) leading to reduced secretion of proteins belonging to the blood coagulation pathway, complement activation or acute-phase response in cell culture supernatants.

**Fig. 7.**
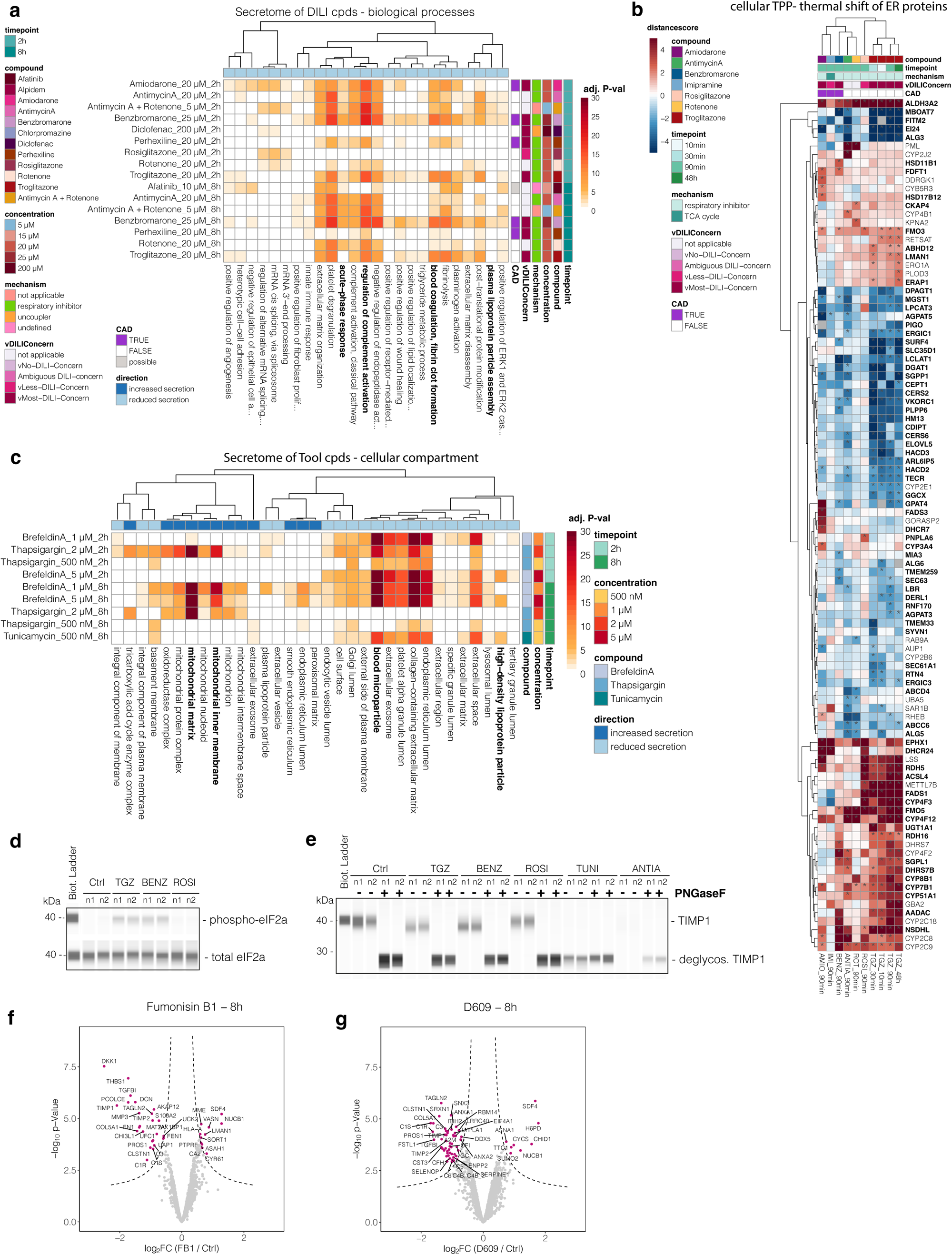
DILI compounds impair basal protein secretion by induction of ER stress in dHepaRG cells. **a**, DILI compounds acutely impair basal secretion in dHepaRG cells. Heatmap displaying significant GO-terms (biological processes) derived from the differential secretome analysis of DILI compound treated HepaRG cells (n=3) versus the time-matched control at 2h post stimulus (n=3) or 8h post stimulus (n =2). Differentially secreted proteins were determined via LIMMA. Significance thresholds were (p(Benjamini Hochberg) < 0.05 and log_2_ fold change > 2x standard deviation of the individual treatment. Color intensities indicate the adjusted (BH-corrected) p-value (-log_10_-transformed) of the GO-term. Only GO-terms are displayed that were significant upon three or more DILI compounds and were a reduced secretion could be observed upon compound treatment. Rows are clustered by Pearson correlation. **b**, DILI compounds change the thermal stability of ER membrane proteins. Heatmap displaying proteins of the ER with significant thermal stability shifts upon treatment of dHepaRG cells with different DILI compounds. Displayed are distance scores. Rows and columns are clustered by Euclidian distance. Statistically significant changes are denoted with asterisks (*). Bold gene names denote transmembrane proteins. **c**, ER stressors mimic DILI-compound induced secretion events. Heatmap displaying significant GO-terms (cellular component) derived from the differential secretome analysis upon treatment of HepaRG cells with different ER-stressors (n=3) versus the time-matched control at 2h post stimulus (n=3) or 8h post stimulus (n =2). Differentially secreted proteins were determined via LIMMA. Significance thresholds were (p(Benjamini Hochberg) < 0.05 and log_2_ fold change > 2x standard deviation of the individual treatment. Color intensities indicate the adjusted (BH-corrected) p-value (-log_10_-transformed) of the GO-term. Only GO-terms are displayed that were significant upon three or more DILI compounds and were an reduced secretion could be observed upon compound treatment. Rows are clustered by Pearson correlation. **d**, WES analysis of eIF2a phosphorylation and total eIF2a in dHepaRG cells lysates after 8h of treatment with troglitazone, benzbromarone and rosiglitazone. **e**, WES analysis of the glycoprotein TIMP1 in HepaRG cell lysates after 8h of treatment with troglitazone, benzbromarone, rosiglitazone, tunicamycin or antimycin a. Lysates were treated with PNGaseF for enzymatic deglycosylation and compared to their untreated counterparts to identify possible DILI-compound induced glycosylation defects. **f**, Volcano plot showing proteins quantified in the secretome of fumonisin b1 treated dHepaRG cells 8h post stimulus. Displayed are the log_2_ fold changes and the p-values (-log_10_-transformed) determined by LIMMA of fumonisin b1 treated dHepaRG cells (n=3) versus the time matched vehicle (DMSO)-controls (n=3). Dotted line indicates significance cut-offs. Proteins passing the significance thresholds (p(Benjamini Hochberg) < 0.05 and log_2_ fold change > 2x standard deviation of the individual treatment are colored in purple. **g**, same as f for the treatment of dHepaRG cells with the sphingomyelin synthase inhibitor D609 after 8h of treatment.

Previous reports indicated that ER stress is an important contributor to the pathogenesis of DILI by promoting the disruption of normal cellular functions and the hepatic metabolism ^7, 67, 69–71^. In our cellular TPP data we observed numerous thermal stability changes of ER proteins (Fig. 7b), e.g. upon treatment with troglitazone. Affected ER proteins were predominantly transmembrane proteins and mediate various pivotal functions (Fig. 7b) including protein glycosylation (e.g. ALG3, ALG6 and DPAGT1) and lipid and sphingolipid metabolism (e.g. SGPP1, CERS2, CERS6, HACD2 and SGPL1).

In addition, proteins involved in cargo transport and COPII coat complex proteins, such as LMAN1, SURF4, ERGIC1, SAR1B, RAB9A, SEC63, SEC61A1 showed significant thermal stability shifts. This led us to hypothesize that troglitazone and other compounds showing this pattern, such as antimycin A and benzbromarone, cause ER stress by compromising the ER and the secretory pathway leading to impaired protein secretion. Consistent with this, we found that the known ER stressors brefeldin A, thapsigargin and tunicamycin acutely impaired basal protein secretion (Fig. 7c, Supplementary Fig. 13a-k).

ER stress and the subsequent accumulation of proteins in the ER typically results in the induction of the unfolded protein response (UPR) via the three ER stress sensors PERK, ATF6 and IRE1α ^72–74^. PERK activation is the first and immediate reaction to ER stress leading to phosphorylation of eukaryotic translation initiation factor 2 subunit-α (eIF2α), and consequently attenuation of protein synthesis ^75^.

We found that troglitazone and benzbromarone but not rosiglitazone induced the phosphorylation of the eukaryotic translation initiation factor-2α (eIF2α) after 8 h treatment, whilst the total level of eIF2α was unaltered (Fig. 7d) which is consistent with detected secretion patterns.

Thapsigargin induces ER stress through ER-Ca^2+^ store depletion which is known to induce the secretion of ER-resident chaperones ^76^. We observed secretion of the co-chaperones DNJAB11 and DNJB9 after 8h upon mild Ca^2+^ store depletion with 500 nM thapsigargin whilst strong Ca^2+^ depletion with 2 µM thapsigargin additionally induced the secretion of the chaperones HSP90B1 and HSPA5 (GRP78/BiP). Interestingly, enhanced secretion of chaperones such as HSP90B1, HSPA5, CALR and DNAJB11 was also observed upon treatment with brefeldin A or the DILI compound perhexiline, suggesting that these compounds also induce Ca^2+^ store depletion from the ER (Supplementary Fig. 13a, Supplementary Table 3). Moreover, 500 nM thapsigargin induced the secretion of the Golgi resident protein NUCB1, whereas 2 µM did not lead to NUCB1 secretion (Fig. 1b, Supplementary Table 3a, 3c), most likely because of Golgi fragmentation caused by severe ER-Ca^2+^ store depletion ^77^. Interestingly, acute NUCB1 secretion was reported upon treatment with chemical stressors suggesting to be essential for ATF6 dependent UPR activation ^78^.

Of note, thapsigargin (2 µM) and brefeldin A (1 µM and 5 µM), induced the secretion of mitochondrial proteins after 2h and 8h, suggesting that massive Ca^2+^ store depletion stimulates mitochondrial stress leading to MDV release (Fig. 7c, Supplementary Fig. 13b-i, Supplementary Table 3a, 3c).

An alternative mechanism to induce ER stress is the inhibition of protein glycosylation. To test for this, dHepaRG cells were treated for 8h with the DILI compounds troglitazone, benzbromarone, rosiglitazone, antimycin A or tunicamycin which blocks the first committed step in the protein N-glycosylation pathway ^79^. We found that only tunicamycin fully blocked the glycosylation of TIMP1 whilst troglitazone and benzbromarone only had minor effects on the molecular weight of TIMP1, indicating little or no impact on glycosylation (Fig. 7e). In summary these data suggest that DILI compounds like troglitazone and benzbromarone acutely impair ER function and induce ER stress in dHepaRG cells, by a mechanism which is not dependent on common mechanisms such as inhibition of the protein glycosylation or Ca^2+^ store depletion but is correlated with widespread effects on protein stability of ER membrane proteins.

### Secretion of the Golgi resident protein SDF4 informs altered sphingolipid metabolism

A recurring pattern for a subset of DILI compounds, including troglitazone, benzbromarone, stavudine, nimesulide, perhexiline, fluoxetine, was the secretion of SDF4 already after 2h (Fig. 1b-f, Supplementary Table 3a, 3c). SDF4 is involved in intra-Golgi trafficking- and trans-Golgi-cargo sorting processes of soluble cargo proteins ^80, 81^. The sorting of soluble secretory proteins requires active sphingomyelin biosynthesis which links Ca^2+^ influx into the trans-Golgi network (TGN) lumen to Ca^2+^ dependent SDF4 oligomerization and subsequent capture of secretory cargo proteins in the TGN ^80, 82–90^.

The precursor of sphingomyelin is ceramide and depletion or disturbance of ceramide synthesis and the transport machinery in the ER are known to interfere with cargo sorting processes and Golgi Ca^2+^ homeostasis ^80, 91^. As such we speculated that these pathways could also be impaired upon DILI compound treatment, leading to dysregulation of sphingomyelin synthesis in the Golgi and subsequent secretion of SDF4. Consistent with this hypothesis we found that proteins involved in sphingolipid- and ceramide metabolism, such as CERS2, CERS6, SGPL1 and SGPP1 showed significant thermal shifts in TPP experiments with troglitazone and benzbromarone (Fig. 7b). Further we observed elevated intracellular phospho-ethanolamine levels in metabolomics analyses providing further evidence for impaired ceramide metabolism (Supplemental Fig. 13l). To test, whether an inhibition of the ceramide metabolism has an impact on the TGN, stimulating the release of SDF4, we treated the cells with different concentrations of the ceramide synthase inhibitor fumonisin b1 (FB1) and analyzed the secretomes (Fig. 7f). FB1 treatment stimulated the release of SDF4 together with NUCB1 and concomitantly interfered with the release of secreted proteins like TIMP1, which is a known SDF4 cargo protein ^88, 90, 92^. To further narrow down the mechanism for compound specific SDF4 secretion, we treated the cells with the sphingomyelin synthase (SMS) inhibitor D609 (Fig. 7g, (Supplementary Fig. 13m). Inhibition of SMS and thus, inhibition of Ca^2+^ influx into the TGN, likewise stimulated the release of SDF4 and NUCB1 and simultaneously impaired protein secretion, including the release of TIMP1 (Fig 7g).

In summary these data demonstrate that secreted SDF4 is a biomarker for molecules that interfere with ceramide and sphingomyelin synthesis pathways.

## Discussion

Profiling of the extracellular proteome of the HepaRG liver cell model in response to treatment with a total of 46 DILI compounds revealed widespread effects including modulation of conventional secretion but also non-canonical secretion including vesicle mediated processes. Using thermal proteome profiling and a number of validation assays, we showed that the altered extracellular proteome is correlated with dysregulation of a diverse range of mechanisms in distinct organelles.

### Secretomics and thermal proteome profiling of lysosomotropic compounds

Amongst those compounds that accumulate reversibly in specific organelles due to their physicochemical properties ^36, 41^, cationic amphiphilic drugs (CADs) are of particular interest in the drug discovery process, as excessive drug accumulation in the lysosome disturbs lysosomal catabolism and can result in drug-induced phospholipidosis (DIPL) ^37, 93^. The exact cellular mechanism relating to how structurally diverse CADs induce DIPL is still not well understood. It is unclear if CADs directly bind lysosomal proteins and thereby inhibiting their function or if the lysosomal dysfunction is the result of an indirect effect e.g. by compensating the negative surface charge of intralumenal vesicles (ILVs) which eliminates the electrostatic binding of lysosomal enzymes to the ILV surface ^37, 40^.

We observed thermal destabilization of soluble lysosomal proteins upon treatment with the CAD amiodarone for 2 h which suggests indirect effect rather than direct binding to all these proteins. Consistent with this, experiments with the selective V-ATPase inhibitor bafilomycin A1 revealed that thermal destabilization of luminal lysosomal proteins is associated with increased lysosomal pH, leading to the release of lysosomal proteins into the cell culture supernatant.

Moreover, we observed thermal stabilization of cytoplasmic V-ATPase subunits (ATP6V1C1, ATP6V1D, and ATP6V1E1) upon amiodarone treatment whereas the V-ATPase inhibitor bafilomycin A1 stabilized subunits of the catalytic membrane-embedded V0 portion of this protein ^45^. Stabilization of V1 subunits was also observed with the unrelated compounds antimycin A, rotenone, and troglitazone (Supplementary Table 4, Supplementary Fig. 5), for which no lysosomal protein release was observed. This suggests that thermal stabilization of V1 subunits does not necessarily indicate inhibition of the V-ATPase but may rather indicate an increase in the activity of this proton pump, e.g. to counteract the CAD-induced alkalinization of the lysosomal lumen.

All together these data show, that lysosomal dysfunction of the CADs tested in this study is not caused by direct binding to lysosomal proteins rather than by miss targeting of lysosomal proteins to the extracellular space because of changes in the lysosomal pH.

The extent of lysosomal trapping and concurrent lysosomal protein release trends with drug basicity as exemplified by the two tyrosine kinase inhibitors afatinib (pKa: 8.81) and lapatinib (pKa: 7.2) or the class III antiarrhytmic drug amiodarone (pKa 8.47). Of note, the stronger lysosomotropic effect of afatinib in our data is in line with lysosomal trapping predictions ^42^ and could also explain the longer half-life of afatinib of 37 h compared to lapatinib, as up to 50 % of the intracellular drug can be sequestered in lysosomes ^42^ and is only gradually released to bind its intended target the EGF receptor. This underlines how the distribution of a drug within highly compartmentalized mammalian cells affects both, its activity and pharmacokinetics.

### Inhibition of the mitochondrial respiratory chain induces secretion of mitochondrial proteins

Most compounds investigated in this study are known mitochondrial toxicants, and our data link mitochondrial protein release with acute mitochondrial dysfunction and oxidative stress. We found that the release of mitochondrial proteins is an active process, dependent on the small GTPase Rab7 and facilitated by a stress-induced burst in MDVs that act as a first line of defense by removing damaged or oxidized proteins and remodeling the mitochondrial proteome before mitophagy occurs ^59, 60, 94, 95^.

Consistent with a prior study characterizing MDV protein content from isolated mitochondria under oxidative stress^62^ , proteins enriched in the secretome were mainly derived from the mitochondrial matrix, including enzymes harboring hyper-reactive cysteine residues and iron-sulfur clusters (Supplementary Fig. 3c, Supplementary Fig. 10c) which are susceptible to oxidative damage. The small GTPase Rab7 was shown to be involved in stress-induced MDV delivery to lysosomes and multivesicular bodies from where they can be either degraded in the lysosomes or shuffled to the extracellular space via EVs ^59, 96, 97^. Our data provide the first report of MDV formation caused by mitotoxic DILI compounds in hepatocyte models.

Intriguingly, the physiological impact of mitochondrial content in EVs and the extracellular space and-most importantly in the context of idiosyncratic DILI - is still elusive. Discharge of dysfunctional oxidized mitochondrial proteins could act as prevention mechanism to reduce potential proteotoxic stress within the cell, especially under conditions where lysosomal degradation could reach its limits, and to allow in-vivo clearance by patrolling macrophages. T-cells have been identified in liver biopsies of patients with DILI, suggesting an active involvement of the immune system. Interestingly, Rab7 dependent MDVs are known to mediate mitochondrial antigen presentation upon infection ^96, 98, 99^. In the context of idiosyncratic DILI, both the release of mitochondrial content and antigen presentation on the surface of hepatocytes could be important drivers for immune mediated and late onset idiosyncratic DILI.

### DILI compounds that induce mitochondrial secretion acutely destabilize proteins of the inner mitochondrial membrane including the respiratory complexes I and III

Based on the thermal proteome data obtained in this study we formed a hypothesis on the underlying molecular mode-of-action of different tool and DILI compounds on the respiratory chain complexes I and III ^21, 100, 101^. We found that the reference CIII inhibitor antimycin A that directly binds to the Qi-site ^23^ formed by MT-CYB induced thermal stabilization of the respective CIII subunits. In contrast, the DILI compounds rosiglitazone and troglitazone showed different mechanisms of action. Besides PPARγ-mediated transcriptional activity, studies suggest the antidiabetic action of glitazones may result from acute inhibition of CI and CIII^102^. Our data are consistent with these reports but further suggest different molecular mechanisms by which these glitazones inhibit the respiratory chain and exert their antidiabetic properties. Increased succinic acid levels and strong thermal stabilization identified DHODH as a likely direct rosiglitazone off-target. DHODH links nucleotide metabolism and mitochondrial function via CIII and further, DHODH inhibitors were found to impair respiration and increase glycolysis explaining rosiglitazone’s blood glucose lowering effect ^51, 103–105^. Notably, DHODH inhibitors have been proposed as anticancer treatments and also for rosiglitazone an antitumor effect was observed ^104, 106^. However, the anticancer effect of rosiglitazone has not been fully elucidated yet and it is tempting to speculate, that the antitumor effect of rosiglitazone may be mediated via inhibition of DHODH.

Unlike rosiglitazone’s distinct effect on DHODH, troglitazone destabilized multiple CI and CIII subunits, indicating an indirect mechanism. Time-dependent TPP experiments with troglitazone (Supplementary Fig. 11a) revealed an initial thermal destabilization of membrane-spanning and proton-transporting ND4- and ND2 module subunits, followed by destabilization of CI matrix arm subunits at later timepoints. Thus, we conclude, that the initiating events happen in the membrane part of the complex leading to structural disassembly of CI and release of these subunits by MDVs into the extracellular space. This hypothesis is further corroborated by the increased abundances of multiple N-module and Q-module subunits (e.g. NDUFS4, NDUFV2, NDUFS1) in cell culture supernatants (Supplementary Fig. 11c). Consistent with our thermal stability data upon troglitazone treatment, structural destabilization leading to complex I disassembly has been observed for the related pioglitazone ^107^.

Besides destabilizing of CI subunits, troglitazone caused thermal destabilization of CIII matrix subunits (UQCRC1, UQCRC2, UQCRFS1) within 10 minutes. Interestingly, the CI subunit NDUFB4 was shown to interact with CIII subunit UQCRC1, crucial for respirasome integrity and mitochondrial function and molecular dynamics simulations indicate that structural changes in UQCRC1 can propagate throughout complex III, affecting its function. Hence our data suggest that impairment of CI integrity is transmitted via NDUFB4 into CIII leading to disruption of the respiratory supercomplexes and mitochondrial stress (Fig. 5c, Supplementary Fig. 11d). Taken together, an initiating event in the CI – membrane interface is observed, which affects the membrane protein interaction of CI and progresses to CIII leading to thermal destabilization and acute impairment of these respiratory chain complexes (Fig. 5c, Supplementary Fig 11d).

Notably, similar secretion patterns containing mitochondrial CI matrix arm subunits were observed for a number of other compounds including benzbromarone and perhexiline suggesting that structural CI disassembly is a common consequence of structurally unrelated DILI compounds.

### DILI compounds impair basal protein secretion by induction of ER stress in dHepaRG cells

Another common observation for multiple DILI compounds was impaired basal secretion. This is consistent with previous studies reporting reduced secretion of individual proteins, such as APOB, with a limited set of compounds, including troglitazone ^7, 67^. However, our data clearly demonstrate that drugs can have broader and more far-reaching effects on cellular protein secretion than previously shown (Fig. 1b, Fig. 7a).

We used the ER-stressors brefeldin A, thapsigargin and tunicamycin to further investigate how drug effects can lead to impaired basal secretion. Our data disprove ER-stress induction by DILI compounds via inhibition of the protein N-glycosylation pathway but instead suggest that ER stress arises from impaired protein folding and export.

Both, brefeldin A and thapsigargin induced chaperone secretion, suggesting Ca^2+^ store depletion, as chaperones are Ca^2+^-binding proteins and their interactions as well as their retention in the ER are dependent on high luminal Ca^2+^ concentrations^76, 108, 109^. These observations are consistent with reports of reduced ER-Ca^2+^ levels upon brefeldin A treatment^110^. Among all DILI compounds tested, only the known ER stressor perhexiline triggered significant secretion of the chaperones HSPA5 and HSP90B1, consistent with ER Ca^2+^ store depletion as the underlying mechanism.

For troglitazone and benzbromarone, however, we demonstrated that ER stress arises from compromised ER protein export as evidenced by significant thermal stability changes of the cargo receptors SURF4 and LMAN1 (Fig. 7b). Both cargo receptors were previously found to regulate the export of soluble proteins from the ER, participate in ER exit site organization ^111–114^ and the structural maintenance of the ERGIC ^115^. SURF4 was identified as the relevant cargo receptor for a broad repertoire of cargo proteins including lipoproteins like APOB ^116, 117^. These proteins were found reduced in cell culture supernatants by a range of DILI compounds consistent with impaired SURF4/LMAN1 function thus explaining the development of drug induced steatosis.

The presence of mitochondrial proteins in the cell culture supernatant upon thapsigargin and brefeldin A treatment further indicates a functional connection between ER stress and mitochondrial dysfunction. Discharge of ER-Ca^2+^ stores is known to trigger Ca^2+^ loading of mitochondria ^118^, which has immediate effects on mitochondrial metabolism ^119^. Mitochondrial Ca^2+^ levels stimulate the TCA cycle and OXPHOS, promoting increased ATP synthesis; however, increased OXPHOS activity results in increased MRC electron leakage, ROS levels, and mitochondrial stress, and can ultimately induce cell death by opening the mitochondrial permeability transition pore (mPTP) ^118, 120^.

The Golgi-resident protein NUCB1 was found to be a negative feedback regulator in the ATF6-mediated branch of the UPR ^78^ and acute NUCB1 secretion was reported upon treatment with chemical stressors suggesting to be essential for UPR activation by facilitating ATF6 cleavage in the Golgi ^78^. In our secretome data (Fig. 1b, Supplementary Table 3a, 3c), the ER stressor thapsigargin and a subset of DILI compounds including troglitazone and benzbromarone induced secretion of NUCB1 after just 2h of incubation further corroborating that DILI compounds induce acute ER stress and establishing NUCB1 is an ER stress informing marker protein for the ATF6-branch of the UPR in hepatocyte cell models.

Furthermore, a partially overlapping subset of DILI compounds, including troglitazone, induced the secretion of the Golgi resident protein SDF4 which is involved in cargo transport- and sorting-processes in the trans-Golgi network ^80, 81, 86^. SDF4 retention in the TGN requires high Ca^2+^ concentrations in the Golgi lumen.

Our data suggests, that SDF4 can be used as diagnostic marker protein to inform on different perturbations and pathways within the cell: SDF4 secretion consistently coincided with destabilization of components of the sphingomyelin and ceramide biosynthesis pathways (Supplementary Fig. 13l), accumulation of phosphoethanolamine and stabilization of SGPL1. These data cumulatively propose dysregulation of sphingomyelin and ceramide biosynthesis as causal for SDF4 secretion which was confirmed by blocking ceramide- and sphingomyelin synthase with specific inhibitors. This is further corroborated by a previous study, which has reported an inhibition of the de-novo ceramide synthesis in skeletal muscle cells upon troglitazone ^121^. Proteins whose secretion was impaired upon treatment with D609 or fumonisin B1 are likely to be SDF4 cargo proteins, such as the previously identified protein TIMP1 ^122^.

Hence, our data establish SDF4 as a mechanism informing marker for perturbations of ceramide- or sphingomyelin biosynthesis in hepatocyte cell models.

### DILI compounds induce off-target effects by non-specific perturbations in lipid bilayers

In our TPP data we found that DILI compounds affected thermal stabilities of a multitude of structurally and functionally unrelated TM proteins in both live cells and crude cell lysates (Fig. 3). Intriguingly, the majority of proteins were destabilized and mapped to those organelles from which proteins observed in the secretomes originated (Fig. 3e-f). Further, thermal stability changes were concentration dependent, in the case of troglitazone showed immediate onset (within 10 min), were likely induced by the parent compound rather than by compound metabolites and were not a result of secondary compound effects as many TM proteins were also destabilized in crude cell extracts and not only in live cells. Moreover, HepG2 cells that exhibit low CYP activity and are thus metabolically inactive induced similar secretion patterns upon troglitazone, benzbromarone and rosiglitazone treatments (Supplementary Fig. 14, Supplementary Table 13), further corroborating that the observed effects are induced by the parent compound. Consistent with our observations, many of the reported off-target proteins of e.g. troglitazone are TM proteins ^123^. In TPP, direct binding of a drug to a protein stabilizes the protein such that it becomes more resistant to heat-induced unfolding (higher melting temperature), whilst a destabilization is often associated with lowered structural stability (lower melting temperature) which can e.g. occur when a protein complex is disturbed ^124^. Hence, the multitude of structurally and functionally unrelated TM proteins that were destabilized upon DILI compound treatment cannot be easily explained by the classic concept of pharmacological regulation that assumes direct drug binding to the protein, which would mean that drugs with totally different chemistries modulate the same off-targets.

The classical model of drug-target interactions tends to ignore the critical role of lipid bilayers, which closely interact with transmembrane (TM) proteins. TM proteins are energetically coupled to the lipid bilayer they reside in, and the composition and properties of lipid bilayers directly regulate protein structure and function^125^. In previous work, we also demonstrated that membrane proteins are stabilized in the lipid bilayer with melting points being correlated with the number of transmembrane domains ^27^. Therefore, transitions in TM protein conformation can lead to changes in adjacent lipid organization and vice versa^126^. Drug-induced disruptions of lipid metabolic pathways, such as those affecting the sphingomyelin pathway, can rapidly alter organelle lipid compositions, thus affecting bilayer properties as well as protein stability and function.

Our data suggest that DILI drugs like troglitazone or benzbromarone act through a combination of mechanisms, including direct drug-protein interactions and indirect, non-specific interactions via the lipid bilayer. Many compounds used in this study are lipophilic or amphiphilic (Supplementary Fig. 1a), allowing them to interact with lipid bilayers either by partitioning into the bilayer-water interface^127, 128^ or by increasing drug concentrations near the membrane^129^. These interactions can modify membrane protein function by altering bilayer properties such as thickness, curvature, and fluidity^126, 128, 130^. Depending on physicochemical properties like logP, drug concentrations in lipid bilayers can be much higher than in the cytoplasm, reaching millimolar concentrations. For example, the higher logP of troglitazone (5.1) facilitates higher potential drug concentrations in lipid bilayers compared to rosiglitazone (3.1) an analogue with much lower DILI concern, potentially explaining observed differences in affected TM proteins and secretion patterns between these related compounds.

Previous work using membrane model systems for mitochondria^131–133^ suggested that NSAIDS have a certain preference for membranes with distinct lipid compositions. Our data, for the first time demonstrates such a segregation for a wide range of compounds in live cells across a number of intracellular organelles, including inner mitochondrial and ER membranes.

Organelle membranes of eukaryotic cells differ in their lipid composition, thickness and physical properties ^126, 134, 135^ ^136–138^, e.g. ER membranes are thinner than the plasma membrane ^134^ ^139^. Consequently, protein transmembrane domains differ in amino acid compositions as well as lengths across organelles. Rusinova et al. ^140^, demonstrated by using artificial vesicle membranes, that drugs can directly alter the function of the proton gated ion channel KcsA through accumulation in the lipid bilayer. Moreover, drug effects on the KcsA activity were dependent on bilayer thickness. Our data show that these processes also translate in-cellulo and are mainly driven by the physicochemical properties of a drug, impacting the activity of a multitude of different TM proteins in certain organellar membranes.

Thus, lipid composition and bilayer thickness and the physicochemical properties of the drug, e.g. lipophilicity and sterical effects, determine to which extent a compound adsorbs to or partitions into a membrane, thus explaining the organellar-specific effects observed in the ER and inner mitochondrial membrane. A compelling example in our data are the structurally related glitazones troglitazone and rosiglitazone, showing that troglitazone which has a higher lipophilicity than rosiglitazone, has the most widespread effects.

In summary, our data provide evidence that regulatory effects of drugs on TM protein function by alterations of membrane properties , which have been observed in artificial cell membrane models ^123, 132, 140, 141^, also translate into in-cellulo biological activity with immediate impact on organellar physiology.

Here, we presented a deep pharmacoproteomic profiling of protein secretion and protein thermal stability of small molecule perturbation processes as exemplified with 46 DILI and non-DILI compounds, annotated for mitochondrial mechanisms and 8 tool compounds. We have systematically investigated the acute effects of drugs and tool compounds on the extracellular proteome of dHepaRG cells and provide a resource for acute secretion changes upon small molecule perturbations. Unlike previous studies that have focused solely on drug effects on protein expression ^1, 142^ and modifications ^2^, our pharmacoproteomics approach reveals additional, often overlooked dimensions of drug action, offering a deeper and more comprehensive understanding. We demonstrated that the observed biology in the secretome can be linked to specific events in cellular organelles and show, that drug effects are much broader than anticipated. We suggest novel ways of modulation that explain the toxicity mechanisms of DILI compounds and demonstrated that the pleiotropy of DILI compounds results in widespread effects accumulating in distinct cellular organelles. Investigating secretion changes upon drug treatment is particularly interesting as it identifies diagnostic tools for delineating drug-induced dysregulation that can potentially translate in vivo informing both on mechanism of dysregulation but also on the potential for inflammatory response. This could be crucial for understanding drug safety and idiosyncratic DILI, especially if hepatocyte priming is already present, such as from prior inflammation.

In conclusion, our study demonstrates that multidimensional pharmacoproteomics profiling integrating secretomics and thermal protein profiling provides unique insights into drug-induced perturbations and can provide diagnostic marker proteins for preclinical studies to elucidate potential drug toxicity mechanisms.

## Abbreviations

ADRs: Adverse drug reactions
CADs: cationic amphiphilic drugs
dHepaRG: differentiated HepaRG cells
DILI: drug-induced liver injury
DIPL: drug-induced phospholipidosis
MDV: mitochondria derived vesicles
MoA: mechanism of action
RLU: Relative Light Unit
TGN: trans-Golgi network
TM: transmembrane
TPP: thermal proteome profiling
MMP: mitochondrial membrane potential
MRC: mitochondrial respiratory chain
MS: mass spectrometry

## Data availability

All data is available in the main text or the supplementary materials. The mass spectrometry proteomics data have been deposited to the ProteomeXchange Consortium via the PRIDE ^143^ partner repository with the dataset identifiers PXD052568, PXD052601, PXD052609 (https://www.ebi.ac.uk/pride).

## Code availability

The code used for this study will be made available by the corresponding authors upon reasonable request.

## Acknowledgements

We thank J. Stuhlfauth, N. Garcia-Altrieth, K. Beß and B. Dlugosch for supporting the cell culture; M. Klös-Hudak, K. Kammerer, P. Sauer and M. Steidel for technical assistance with mass spectrometry, C. Boecker for IT and computational support as well as B. Heller for scientific discussion.

## Author contributions

M.B., B.K. and H.C.E. conceived and supervised the project. S.K. performed experiments. S.K., H.C.E., M.K., J.K., R. K., M. Z. S., D.C.S., and M.B. analyzed data and S.K. and T.M. curated the data. S.K., H.C.E., and M.B. wrote the manuscript with input from all authors.

## Materials and Methods

### Reagents

Acetaminophen, Acetylsalicylic acid, Afatinib, Alendronate, Alpidem, Ambrisentan, Amiodarone, Benzbromarone, Betaine, Busulfan, Chloroquine, Chlorpromazine, Clotrimazole, Clozapine, Cromoglicic acid, Diclofenac, Disulfiram, Entacapone, Fluoxetine, Flutamide, Folic acid, Haloperidol, Imipramine, Lapatinib, Leflunomide, Lumiracoxib, Metformin, Methotrexate, Nefazodone, Nimesulide, Nitrofurantoin, Oxybutynin, Pargyline, Perhexiline, Pioglitazone, Piroxicam, Propranolol, Rosiglitazone, Rotenone, Simvastatin, Stavudine, Tamoxifen, Tetracycline, Troglitazone, Zafirlukast were obtained from a GSK internal compound library as 10 mM compound stocks in DMSO. Rotenone (3616), Brefeldin A (1231), Thapsigargin (1138), Bafilomycin A1 (1334), Tunicamycin (3516), FumonisinB1 (3103) and D609 (1437) were purchased from Tocris Bioscience (United Kingdom). Exosome inhibitors CID1067700 (SML0545), GW4869 (D1692), Manumycin A (M6418) and Complex III inhibitor Antimycin A (A8674) were purchased from Merck (Darmstadt, Germany). Human phospho-eIF2α (Ser51) Antibody (#9721) was purchased from Cell Signaling Technologies. Human total eIF2α Antibody (#AF3997-SP) and human TIMP-1 antibody (#AF970-SP) were purchased from Tocris (Biotechne, United Kingdom). PNGaseF (V4831) was purchased from Promega (Mannheim, Germany).

### Cell culture

Undifferentiated human HepaRG cells (HPR101056, Biopredic International, Saint-Gregoire, France) were cultured and differentiated according to the supplier’s instructions. Briefly, cryopreserved cells were thawed and maintained in William’s E-medium (Thermo Fisher) supplemented with HepaRG Growth medium with antibiotics (ADD710, Biopredic International). Growth medium was exchanged twice a week. After two weeks in HepaRG Growth medium, fully confluent HepaRG cells were trypsinized and passaged to maintain the cell line or seeded into 12-well plates with a density of 2.7 x 10^4^ cells/cm^2^. After 2 weeks of proliferation in 12-well plates HepaRG growth medium was replaced by differentiation medium with antibiotics (ADD720, Biopredic International), which causes the cells to differentiate to hepatocyte colonies and primitive biliary cells within 2 weeks. Differentiated HepaRG cells were maintained in differentiation medium until use for a maximum of 4 further weeks. Human hepatoma cell line HepG2 (obtained from ATCC)) was maintained in MEM medium (Thermo Fisher) supplemented with 10% FCS, 2 mM glutamine, 1 % non-essential amino acids and 1 mM sodium pyruvate. All cell lines were maintained at 37°C in a humidified atmosphere of 5 % CO_2_. If not otherwise stated, all experiments used differentiated HepaRG cells in 12 well plates as starting material.

### Cell treatments and preparation of secretome samples

All secretomics experiments were conducted as biological triplicates in 12 well plates. For the interval-based secretomics ^15^ screening experiment, differentiation medium was exchanged with fresh differentiation medium containing the respective compound as indicated in the text or differentiation medium only. The DILI compound screen was conducted at two concentration regimens per compound. Compounds were screened, if not otherwise stated, at 2 or 20 µM end concentration. Benzbromarone was always used at 25 µM end concentration. Acetaminophen and Diclofenac were used at 30 and 300 µM. HepaRG cells were incubated for 6h at 37 °C in an atmosphere of 5 % CO_2_. After 6h, the differentiation medium was removed, and the cells were washed carefully three times with serum-and phenol red-free Williams E-medium. Cells were incubated for an additional two hours in serum-and phenol red-free Williams E-medium with the respective compound concentrations or with DMSO as vehicle control. Final DMSO concentration was set to 1.7% and the final medium volume was set to 1 mL per well. Cell supernatants from all samples were carefully collected and 230 µL supernatant was transferred into a 0.45 µm 96-well filter-plate (Durapore, low protein-binding PVDF membrane, Merck Millipore) to remove detached cells and cell debris. Supernatants were cleared by centrifugation (2 minutes at 100 x g) and collected in a 96 well polypropylene plate. 200 µL cleared supernatants were then transferred into a fresh 96 well polypropylene plate and subsequently sealed and stored at -80 °C.

For secretomics experiments with inhibition of protein secretion pathways, the medium was exchanged with fresh differentiation medium containing either Brefeldin A, GW4869, Manumycin, CID1067700, Imipramine or were left untreated. HepaRG cells were pre-incubated in differentiation medium with the inhibitors or DMSO as control for 30 min at 37 °C in an atmosphere of 5 % CO_2_. After this preincubation period, the medium was removed, and the cells were carefully washed three times with prewarmed Williams E-medium. Samples were either mock treated (DMSO) as control, 25 µM Benzbromarone alone, inhibitor or the combination of inhibitor and 25 µM Benzbromarone. The final DMSO concentration was set to 1.7 %. All treatments were performed in phenol red- and serum-free Williams E-medium, with 1 mL final volume per well. Cells were incubated for 21h at 37°C in a humidified atmosphere of 5 % CO2. After 2 h, cell supernatants were carefully collected and processed as described above. Secretion inhibitors were used at the following concentrations: Brefeldin A was used at 1 µM, and GW4869 and Manumycin A were used at 500 nM, CID1067700 was used at 25 µM and Imipramine was used at 10 µM.

### LDH-assay

Cell viability was assessed by determination of LDH release into cell culture supernatants using the LDH-GLO cytotoxicity assay (Promega, Mannheim, Germany) following the manufacturer’s instructions. The maximal LDH activity was determined by addition of 20 µL 10 % Triton X-100 to the wells of the control cells.

### Enzymatic lactic acid assay

Lactate in the cell culture supernatants was measured with the Lactate-Glo assay (Promega, Mannheim) according to the manufacturer’s instructions. Nimesulide samples were lost during sample preparation.

### JC-10 assay for measurement of the mitochondrial membrane potential

HepaRG cells were seeded in black 96 well plates (Thermo Scientific, # 10281092) with a density of 0.009 x 10^6^ cells per well following the differentiation procedure as described above. dHepaRG cells were then treated with compounds at the indicated timepoints. Mitochondrial membrane potential was measured using a microplate JC-10 Assay Kit (Abcam, #ab112134). For determination of the mitochondrial membrane potential after 2h of compound treatment, cells were incubated in Williams E medium containing the compounds or DMSO as control and the JC-10 dye. JC-10 dye was used with the final concentration described by the manufacturer. In case of the 8h timepoint cells were, analogue to the interval-based secretomics procedure, first treated in differentiation medium containing the indicated compounds or DMSO as control. After 6h, medium was shifted to serum-free conditions using Williams E medium containing the indicated compounds or DMSO as control and the JC-10 assay. Cells were incubated for additional 2h. All pipetting steps were performed using a VIAFLO 96 automatic pipetting robot (Integra Biosciences). All treatments were performed with four biological replicates. Fluorescence was measured as described by the manufacturer using an EnVision Multimode Plate Reader (PerkinElmer).

### LysoTracker Green DND-26 staining for live cell imaging of lysosomes

HepG2 cells were seeded in µ-Slide 8 Well high Glass Bottom slides (IBIDI, #80807) with a density of 15.000 cells per well. When the cells have reached a confluence of 80 %, the culture medium was carefully removed and 100 µL fresh culture medium containing 500 nM bafilomycin A1 or DMSO as control was added. Cells were incubated for 1.5 or 7.5 h at 37°C in a humidified atmosphere of 5 % CO2. After this 100 µL pre-warmed culture medium containing 20 µg/mL DAPI (ThermoFisher, #62248) for nuclear staining and 100 nM Lysotracker Green DND-26 (Invitrogen, # 11594976) for staining of acidic organelles, was added to each well, and cells were incubated for additional 30 min at 37°C in a humidified atmosphere of 5 % CO2. Live cell imaging was performed with an APX100 Benchtop Fluorescence Microscope (Evident) equipped with a Xylis fluorescence illuminator and operated with the Olympus imaging software cellSens (version 4.1). Exposure times for each individual experiment were determined first for the brightest sample, avoiding overexposure, and remained constant throughout the imaging procedure.

### Capillary western blot (WES) analysis of cell culture supernatants

Frozen secretome samples were dried in vacuo. Pellets were resuspended in 40 µL sample buffer (100 mM TRIS-HCl, 125 mM Trizma base, 10 % glycerol, 2% SDS, 0.005 bromphenol blue, 50 mM DTT) and incubated for 15 minutes at room temperature on an orbital shaker (Heidolph Titramax 1000) at 800 rpm. Afterwards samples were transferred into a 96well PCR-plate,and heated for 5min at 95°C in an PCR cycler. Capillary Western analyses were performed according to the manufacturer’s instructions using the ProteinSimple Wes System. Briefly, 4 µL of the denatured sample were combined with 1 µL 5 × Fluorescent Master Mix (containing 5 × sample buffer, 5 × fluorescent standard, and 2001mm DTT) in the designated wells of a 12-230 kDa separation module (cat.no. # SM-W001). Afterwards the prepared reagents were dispensed step-by-step into the designated wells of the separation module. A biotinylated ladder provided molecular weight standards for each assay. For the detection of human TIMP1, the antibody (# AF970-SP) was used with the recommended end concentration of 50 µg/mL. After plate loading, the separation electrophoresis and immunodetection steps, via chemo luminescence readout, take place in the fully automated capillary system.

### Preparation of cell lysates for WES analysis

After preparation of the cell culture supernatants for secretomics analysis, cells were lysed directly in the 12well plate using a SDS-based-lysis buffer (50 mM TRIS-HCl (pH 7.4), 1.25% SDS, 1x C0mplete EDTA free protease inhibitor (Roche Mannheim), 1x PhosStop (Roche, Mannheim), 1mM MgCl_2_, 2U/µL Benzonase (Merck, #E1014-25KU)). Briefly, cells were washed twice with warm DPBS. Residual DPBS was thoroughly aspirated. 50 µL of the prepared lysis buffer was added to each well, distributed with a cell scraper and afterwards transferred into a 1.5 mL Eppendorf tube. Lysates were heated for 3 min to 95°C in thermomixer at 900 rpm. Afterwards, lysates were cooled down to room temperature and 0.48 µL benzonase (Merck, #E1014-25KU) was added to each sample. Samples were incubated for 30 min at 37°C and 900 rpm in a thermomixer. Lysates were cleared through centrifugation for 20 min at 20,000 xg at room temperature and cleared lysates were then transferred into fresh Eppendorf tubes. Protein concentration was determined with an BCA-assay (Pierce, #23225) and set to 2 µg/µL with dilution buffer (50 mM TRIS-HCL (pH 7.4), 50 mM DTT). Samples were stored until use at -80 °C.

### PNGaseF digest of dHepaRG cell lysates

Cell lysates were thawed to room temperature and 1 % IGEPAL CA-630 was added to each sample. Afterwards, 10 ng PNGaseF (Biotechne, #9109-GH-020) were added to each sample, following incubation for 1 h at 37 °C in a thermomixer (800 rpm). PNGaseF treatment was repeated by adding 5ng PNGaseF to each sample following incubation for additional 30 min in a thermomixer. Samples were then heated for 5 min to 95°C to inactivate PNGaseF.

### Capillary western blot (WES) analysis of cell lysates

Cell lysates and PNGaseF digested samples were transferred to a 96well PCR-plate, diluted to a protein concentration of 0.8 µg/µL using 0.1 x WES sample buffer (ProteinSimple) and heated to 95°C for 5 min in a PCR cycler. Capillary Western analyses were performed as described above using a 12-230 kDa separation module (cat.no. # SM-W001). For the detection of human TIMP1, the antibody (# AF970-SP) was used with the recommended end concentration of 50 µg/mL. For the detection of total human eIF2a (Biotechne, # AF3997-SP) the antibody was used at 10 µg/mL. For the detection of phospho-eIF2a (Cell Signaling Technologies, # 9721S), the antibody was diluted 1:50.

### Preparation of crude cell extracts

HepaRG cells were pelleted by centrifugation and resuspended in one pellet-volume of ice-cold DPBS. Cell suspension was transferred to pre-cooled 0.5 mL microtubes pre-filled with 1.4 mm ceramic beads (Omni International, # 19626). Cells were dissociated with the Omni Bead Ruptor (Omni International) pre-cooled to 4°C and operated with the following settings: Speed was set to 4 m/s; cycle time was set to 0.10 s; number of cycles was set to 1 and the dwell time was set to 0. One pellet volume of ice-cold DPBS containing 3 mM MgCl_2_ was added followed by benzonase (Merck, #E1014-25KU) in a 1:1000 ratio. The crude lysate was incubated for 1h on a rotating shaker at 4°C and the protein concentration was determined using a BCA-assay. Crude lysate was then diluted to 15 mg/mL and distributed into 100 µL aliquots, snap frozen in liquid nitrogen and stored until use at -80°C.

### TPP-TR experiments

#### TPP-TR experiments in cells

TPP-TR was performed as described in ^24^. In brief, dHepaRG cells in 12 well plates were either treated with a compound or an equivalent amount of DMSO (vehicle) was added. Treatments were performed in HepaRG differentiation medium. Every treatment was performed in duplicates using 6 wells for each replicate. Incubation of cells with compound and vehicle was conducted in parallel for the indicated durations at 37°C and 5% CO_2_. Cells were collected by trypsination and for each biological replicate, 6 wells from the same treatment were pooled. Cells were washed three times with DPBS carefully and then resuspended in 1200 μl DPBS. 100 μl of this cell suspension was transferred into 0.2-ml PCR plates and subjected to a centrifugation step at 400 xg for 3 min at RT. Then, 80 μl of the PBS supernatant was removed using a VIAFLO 96 automatic pipetting robot (Integra Biosciences). Samples were then heated in parallel in a PCR cycler for 3 min to a temperature range of 37 to 66.3 °C, followed by a 3-min incubation time at room temperature. IGEPAL CA-630 (I3021) was added to a concentration of 0.8%, MgCl2 (M1028) to 1.5 mM and benzonase to 1 kU/mL, and samples were incubated for 1 h at 4 °C and 750 rpm on an orbital shaker. Cell lysates were transferred to a 0.45 µm 96-well filter-plate (Durapore, low protein-binding PVDF membrane, Merck Millipore), centrifuged for 5 min at 500 xg and 4°C to remove aggregates and collected in a 96 well polypropylene plate. 30 µL of the filtered lysate were transferred into a fresh 96 well plate and dried in vacuo. The residual filtered lysate was used to determine the protein concentrations with a BCA-assay (Pierce, #23225). Afterwards, dried down lysates were reconstituted in 30 µL sample buffer (100 mM TRIS-HCl, 125 mM Trizma base, 10 % glycerol, 2% SDS, 0.005 bromphenol blue, 50 mM DTT) and stored at -80 °C.

#### TPP-TR experiments in crude cell extracts

Crude extracts at 0.3 mg/mL protein concentration were incubated with compound or vehicle (DMSO, 0.5% end concentration) for 15 min at 25 °C, then 10 wells of a PCR plate were filled with 100 μl of each reaction. Samples were subsequently heated in parallel in a PCR cycler for 3 min to a temperature range of 37 to 66.3 °C, followed by a 3 min incubation time at room temperature. IGEPAL CA-630 (I3021) was added to a concentration of 0.8 %, MgCl2 (M1028) to 1.5 mM and benzonase to 1 kU/mL, and samples were incubated for 1 h at 4 °C and 750 rpm on an orbital shaker. Aggregated proteins were separated by ultracentrifugation (30 min, 100,000 xg and 4°C). 60 µL were carefully taken off from each ultra-centrifugation tube and transferred into a 0.2 µm 96-well filter-plate (Durapore, low protein-binding PVDF membrane, Merck Millipore), centrifuged for 5 min at 500 xg and 4°C and the flow through was collected in a 96 well polypropylene plate. 50 µL of the filtered lysate was then transferred into a fresh 96 well polypropylene plate, dried down in vacuo and reconstituted in 25 µL sample buffer (100 mM TRIS-HCl, 125 mM Trizma base, 10 % glycerol, 2% SDS, 0.005 bromphenol blue, 50 mM DTT) and stored at -80 °C. Samples were further processed for LC–MS/MS analysis.

### Extraction of polar metabolites for untargeted metabolomics analysis

dHepaRG cells in 12well plates were treated with compounds as described for the interval-secretomics experiments. After the incubation time, medium was thoroughly aspirated and cells were washed two times with 2 mL pre-warmed wash solution (75 mM Ammonium bicarbonate, pH 7.4). Subsequently, cellular metabolism was quenched by snap-freezing the cells in liquid nitrogen. Plates were stored until use at -80°C. Polar metabolites were extracted with -20°C cold 80 % acetone. 0.5 mL pre-cooled acetone were added to each well without detaching the cells. Plates were handled on wet ice throughout the procedure. After addition of acetone plates were stored for 2h at -20 °C. The supernatant, which contains the polar metabolite fraction, was carefully transferred to a 96well deep-well plate and subsequently dried in vacuo. Dried metabolites were resuspended in 60 µL 30 % AcN and transferred to a 0.45 µm 96-well filter plate. Plates were centrifuged for 2 min at 2000 xg and the filtrate was collected in a 96-well polypropylene plate and subjected to MS analysis.

### Untargeted metabolomics analysis by mass spectrometry

Untargeted metabolomics analysis was performed using a platform modeled on a previously described platform ^145^. Briefly, samples were analyzed on a LC-MS platform consisting of a Thermo Scientific Ultimate 3000 liquid chromatography system with autosampler temperature set to 10°C coupled to a Thermo Scientific Q-Exactive Plus Fourier transform mass spectrometer equipped with a heated electrospray ion source and operated in negative. For negative ionization mode, the isocratic flow rate was 250 μL/min of mobile phase consisting of 60:40% (v/v) isopropanol: water buffered with 11mM ammonium fluoride at pH 9 containing 10 nM taurocholic acid and 20 nM homotaurine as lock masses. Mass spectra were recorded in profile mode from 50 to 10001m/z with the following instrument settings: sheath gas, 151a.u.; aux gas, 5 a.u.; aux gas heater, 400°C; sweep gas, 0 a.u.; spray voltage, −3 kV; capillary temperature, 250° C; S-lens RF level, 50 a.u; resolution, 701k @ 2001m/z; AGC target, 3x106 ions, max. Inject time, 120 ms; acquisition duration, 60 s. Spectral data processing including peak detection and alignment was performed using an automated pipeline in R that was designed analogously to the previously published MATLAB version ^145^. In brief, the pipeline aligns and averages scans of each individual file, then aligns spectra across all files in the dataset, performs centroiding and peak binning, re-calibrates the mass axis based on the above lock masses and finally assembles a consolidated matrix of ion intensities. Detected ions were tentatively annotated as metabolites based on matching accurate mass within a tolerance of 5 mDa using the Human Metabolome database ^146^ as reference considering [M-H] and [M-2H] (negative mode) ions and up to two 12C to 13C substitutions. Importantly, this approach does not enable to conclusively identify metabolites, as for instance metabolites with identical chemical formula, with different adducts or within the 5 mDa annotation window can map to the same ion. When interpreting our data, we therefore made sure to be as conservative as possible, focusing on metabolites that are known to be highly abundant or part of conserved metabolic pathways. Also, in cases where annotations of multiple ions responding to a change in feeding strategy mapped to a shared pathway, we favored the parsimonious explanation that this shared pathway was affected over alternative and less likely scenarios invoking effects on multiple distant pathways. We therefore made sure to interpret our data conservatively and focus on most likely annotations, for instance known highly abundant metabolites in conserved metabolic pathways.

### Secretome sample preparation for Mass Spectrometry

Frozen secretome samples were dried in vacuo. Pellets were resuspended in 115 µL resuspension buffer (2 % SDS, 0.5 % IGEPAL CA-630) and incubated for 15 minutes at room temperature on an orbital shaker (Heidolph Titramax 1000) at 800 rpm. 100 µL resuspended samples were processed through a modified version of the single pot solid-phase sample preparation (SP3) protocol as described in ref ^147^. Briefly, proteins in resuspension buffer were bound to paramagnetic beads (SeraMag Speed beads, GE Healthcare, CAT#45152105050250, CAT#651521050502) by addition of 160 µl cleanup solution (130 µL ethanol, 27.5 µL 15 % formic acid, 2.5 µL bead slurry) to a final ethanol concentration of 50 %. Beads were washed 4 times with 200 µL 70 % ethanol. Proteins were digested by resuspending in 40 µL 0.1 mM HEPES (pH 8.5) containing 1.25 mM TCEP, 5 mM chloroacetamide, 5 ng/µL trypsin, and 5 ng/µL LysC following o/n incubation. Peptides were labeled with isobaric mass tags (TMT-11 or TMTpro-16, Thermo Fisher Scientific). The labeling reaction was performed in 100 mM HEPES (pH 8.5) 50 % acetonitrile at 22°C and quenched with 2.5 % hydroxylamine. Labeled peptide extracts were pooled and purified using C18SCX stage-tips as described in ref ^147^. TMT-labeled samples were fractionated into 8 fractions prior to LC-MS/MS analysis using the high pH reversed-phase peptide fractionation kit (Thermo Fisher Scientific), following the manufacturer’s instructions.

### TPP sample preparation for Mass Spectrometry

All samples were processed through a modified version of the single pot solid-phase sample preparation (SP3) protocol ^148, 149^ as described in ref ^147^. Briefly, proteins in 2% SDS were bound to paramagnetic beads (SeraMag Speed beads, GE Healthcare, CAT#45152105050250, CAT#651521050502) by addition of ethanol to a final concentration of 50%. Contaminants were removed by washing 4 times with 70% ethanol. Proteins were digested by resuspending in 40 µL 0.1 mM HEPES (pH 8.5) containing 1.25 mM TCEP, 5 mM chloroacetamide, 5 ng/µL trypsin, and 5 ng/µL LysC following o/n incubation. Derived peptides were subjected to TMT labeling using TMTpro reagents, enabling relative quantification of 16 conditions in a single experiment ^150^. The labeling reaction was performed in 100 mM HEPES (pH 8.5) 50% Acetonitrile at 22 °C and quenched with hydroxylamine. Labeled peptide extracts were combined to a single sample per experiment by combining 8 temperatures of the control samples and 8 temperatures of compound treated samples, as described in Fig. 3a. The labeled samples were offline pre-fractionated using high pH reversed phase chromatography ^151^ into 24 individual fractions prior to LC-MS/MS using an Ultimate3000 RLSC (Dionex).

### LC-MS/MS analysis of proteomics samples

Fractionated and lyophilized samples were resuspended in 0.05% trifluoroacetic acid in water and 30 % of each sample was injected into an Ultimate3000 nanoRLSC (Dionex) coupled to a Q Exactive (Thermo Fisher Scientific) or an Orbitrap Fusion Lumos mass spectrometer (Thermo Fisher Scientific). Peptides were separated on custom-made 50 cm × 100 μm (ID) reversed-phase columns (C18, 1.9 μm, Reprosil-Pur, Dr. Maisch) at 55 °C. Gradient elution was performed from 2 % acetonitrile to 40 % acetonitrile in 0.1 % formic acid and 3.5 % DMSO over 65 min at a flow rate of 350 nL/min. Samples were online injected into the mass spectrometer. The Q Exactive Plus was operated in a data-dependent top 10 acquisition method. MS spectra were acquired using 70.000 resolution and an ion target of 31×110^6^ for MS1 scans. Higher energy collisional dissociation (HCD) scans were performed with 35 % NCE at 35.000 resolution (at m/z 200), and ion target setting was set to 21×110^5^ so as to avoid coalescence ^152^. The instruments were operated with Tune 2.4 and Xcalibur 3.0 build 63. The Orbitrap Fusion Lumos operated with a fixed cycle time of 3 s. MS1 spectra were acquired at a resolution of 60,000 and an ion target of 41×110^5^. HCD fragmentation was performed at 38 % NCE at a resolution of 30,000 and an ion target 11×110^5^. The instrument was operated with Tune v.2.1.1565.23 and Xcalibur v.4.0.27.10.

### Peptide and Protein Identification

Raw data were processed using an in-house pipeline based on the isobar quant package ^25^. Mascot 2.5 (Matrix Science, Boston, MA) was used for protein identification. In a first search 30 ppm peptide precursor mass and 30 mDa (HCD) mass tolerance for fragment ions was used for recalibration according to Cox et al. ^153^ followed by search using a 10 ppm mass tolerance for peptide precursors and 20 mDa (HCD) mass tolerance for fragment ions. Enzyme specificity was set to trypsin with up to three missed cleavages. The search database consisted of the SwissProt sequence database (SwissProt Human release December 2018, 42 423 sequences) combined with a decoy version of this database created using scripts supplied by Matrix Science. Carbamidomethylation of cysteine residues and TMT modification of lysine residues were set as fixed modification. Methionine oxidation, and N-terminal acetylation of proteins, and TMT modification of peptide N-termini were set as variable modifications.

Unless stated otherwise, we accepted protein identifications as follows. (i) For single-spectrum to sequence assignments, we required this assignment to be the best match and a minimum Mascot score of 31 and a 10× difference of this assignment over the next best assignment. Based on these criteria, the decoy search results indicated <1% false discovery rate (FDR). (ii) For multiple spectrum-to-sequence assignments and using the same parameters, the decoy search results indicate <0.1 % FDR. All identified proteins were quantified; FDR for quantified proteins was below 1 %.

### Peptide and Protein quantification

Reporter ion intensities were read from raw data and multiplied with ion accumulation times (the unit is milliseconds) so as to yield a measure proportional to the number of ions; this measure is referred to as ion area ^154^. Spectra matching to peptides were filtered according to the following criteria: mascot ion score >15, signal-to-background of the precursor ion >4, and signal-to-interference >0.5 ^155^. Fold-changes were corrected for isotope purity as described and adjusted for interference caused by co-eluting nearly isobaric peaks as estimated by the signal-to-interference measure ^156^. Protein quantification was derived from individual spectra matching to distinct peptides by using a sum-based bootstrap algorithm; 95% confidence intervals were calculated for all protein fold-changes that were quantified with more than three spectra ^154^. Only proteins quantified with more than one quantified spectrum (qusm) and more than one unique peptide (qupm) were considered for downstream analysis.

### Data analysis

All protein annotations were based on the UniProtKB database (April 4, 2022). Data analysis and visualizations were performed in R (version 4.0.2; www.r-project.org). All experiments were performed using a TMT labeling strategy.

In all secretomics experiments, identified proteins were filtered for qusm >1 and qupm >1. Secretomics time course experiments covering treatment times of 2h and 8h, each biological replicate of compound treatment and the corresponding time-matched controls for each of the two timepoints were combined into one multiplexed TMT-11 experiment. Briefly, for 2h timepoints, three independent biological replicates of treatments and vehicle control were included into the TMT-11 experiment. The residual 5 TMT channels were used for the 8h timepoints, made up of two independent biological replicates of the controls and three independent biological replicates of the treatments. For the secretomics compound screen, two of the three biological replicates of each compound and each concentration together with 2 biological replicates of time matched controls and two biological replicates of benzbromarone treated cells (served as positive control) were combined into a TMTpro-16 experiment, resulting in 15 single TMTpro experiments (Fig. 1b). Only samples were included that featured LDH-values which were as high as in their time matched controls and did not indicate cell death. For the inhibition of cellular secretion pathways with the small molecule inhibitors Brefeldin A, GW4869, Manumycin A, CID1067700 or imipramine, three biological replicates of DMSO treated control samples and three biological replicates of each treatment (benzbromarone only, inhibitor only or benzbromarone + inhibitor) were combined into one sample for mass spectrometric analysis per inhibitor using TMTpro-12 isobaric mass tags (Fig 6a).

In all secretomics experiments, data analysis was performed as described in reference ^15^. Briefly, summed up ion areas of each TMT-channel and each replicate were log_2_ transformed and normalized to the median of the density maxima of all samples. The normalization factors of each TMT-channel and each replicate were used to calculate a median of normalization factors for every TMT channel across the replicate measurements. A TMT-channel was identified as an outlier, if 1) the normalization factor of this individual TMT-channel was greater than 2 or 2) the median normalization factor of the triplicates from this treatment was greater than 1.5 due to this individual TMT-channel. Statistical analysis was done separately for each timepoint and each treatment by contrasting treatment to the time-matched control. All calculations were performed using LIMMA ^157^. Proteins were considered as significant when passing the following cut-offs: 1.) 2-times the median standard deviation of ratios between replicates across all tested proteins of each timepoint and treatment and 2.) by an adjusted p-value cut-off of 0.05 using Benjamini-Hochberg approach as described in ^27^. Gene ontology enrichments were obtained using the topGO package in R ^158^. The enriched GO-Terms were filtered, if not otherwise indicated, for Benjamini-Hochberg corrected p-values of < 0.05.

Cellular and crude lysate TPP data analysis were performed according to reference ^27^ using the ratio-based thermal shift assay approach (RTSA). Briefly, treatments and controls were combined in the same TMT experiment. This was done for 8 different temperatures resulting in two TMTpro-16 experiments with the reference temperature 37 °C in each TMTpro-16 experiment. These experiments were done in two biological replicates. Protein fold changes were calculated relative to the 37 °C vehicle-treated sample. These fold changes represent the relative amount of nondenatured protein at the corresponding temperature. The data were filtered for quantified proteins with qusm ≥ 3 and qupm ≥ 2. Vehicle-treated samples were normalized at each temperature as follows. First, proteins fulfilling the following criteria were selected for the calculation of the normalization factors: (1) quantified with qusm ≥ 3 and qupm ≥ 2 and (2) an abundance of ≥20% relative to the 37 °C vehicle sample. The distribution of the fold change was generated for each replicate and temperature. A normalization factor per temperature was calculated as the mean of the fold change at the respective distribution maximum from all replicate. All protein fold changes of vehicle-treated samples in the respective experiments were corrected with these normalization factors. Second, the fold changes of the compound-treated sample were normalized at each temperature as follows: proteins fulfilling following criteria were selected for the calculation of the normalization factors: (1) quantified with qusm ≥ 3 and qupm ≥ 2 and (2) an abundance of ≥20% relative to the 37 °C vehicle sample. The distributions of the fold changes were generated at each replicate and temperature. A normalization factor per temperature and replicate was calculated as the difference between the fold change at the distribution maximum of the compound-treated and the vehicle-treated groups. These normalization factors were then applied to all protein fold changes of the compound-treated condition. The normalized datasets of both TMTpro-16 experiments covering all temperature points were then combined. Based on this data, for each protein three effect measures including test statistics were determined: (1) a distance score that represents a sensitive measure to detect treatment-induced effects, (2) an abundance score indicating treatment-induced abundance changes measured at the reference temperature and (3) the thermal stability difference between treatments and controls by fitting sigmoidal denaturation curves and estimating melting point shifts. In case of the distance score, we started by calculating two parameters for each protein at each temperature: the mean abundance ratio between treatment and control, and the significance of an abundance difference at the respective temperature using LIMMA based on log_2_ transformed treatment to control ratios. If the protein relative fold change in a replicate at a temperature was <0.15 in the vehicle-treated and the compound-treated condition, the log_2_ ratio at the specific temperature in that replicate was removed. As effects are often expected on multiple neighboring temperatures, neighboring data points can be combined to further increase statistical power. Here the most significant window of three P values at successive temperatures was selected and the P values were combined by the empirical Brown’s method ^159^ (collated P value). For the covariance matrix calculation, the log_2_ ratios of each replicate and each P value was used. A protein abundance ratio was used as second effect size to evaluate if the treatment resulted in a significant effect. If the P value of the above described LIMMA analysis on the log_2_ ratios were smaller than 0.05 at several consecutive temperatures, these log_2_ ratios were summed (collated log_2_ ratio). If several independent ‘blocks’ could be assigned, the one with the highest summed ratio was selected. If a protein showed no or only one P < 0.05, the average log_2_ ratio at the lowest P value was reported. Collated ratios were finally normalized to distance scores according to the following equation:

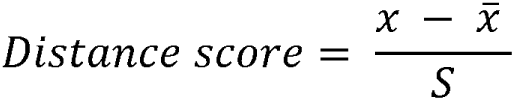

where x is the collated rato of the respectve protein, x̄ is the mean of collated ratos and S the standard deviation of collated ratios. The second effect measure, the abundance score, was determined by normalizing observed ratios between treatments and controls at reference temperature as described for distance scores. In the case of the third effect measure, melting curves and melting points were calculated based on fold changes relative to reference temperature as described above and LIMMA based significances of differences in the melting points between treatments and controls were calculated. After determining these effect measures, data were filtered for proteins fulfilling following criteria: median R^2^ of denaturation curve fits in control or treatment ≥0.8, average relative fold change in treatment or control of ≤0.4 at the highest temperature, qusm ≥ 3 and qupm ≥ 2 in at least two replicates (either in the first or second gradient part). Proteins fulfilling the following criteria were regarded as significantly affected by the treatment: (1) cumulative significance of protein abundance between treatments and controls fulfills the criteria:

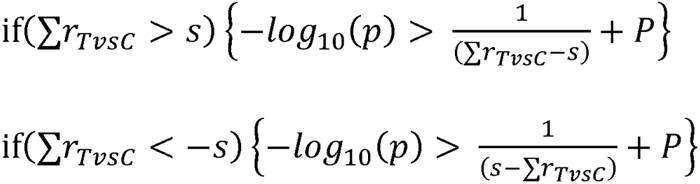

where 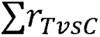 is the cumulative log2 ratio between treatments and controls (or log2 ratio at the most significant temperature), s the standard deviation of the dataset defined as 2× median standard deviation of ratios for all proteins between replicates and P the minimal accepted log10 transformed P value set to P = 0.05 and (2) absolute cumulative log2 ratio between treatments and controls 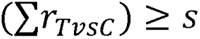.

To reduce overestimations of significances of protein abundance differences at high temperatures, proteins—for which most (≥50%) temperatures, selected for ratio collation, were in the melting curve plateau (denatured)—were excluded. The temperature at which the melting curve plateau starts, was defined as follows: the minimal residual protein abundance of the melting curve fit at 100 °C plus 10%. For proteins showing a significant distance score, two treatment-dependent effects could be distinguished based on the additional two effect measures: cell surface abundance changes and thermal stability changes. Proteins with a significant (P < 0.01) and an absolute relative fold change of ≥10% at 37°C in treatments compared to controls were regarded as significantly internalized/surface presented. Proteins with a significant (P < 0.05) and an absolute thermal shift of ≥0.5 °C in treatments compared to controls were regarded as significantly thermally shifted. Proteins that were significant after Benjamini– Hochberg correction of collated P values were highlighted by bold boxes in the plots. A value of 5% was selected as cut-off. The Benjamini–Hochberg correction was calculated with twice the number of proteins, since two P values were calculated per temperature and protein.

