## Supplementary Figures for "Pharmacoproteomic profiling identifies secreted markers for aberrant drug action"

**Supplementary Figure 1 |** Enzymatic measurements of lactic acid in the cell culture supernatants of dHepaRG cells.
**Supplementary Figure 2 |** DILI compounds acutely modulate the secretome of dHepaRG cells.
**Supplementary Figure 3 |** GO term enrichments of secretomes after treatment with DILI compounds for 2 and 8h.
**Supplementary Fig. 4 |** Secretion patterns of DILI compounds correlate with secretion changes induced by tool-compounds.
**Supplementary Figure 5 |** DILI compounds induce widespread protein thermal stability changes in dHepaRG cells.
**Supplementary Figure 6 |** Thermal proteome profiling confirms organelle specific effects of DILI compounds.
**Supplementary Figure 7 |** Effect of DILI compounds on membrane spanning proteins.
**Supplementary Figure 8 |** Secretomics and thermal proteome profiling of cationic amphiphilic drugs (CADs).
**Supplementary Figure 9 |** Drug-induced toxicity on mitochondria.
**Supplementary Figure 10 |** Benzbromarone and Troglitazone induce the secretion of a subnetwork of mitochondrial proteins into the cell culture supernatant.
**Supplementary Figure 11 |** DILI compounds that induce mitochondrial secretion acutely destabilize proteins of the respiratory complexes I and III.
**Supplementary Figure 12** | Metabolic perturbations upon DILI compound treatment confirm an acute MRC inhibition.
**Supplementary Figure 13 |** ER stressors mimic DILI-compound induced secretion events.
**Supplementary Figure 14 |** Metabolically inactive HepG2 cells induce similar secretion patterns upon troglitazone, benzbromarone and rosiglitazone treatments.

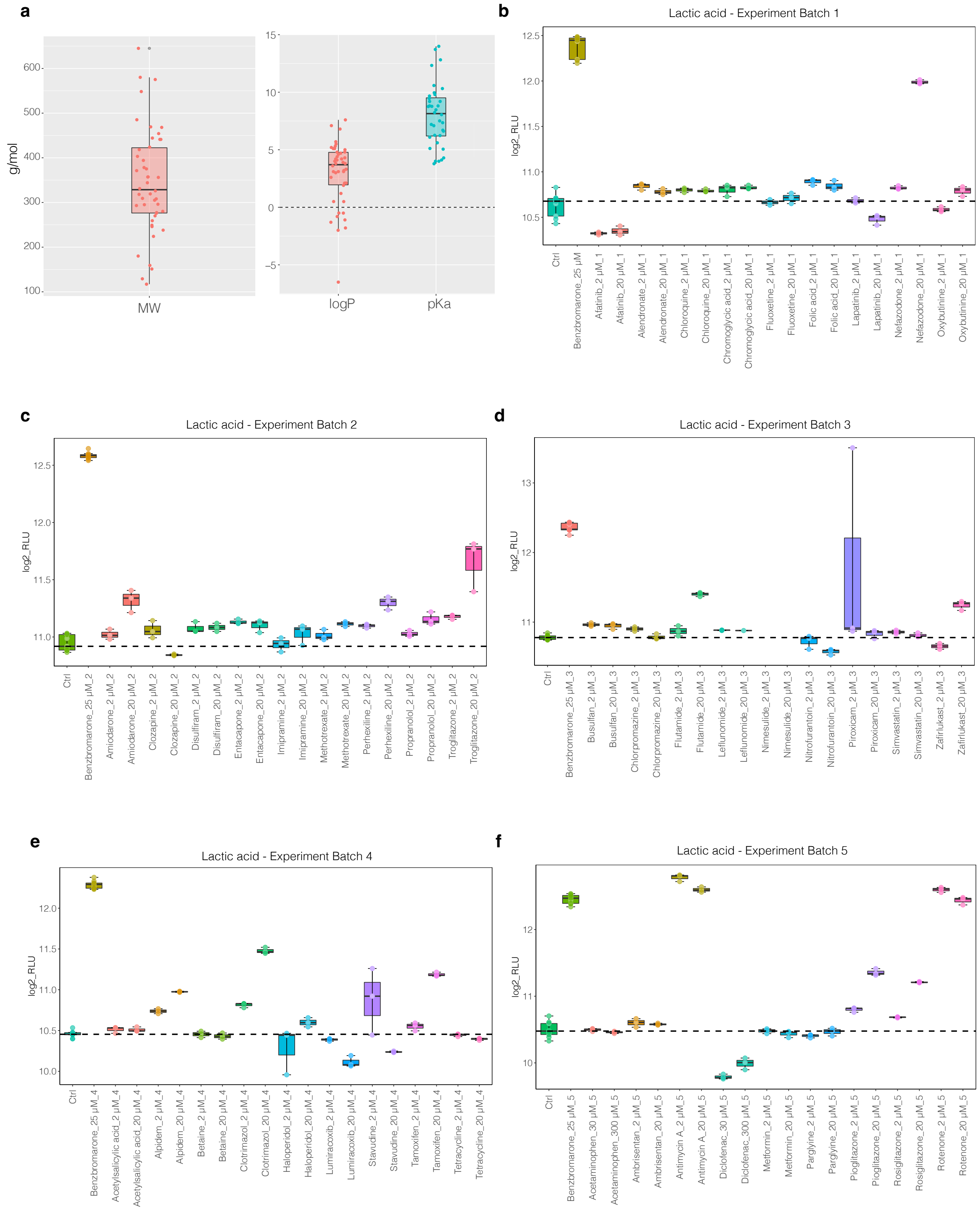


**Supplementary Figure 1| Enzymatic measurements of lactic acid in the cell culture supernatants of dHepaRG cells.**

**a,** Boxplots showing the molecular weight distributions (left panel) and logP and pKa distributions (right panel) for all tested compounds. **b,** Boxplots depict log2 RLU of lactic acid after 8h of compound treatment from experiment batch 1. Center line of each box, median; box limits, upper and lower quartiles; whiskers, maximum and minimum value of the dataset. Dotted line, median of the control samples. Data points denote log2 RLU derived from each of the three independent biological replicates. **c,** same as b for experiment batch 2. **d,** same as b for experiment batch 3. **e,** same as b for experiment batch 4. **f,** same as b for experiment batch 5.

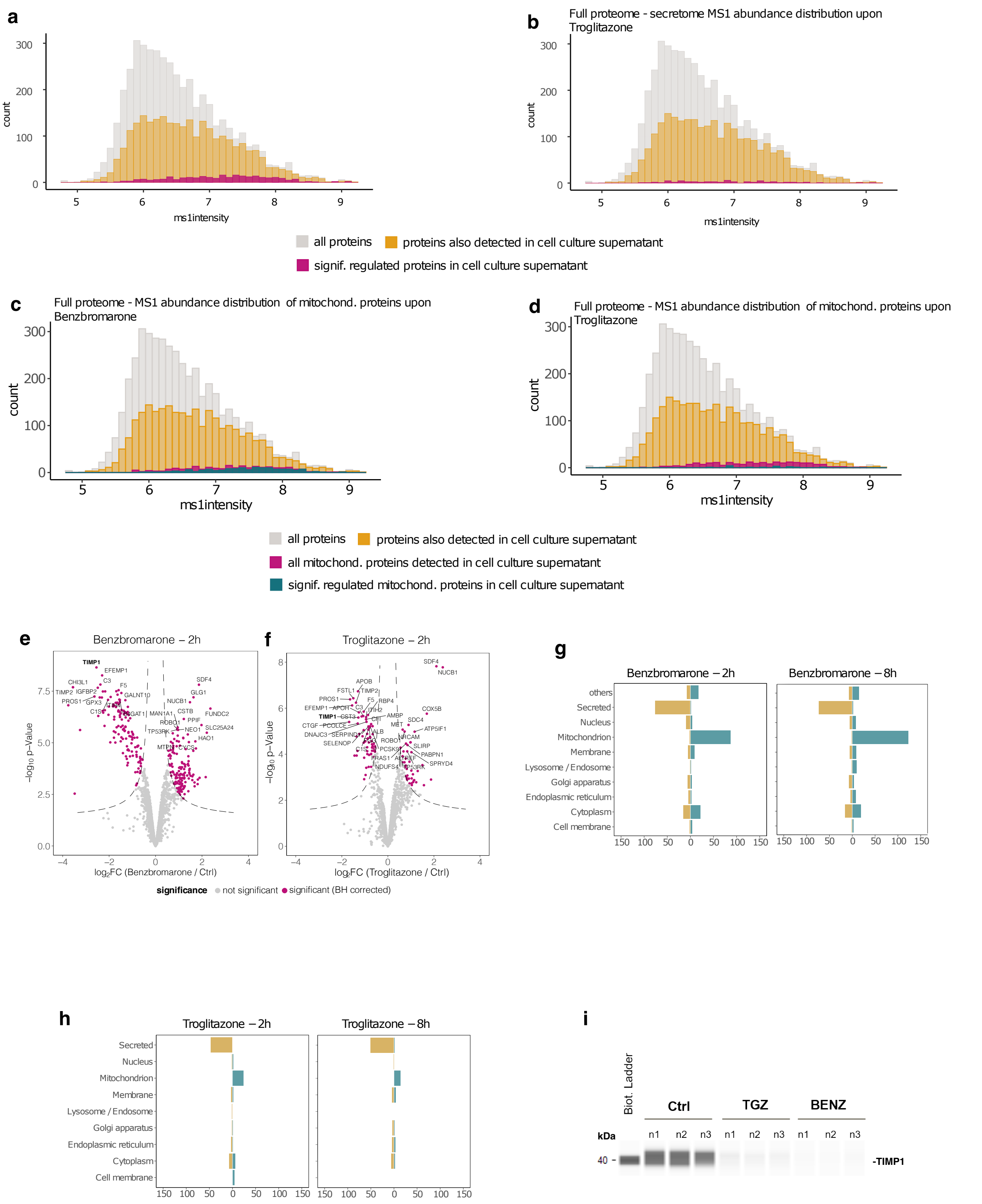


**Supplementary Figure 2 | DILI compounds acutely modulate the secretome of dHepaRG cells.** **a,** MS1 abundance distributions of the dHepaRG full proteome (grey) compared with the MS1 abundance distributions of all proteins detected in the cell culture supernatant (yellow) and the MS1

abundance distributions of all significantly regulated proteins in the cell culture supernatants upon benzbromarone treatment (purple). **b,** same as a for troglitazone.
**c,** MS1 abundance distributions of the dHepaRG full proteome (grey) compared with the MS1 abundance distributions of all proteins (yellow), all mitochondrial proteins (purple) and all significantly regulated mitochondrial proteins (teal) detected in the cell culture supernatants upon benzbromarone treatment. **d,** same as c for troglitazone. **e,** Volcano plot showing proteins quantified in the secretome of benzbromarone treated dHepaRG cells 2h post stimulus. Displayed are the $\log_2$ fold changes and the p-values (-$\log_{10}$-transformed) determined by LIMMA of benzbromarone treated dHepaRG cells (n=3) versus the time matched vehicle (DMSO)- controls (n=3). Dotted line indicates significance cut-offs. Proteins passing the significance thresholds (p(Benjamini Hochberg) < 0.05 and $\log_2$ fold change > 2x standard deviation of the individual treatment are colored in purple. **f,** same as e for 2h troglitazone treatment.
**g,** Bar graph showing protein counts of significantly affected proteins in the secretomes of benzbromarone treated dHepaRG cells grouped by their subcellular location annotation (based on SwissProt). Brown bars depict proteins with reduced secretion, green bars depict proteins with increased secretion. **h,** same as f for troglitazone. **i,** WES analysis of TIMP1 (representative protein with basal secretion in dHepaRG cells) secretion in the supernatants of control-, troglitazone- or benzbromarone treated dHepaRG cells after 8h.

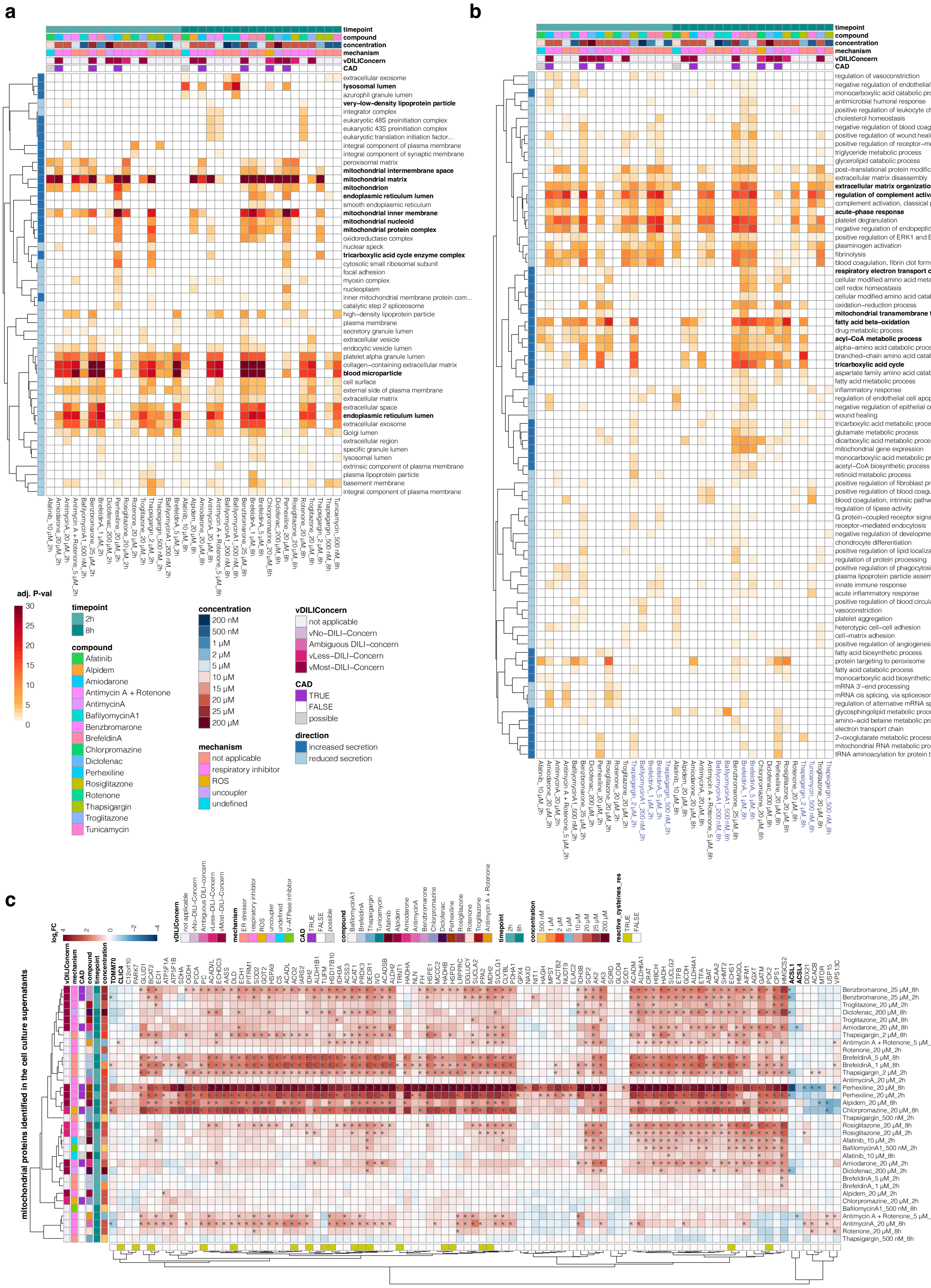


**Supplementary Figure 3 | GO term enrichments of secretomes after treatment with DILI compounds for 2 and 8h.**

**a,** Heatmap displays significant GO-terms (cellular component) derived from the differential secretome analysis of DILI compound treated HepaRG cells (n=3) versus the time matched control at 2h post stimulus (n=3) or 8h post stimulus (n =2). Differentially secreted proteins were determined via LIMMA. Significance thresholds were (p(Benjamini Hochberg) < 0.05 and log2 fold change > 2x standard deviation of the individual treatment. Color intensities indicate the adjusted (BH-corrected) p-value (-log10-transformed) of the GO-term. Only GO-terms are displayed that were significant upon three or more DILI compounds and were an increased secretion could be observed upon compound treatment. Rows are clustered by Pearson correlation. **b,** same as a displaying significant GO-terms (biological process). **c,** Heatmap displaying mitochondrial proteins found in the secretome upon treatment of dHepaRG cells with DILI compounds across different timepoints. Displayed are log2 fold changes to the respective time matched control. Statistically significant changes are denoted with asterisks (*). Row annotation indicates whether proteins harbor reactive cysteine residues (yellow).

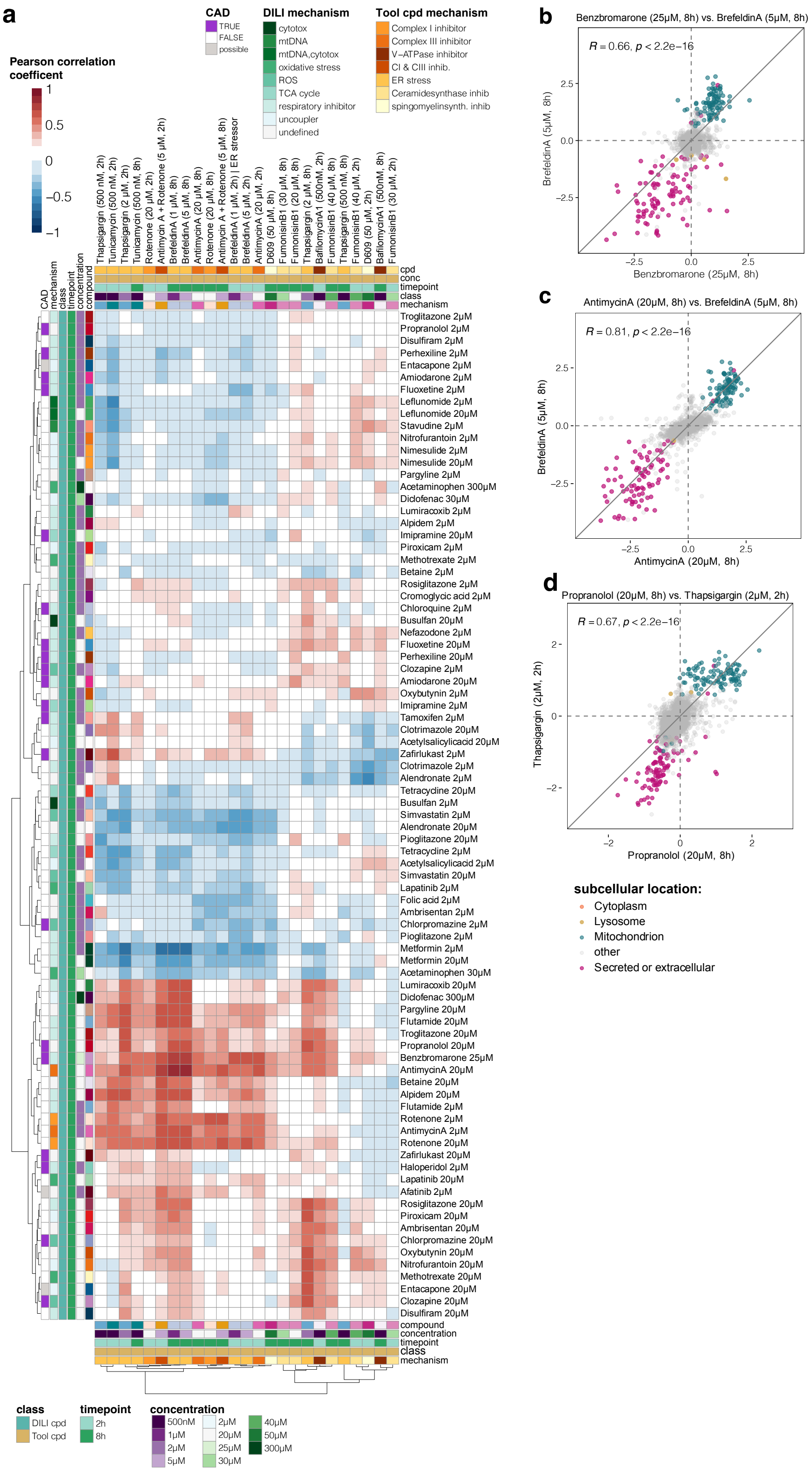


**Supplementary Fig. 4 | Secretion patterns of DILI compounds correlate with secretion changes induced by tool-compounds.**

**a,** Correlation heatmap displaying the pairwise Pearson correlation of DILI compounds (rows) and tool compounds (columns) based on their secretion patterns. Color intensities indicate strength and direction of the correlation, with darker colors indicating stronger correlations. Rows are clustered by Pearson correlation. **b-d,** Pairwise correlation plots of compound induced secretion patterns of: **b,** benzbromarone and brefeldin A, **c,** antimycin A and brefeldin A, **d,** propranolol and thapsigargin. x- and y-axes are represented as $\log_2$ fold change (compound vs. DMSO). Each point represents a protein, color represents the subcellular location based on the UniprotKB annotation: purple: proteins annotated as secreted or extracellular, teal: mitochondrial proteins, yellow: lysosomal proteins, orange: cytoplasmic proteins, grey: proteins with subcellular locations other than dose described. Pearson correlation (R) is given, statistical significance is given by p-value (p).

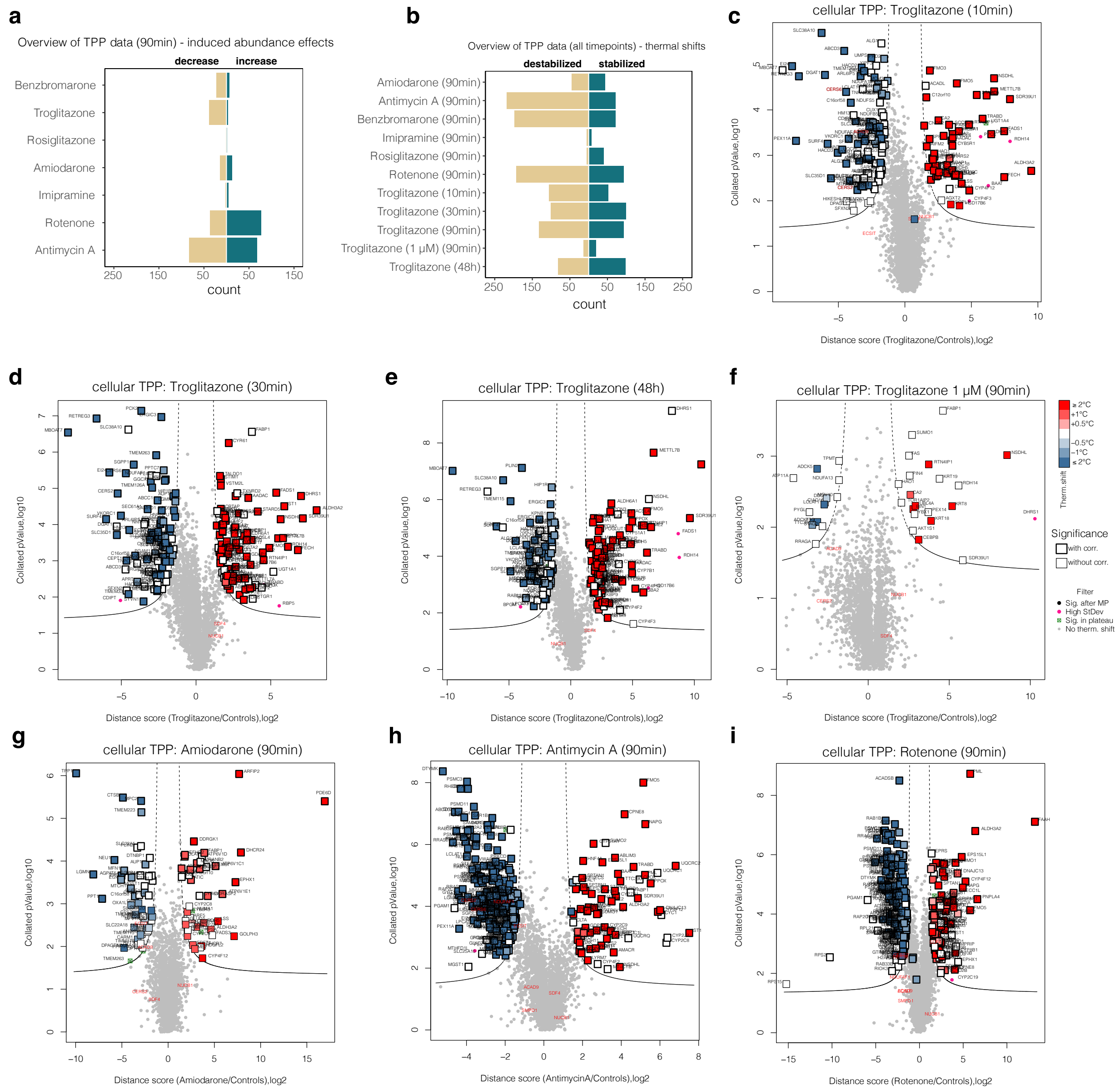

**Supplementary Figure 5 | DILI compounds induce widespread protein thermal stability changes in dHepaRG cells.**

**a,** Overview of significant induced abundance changes after 90 min of compound treatment. Bar graphs show number of proteins with significant abundance increase (green) or decrease (brown) grouped by treatment. **b,** Overview of significant protein thermal stability changes for all treatments. Bar graphs show number of significant proteins stabilized (green) or destabilized (brown) grouped by treatment. **c,** Protein thermal stability changes in dHepaRG cells upon treatment with troglitazone (10 min). Volcano plot displays distance scores and collated P values (-log10 transformed) of proteins quantified in troglitazone (20 μM, n=2) versus DMSO-treated (n=2) dHepaRG cells. The ratio-based approach included LIMMA analysis for ratios between treatment and control groups obtained at each temperature, aggregation of retrieved P values per protein by Brown's method and multiple testing adjustment Benjamini–Hochberg (BH) correction. Dotted line indicates significance cut-offs. Proteins passing significance cut-off are colored according to their Tm shift, bold edges indicate Benjamini-Hochberg corrected P-values with $P < 0.05$, light gray dots depict proteins that were not significantly affected. **d,** same as c for protein thermal stability changes in dHepaRG cells upon treatment with 20 μM troglitazone (30 min). **e,** same as c for protein thermal stability changes in dHepaRG cells upon treatment with 20 μM troglitazone (48 h). **f,** same as c for protein thermal stability changes in dHepaRG cells upon treatment with 1 μM troglitazone (90 min). **g,** same as c for protein thermal stability changes in dHepaRG cells upon treatment with 20 μM amiodarone (90 min). **h,** same as c for protein thermal stability changes in dHepaRG cells upon treatment with 20μM antimycin a (90 min). **i,** same as c for protein thermal stability changes in dHepaRG cells upon treatment with 20 μM rotenone (90 min).

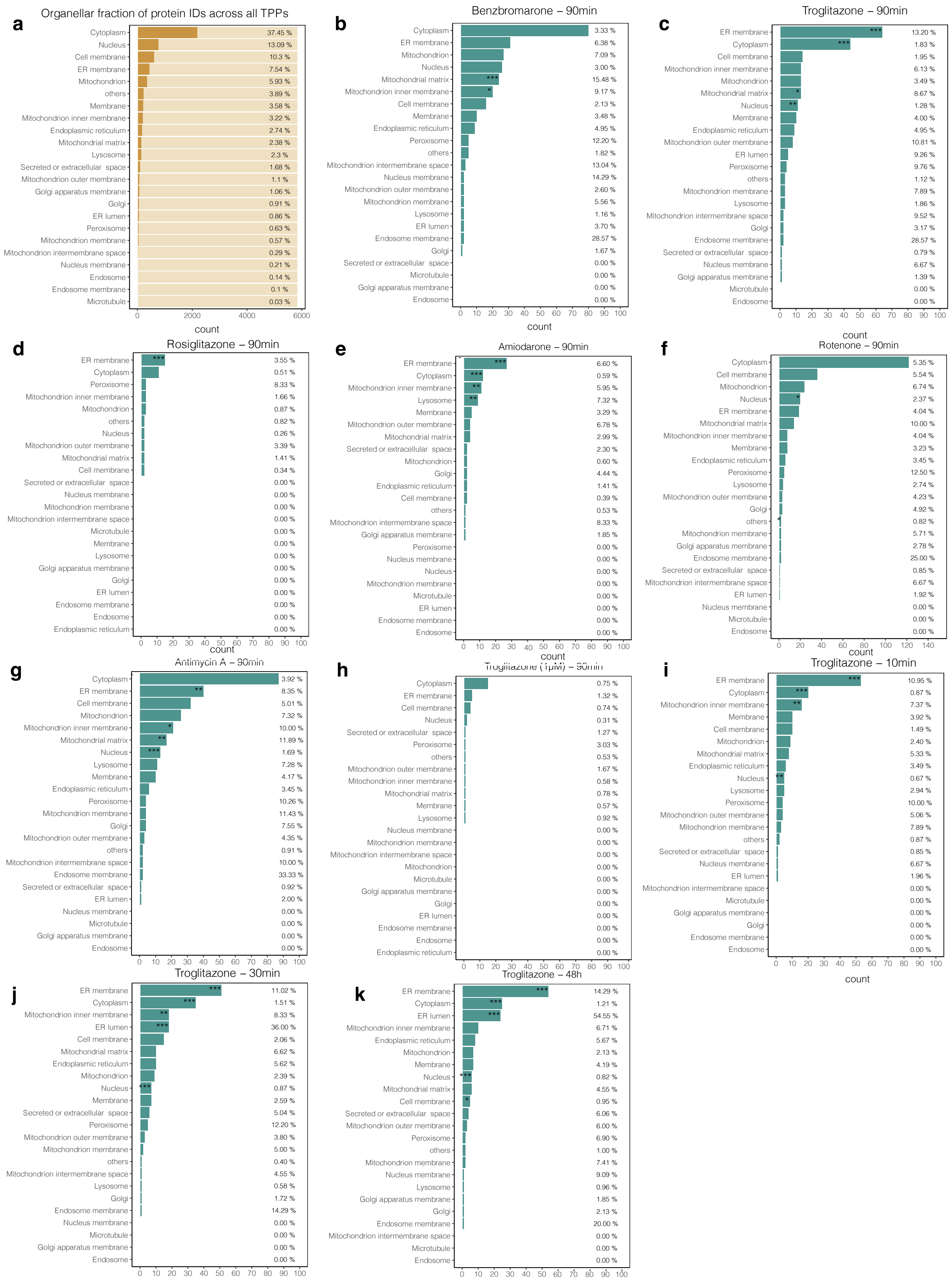

**Supplementary Figure 6 | Thermal proteome profiling confirms organelle specific effects of DILI compounds.**

**a,** Number of proteins identified across all TPP experiments grouped by their subcellular location annotation. Bar graphs show the average number of proteins identified across all TPP experiments per cellular organelle. Length of the dark brown bars represent protein counts, total count of protein identifications across all TPP experiments is represented by the length of the light brown bar, percentage values represent the organellar fraction relative to the average total protein count.
**b,** Total count of proteins with significant protein thermal stability changes upon 90 min treatment with benzbromarone grouped by their subcellular location. Percentage values represent the protein fraction relative to the average protein count of the respective cellular organelle. Asterisks denote significant enrichments for proteins of a respective cellular organelle encoded by: $P \leq 0.05$: *; $P \leq 0.01$: **; $P \leq 0.001$: ***. P-values were calculated with a Fisher-exact test. **c,** same as b for troglitazone (90 min). **d,** same as b for rosiglitazone (90 min). **e,** same as b for amiodarone (90 min). **f,** same as b for rotenone (90 min). **g,** same as b for antimycin A (90 min). **h,** same as b for 1 μM troglitazone (90min). **i,** same as b for troglitazone (10 min). **j,** same as b for troglitazone (30 min).
**k,** same as b for troglitazone (48 h).

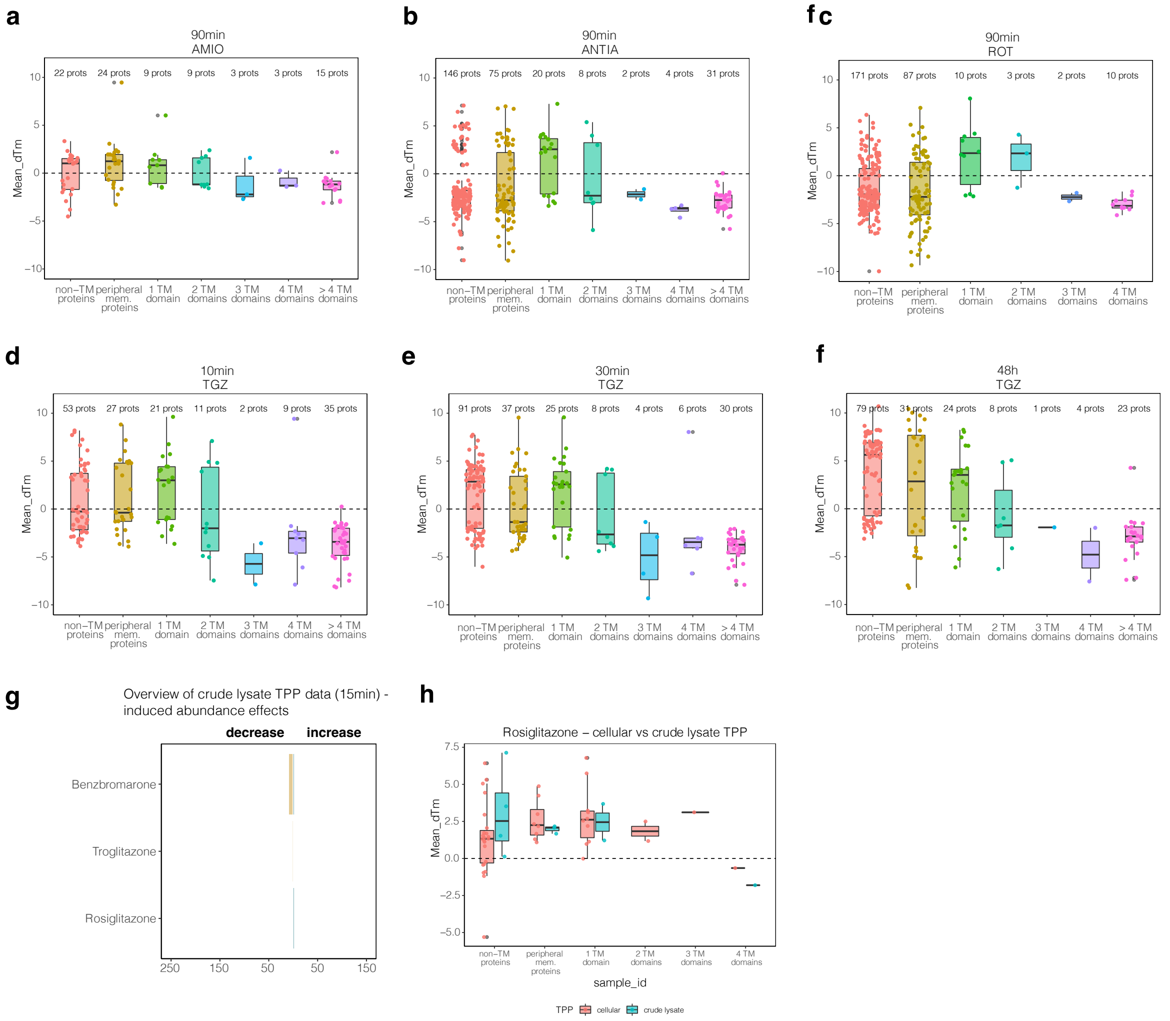


**Supplementary Figure 7 | Effect of DILI compounds on membrane spanning proteins.**

**a,** Comparison of mean dTm (°C) for proteins with significant thermal stability shift > 1 °C or < 1 °C upon amiodarone (90 min) treatment of dHepaRG cells and grouped by their number of transmembrane domains as annotated in Uniprot. Number of proteins per group are indicated. Dotted line denotes a dTm of 0. Center line, median; box limits, upper and lower quartiles; whiskers, maximum and minimum value of the dataset. Points denote the dTm value for each protein in the respective group.

**b,** same as a for antimycin A (90 min). **c,** same as a for rotenone (90 min). **d,** same as a for troglitazone (10 min). **e,** same as a for troglitazone (30 min). **f,** same as a for troglitazone (48 h). **g,** Overview of significant induced abundance changes after 15 min of compound treatment in crude cell lysates. Bar graphs show number of proteins with significant abundance increase (green) or decrease (brown) grouped by treatment. **h,** Comparison of mean dTm (°C) for proteins with significant thermal stability shift from cellular TPP (red) and crude lysate TPP experiments (blue) with rosiglitazone and grouped by their number of transmembrane domains as annotated in Uniprot. Dotted line denotes a dTm of 0. Center line, median; box limits, upper and lower quartiles; whiskers, maximum and minimum value of the dataset. Points denote the dTm value for each protein in the respective group.

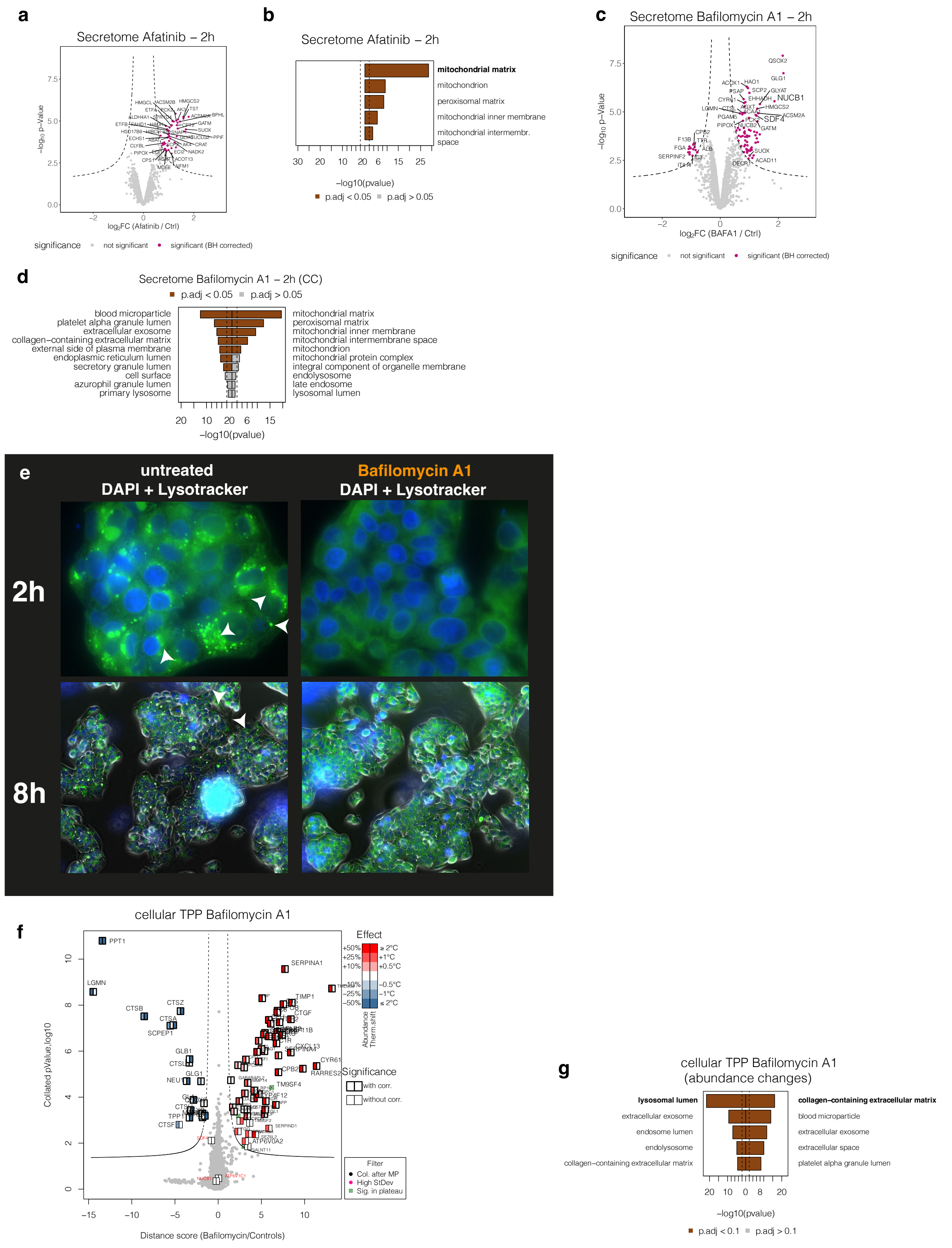


**Supplementary Figure 8 | Secretomics and thermal proteome profiling of cationic amphiphilic drugs (CADs).**

**a,** Changes in protein secretion upon treatment with afatinib 2h post treatment. Volcano plot shows proteins quantified in the secretome of afatinib treated dHepaRG cells 2h post stimulus. Displayed are the log2 fold changes and the pvalues (-log10-transformed) determined by LIMMA of bafilomycin a1 treated dHepaRG cells (n=3) versus the time matched vehicle (DMSO)- controls (n=3). Dotted line indicates significance cut-offs. Proteins passing the significance are colored in purple. **b,** Top five most significantly enriched GO-terms (cellular component) in the secretome of afatinib treated dHepaRG cells (2h). **c,** same as a for treatment with the V-ATPase inhibitor bafilomycin A1 (2h). **d,** same as b for the treatment with V-ATPase inhibitor bafilomycin A1 (2h).
**e,** Bafilomycin A1 leads to a decrease in fluorescence in acidic vesicular structures. Fluorescent analysis of live HepG2 cells, untreated (DMSO only) (left) or bafilomycin A1-treated (right) (500 nM, 2h and 8h), using LysoTracker® Green DND-26 (green). Nuclei were stained with DAPI (blue color). A decrease in fluorescence of acidic vesicular structures stained with Lysotracker Green DND-26 upon treatment with Bafilomycin A1 can be observed. **f,** Protein thermal stability- and abundances changes in dHepaRG cells upon treatment with bafilomycin A1 (90 min). Volcano plot displays distance scores and collated P values (-log10 transformed) of proteins quantified in bafilomycin a1 (20 μM, n=2) versus DMSO-treated (n=2) dHepaRG cells. The ratio-based approach included LIMMA analysis for ratios between treatment and control groups obtained at each temperature, aggregation of retrieved P values per protein by Brown's method and multiple testing adjustment Benjamini–Hochberg (BH) correction. Dotted line indicates significance cut-offs. Proteins passing significance cut-off are colored according to their Tm shift, bold edges indicate Benjamini-Hochberg corrected P-values with $P < 0.05$, light gray dots depict proteins that were not significantly affected. **g,** Top five most significantly enriched GO-terms (cellular component) in the TPP of bafilomycin A1 treated dHepaRG cells. GO enrichment was tested on proteins with abundance change using a Fisher` s exact test. Significant GO-terms (P(BH corrected) < 0.05) are depicted as brown bars.

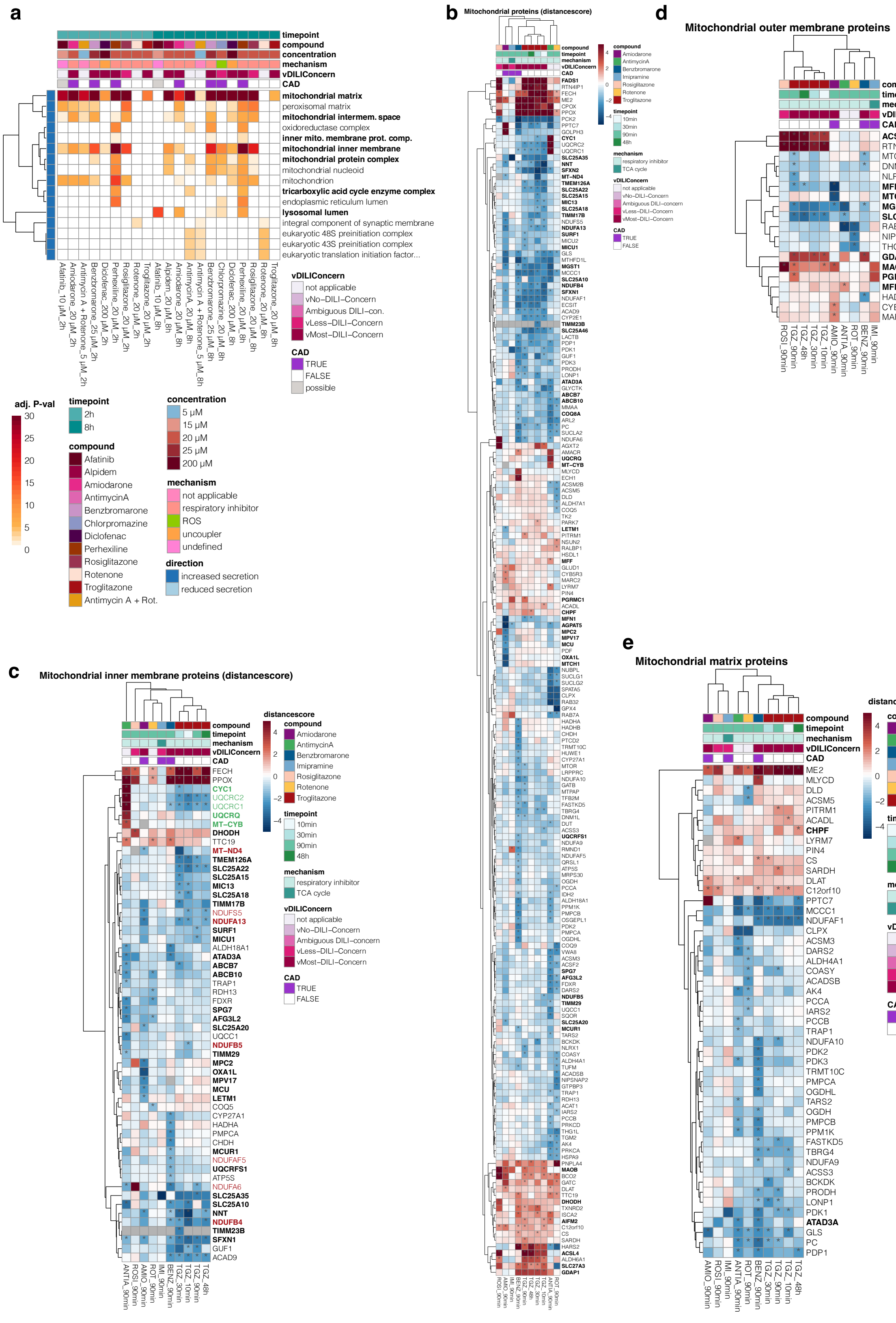


**Supplementary Figure 9 | Drug-induced toxicity on mitochondria.**

**a,** DILI compounds induce an early secretion of mitochondrial proteins. Heatmap displays significant GO-terms (cellular component) derived from the differential secretome analysis of DILI compound treated HepaRG cells (n=3) versus the time-matched control at 2h post stimulus (n=3) or 8h post stimulus (n =2). Colors indicate the adjusted (BH-corrected) p-value (-log10-transformed) of the GO-term. Only GO-terms are displayed that were significant upon three or more DILI compounds and were an increased secretion could be observed upon compound treatment. Rows were clustered by Pearson correlation. **b,** TPP identifies proteins mitochondria affected by DILI compounds. Heatmap displays all proteins of mitochondria with significant thermal stability shifts upon treatment of HepaRG cells with DILI compounds. Displayed are distance scores. Rows and columns are clustered by Euclidian distance. Statistically significant changes are denoted with asterisks (*). Bold gene names denote transmembrane proteins.**c,** same as b for proteins of the inner mitochondrial membrane. Gene names in red denote subunits of MRC complex I, gene names in green denote subunits of MRC complex III. **d,** same as b for proteins of the outer mitochondrial membrane.**e,** same as b for mitochondrial matrix proteins.

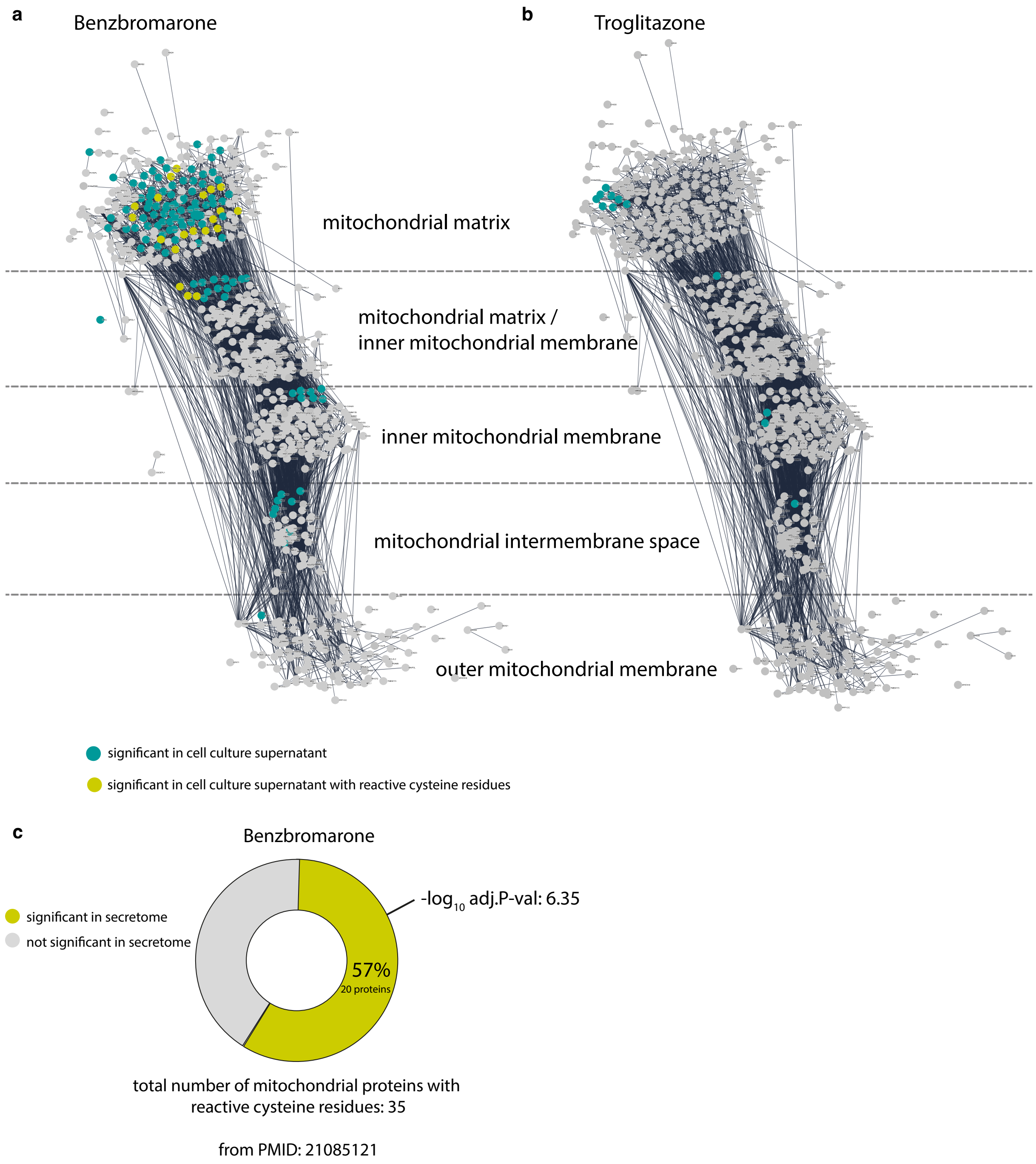


**Supplementary Figure 10 | Benzbromarone and Troglitazone induce the secretion of a subnetwork of mitochondrial proteins into the cell culture supernatant.**

**a,** STRING analysis of mitochondrial proteins identified in the full proteome analysis of dHepaRG cells. Proteins are clustered according to their mitochondrial location. Proteins in teal were found to be significantly released into the cell culture supernatant upon benzbromarone treatment (8h timepoint). Proteins in yellow were found to be significantly released into the cell culture supernatant upon benzbromarone and harbor reactive cysteine residues. **b,** same as a for troglitazone. **c,** a proteomics dataset[144] was used to identify mitochondrial proteins harboring reactive cysteine residues susceptible to oxidative stress upon benzbromarone treatment. A hypergeometric test was used to calculate a p-value for the enrichment.

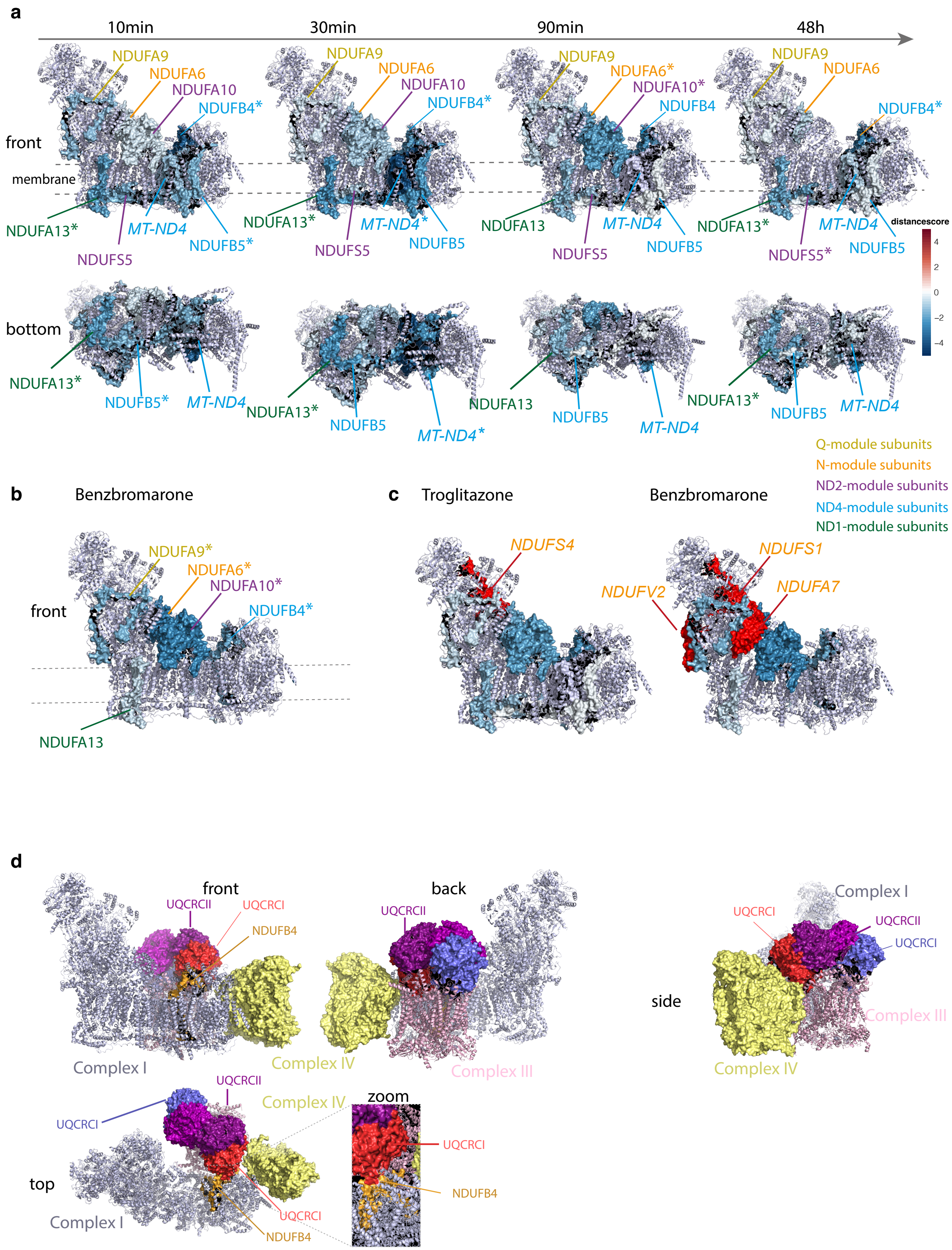


**Supplementary Figure 11 | DILI compounds that induce mitochondrial secretion acutely destabilize proteins of the respiratory complexes I and III .**

**a,** Time-dependent TPP experiments with troglitazone reveal an initial thermal destabilization of membrane-spanning and proton-transporting ND4- and ND2 module subunits, followed by destabilization of CI matrix arm subunits at later timepoints. Molecular visualization of MRC complex I (structure was derived from PDB database, identifier: 5XTH). Surface representations indicate thermal destabilized subunits after 10 min, 30 min, 90 min and 48h of compound treatment with the MRC inhibitor troglitazone. Color indicates the distancescore. Dotted lines represent the inner mitochondrial membrane. Statistically significant thermal stability changes are denoted with asterisks (*).
**b,** same as a for 90 min treatment with benzbromarone. **c,** same as a for troglitazone and benzbromarone after 90 min of compound treatment. Surface representations in red indicate complex I matrix arm subunits with significantly increased abundances in cell culture supernatants after compound treatment.
**d,** Molecular visualization of the human respiratory supercomplex I1III2IV1 (structure was derived from PDB database, identifier: 5XTH). Surface representations indicate CI, CIII and CIV subunits of interest. Zoom shows interaction-site of CI subunit NDUFB4 with the CIII subunit UQCRCI.

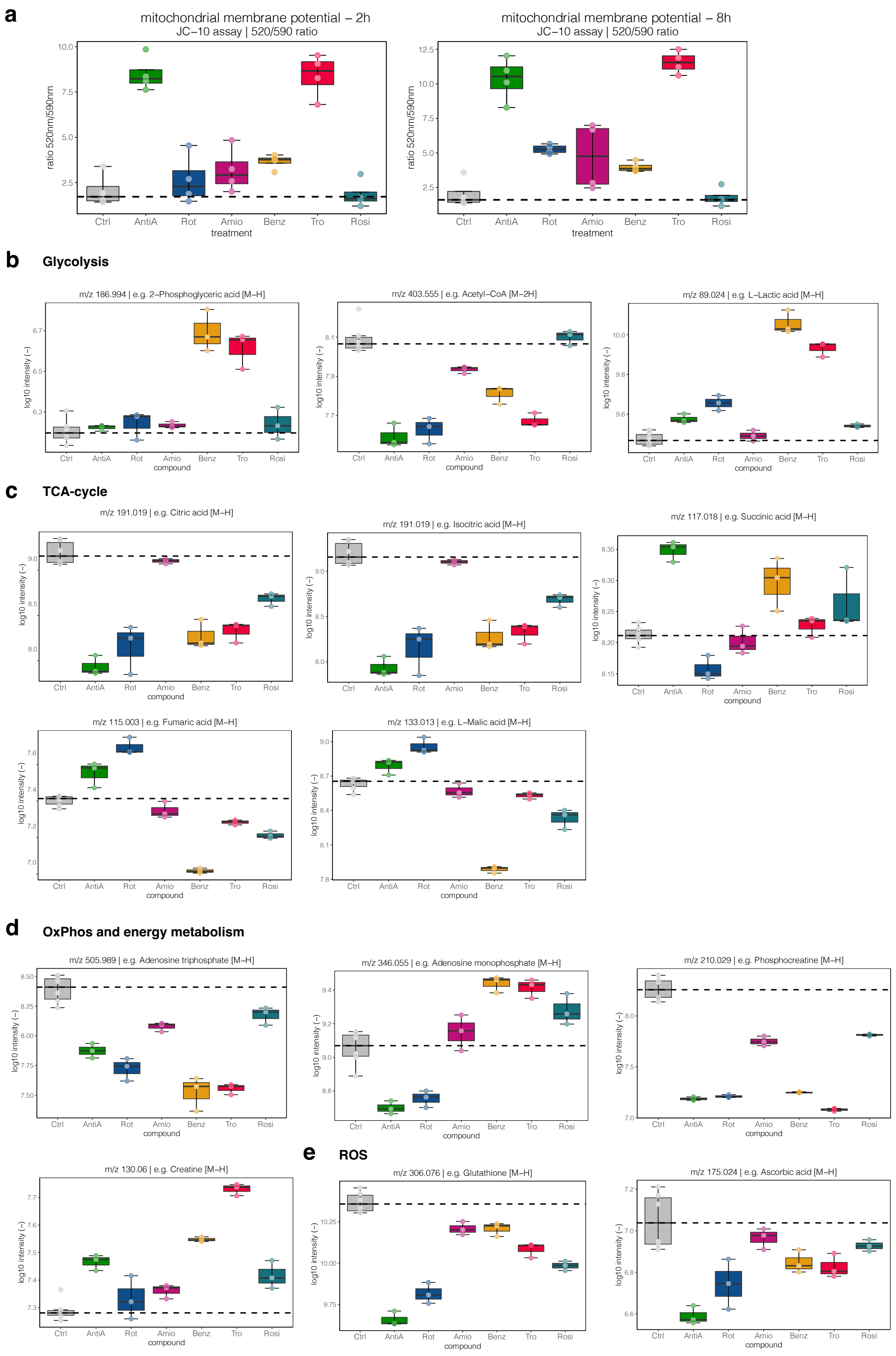


**Supplementary Figure 12 | Metabolic perturbations upon DILI compound treatment confirm an acute MRC inhibition.**

**a,** Boxplots display changes of the mitochondrial membrane potential as ratios of the fluorescence measurement at 520 and 590 nm determined with an JC-10 assay after 2h (left panel) and 8h (right panel) of compound treatment. Points denote the 520/590nm ratios derived from 4 independent biological replicates. **b,** Boxplots depict MS1 intensities (-$log_{10}$ transformed) of characteristic glycolytic metabolites after 8h of compound treatment. Data points denote MS1 intensities derived from each of the three independent biological replicates. **c,** same as b for TCA-cycle metabolites. **d,** same as b for metabolites involved in the oxidative phosphorylation and energy metabolism. **e,** same as b for metabolites involved in the redox homeostasis and ROS scavenging.

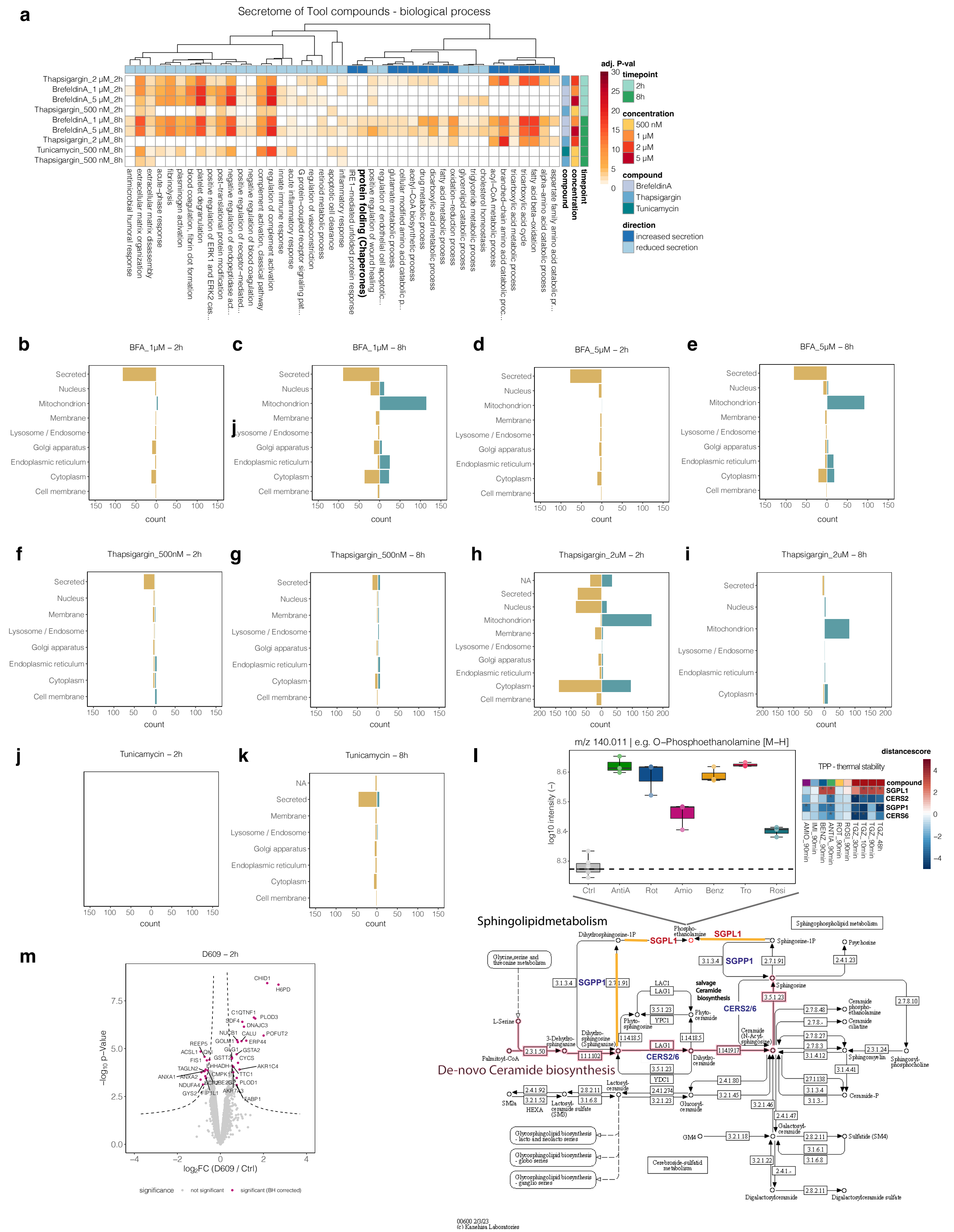


**Supplementary Figure 13 | ER stressors mimic DILI-compound induced secretion events.**

**a,** Heatmap displays significant GO-terms (biological process) derived from the differential secretome analysis upon treatment of HepaRG cells with different ER-stressors (n=3) versus the time-matched control at 2h post stimulus (n=3) or 8h post stimulus (n =2). Differentially secreted proteins were determined via LIMMA. Significance thresholds were (p(Benjamini Hochberg) < 0.05 and log2 fold change > 2x standard deviation of the individual treatment. Color intensities indicate the adjusted (BH-corrected) p-value (-log10-transformed) of the GO-term. Only GO-terms are displayed that were significant upon three or more DILI compounds and were a reduced secretion could be observed upon compound treatment. Rows are clustered by Pearson correlation. **b,** Bar graph shows proteins significantly changing in the secretomes of brefeldin A (1 µM) treated dHepaRG cells (2h) grouped by their subcellular location annotation (based on UniProt annotation). Brown bars indicate significantly lower secreted proteins, green bars indicate a significant upregulation in protein secretion. Displayed are counts for each subcellular location. **c,** same as b for proteins significantly changing in the secretomes of brefeldin A (1 µM) treated dHepaRG cells (8h). **d,** same as b for proteins significantly changing in the secretomes of brefeldin A (5 µM) treated dHepaRG cells (2h). **e,** same as b for proteins significantly changing in the secretomes of brefeldin A (5 µM) treated dHepaRG cells (8h). **f,** same as b for proteins significantly changing in the secretomes of thapsigargin (500 nM) treated dHepaRG cells (2h). **g,** same as b for proteins significantly changing in the secretomes of thapsigargin (500 nM) treated dHepaRG cells (8h). **h,** same as b for proteins significantly changing in the secretomes of thapsigargin (2 µM) treated dHepaRG cells (2h).**i,** same as b for proteins significantly changing in the secretomes of thapsigargin (2 µM) treated dHepaRG cells (8h). **j,** same as b for proteins significantly changing in the secretomes of tunicamycin treated dHepaRG cells (2h). **k,** same as b for proteins significantly changing in the secretomes of tunicamycin treated dHepaRG cells (8h). **l,** drug induced impairment of ceramide metabolism leads to increased abundance of phosphoethanolamine. Boxplots depict MS1 intensities (-log10 transformed) of phoshphoethanolamine after 8h of compound treatment. Center line of each box, median; box limits, upper and lower quartiles; whiskers, maximum and minimum value of the dataset. Dotted line, median of the control samples. Data points denote MS1 intensities derived from each of the three independent biological replicates. Heatmap displays proteins of the ceramide and sphingolipid metabolism exhibiting a thermal stability shift upon treatment of dHepaRG cells with DILI compounds for 90min irrespective of their significance. Displayed are distancescores. Statistically significant changes are denoted with asterisks (*). Pathway map was derived from KEGG. The de-novo ceramide biosynthetic pathway is colored in pink. Upon inhibition of cellular ceramide synthases, accumulating dihydrosphinganine is stepwise converted (yellow pathway) to phosphoethanolamine.

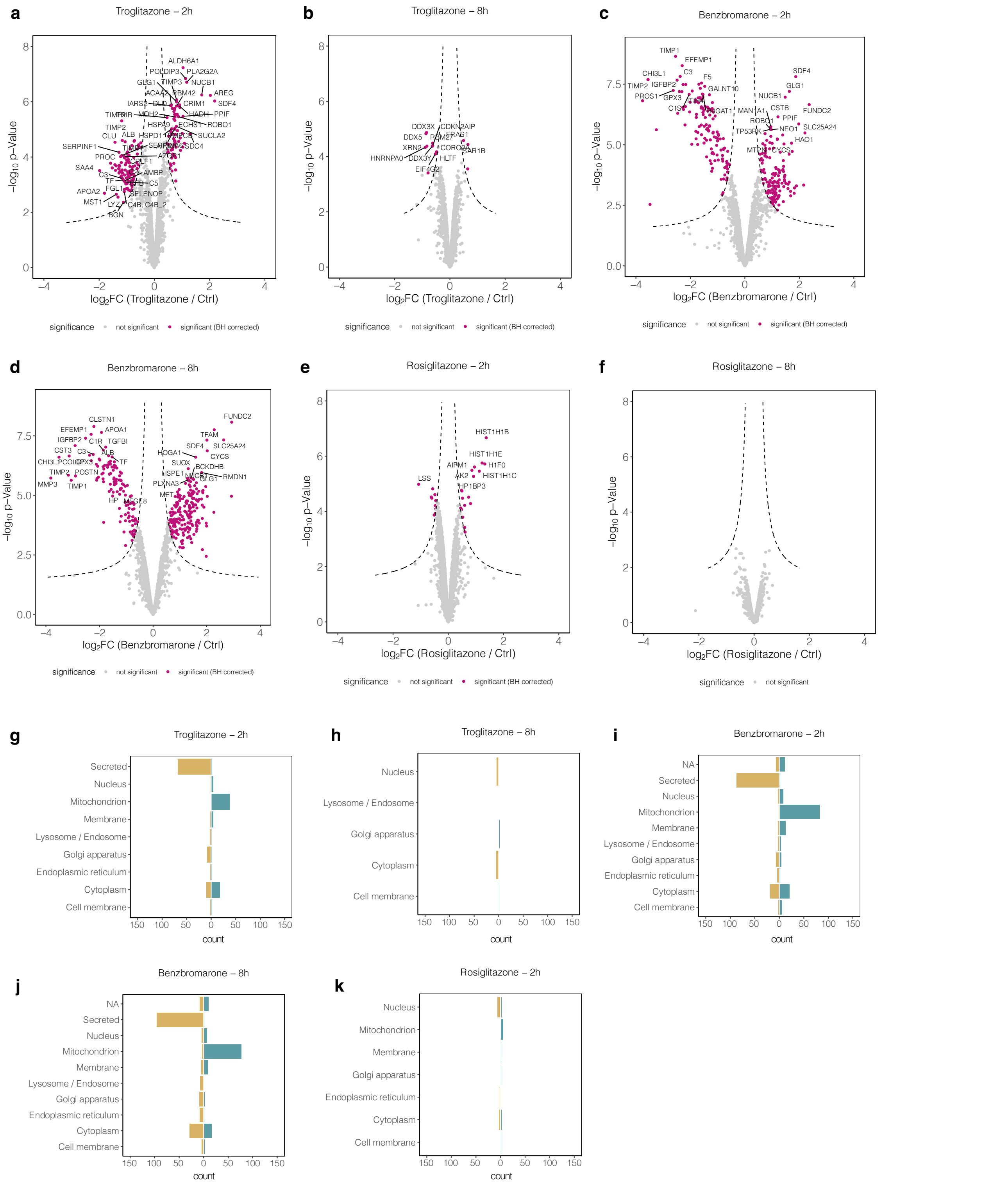


**Supplementary Figure 14 | Metabolically inactive HepG2 cells induce similar secretion patterns upon troglitazone, benzbromarone and rosiglitazone treatments.**

**a,** Volcano plot shows proteins quantified in the secretome of troglitazone (20 μM) treated HepG2 cells 2h post stimulus. Displayed are the log2 fold changes and the p-values ($-\log_{10}$-transformed) determined by LIMMA of benzbromarone treated HepG2 cells (n=3) versus the time matched vehicle (DMSO)-controls (n=3). Dotted line indicates significance cut-offs. Proteins passing the significance thresholds are colored in purple. **b,** same as a for 8h troglitazone (20 μM) treatment. **c** and d same as a and b for treatment with benzbromarone (25 μM). **e** and f, same as a and b for treatment with rosiglitazone (20 μM). **g** and h, Bar graphs displaying protein counts of significantly affected proteins in the secretomes of troglitazone (2h and 8h) treated HepG2 cells grouped by their subcellular location annotation (based on SwissProt). Brown bars depict proteins with reduced secretion, green bars depict proteins with increased secretion. **I** and **j**, same as g and h for benzbromarone. **k**, same as a for rosiglitazone
